# Characterization, modelling and mitigation of gene expression burden in mammalian cells

**DOI:** 10.1101/867549

**Authors:** T Frei, F Cella, F Tedeschi, J Gutierrez, GB Stan, M Khammash, V Siciliano

**Affiliations:** Department of Biosystems Science and Engineering, ETH Zürich; Istituto Italiano di Tecnologia-IIT, Largo Barsanti e Matteucci, Naples ITA; University of Genoa, Genoa ITA; Department of Bioengineering and Centre for Synthetic Biology, Imperial College London, U.K

## Abstract

Despite recent advances in genome engineering, the design of genetic circuits in mammalian cells is still painstakingly slow and fraught with inexplicable failures. Here we demonstrate that competition for limited transcriptional and translational resources dynamically couples otherwise independent co-expressed exogenous genes, leading to diminished performance and contributing to the divergence between intended and actual function. We also show that the expression of endogenous genes is likewise impacted when genetic payloads are expressed in the host cells. Guided by a resource-aware mathematical model and our experimental finding that post-transcriptional regulators have a large capacity for resource redistribution, we identify and engineer natural and synthetic miRNA-based incoherent feedforward loop (iFFL) circuits that mitigate gene expression burden. The implementation of these circuits features the novel use of endogenous miRNAs as integral components of the engineered iFFL device, a versatile hybrid design that allows burden mitigation to be achieved across different cell-lines with minimal resource requirements. This study establishes the foundations for context-aware prediction and improvement of *in vivo* synthetic circuit performance, paving the way towards more rational synthetic construct design in mammalian cells.

## Introduction

Mammalian synthetic biology facilitates the study of diverse biological processes including gene regulation^1^, developmental patterns^2^, and cancer progression^3^. More recently, it has gained clinical relevance, offering powerful new tools for the engineering of recombinant protein-producing cells^4^ and for the creation of novel cell-based therapies for clinical use^5–7^. These critical applications require synthetic gene circuits that are robust and predictable with minimal impact on their host cells. However, engineering such circuits requires numerous design-build-test-learn iterations^8,9^, which are particularly expensive and time consuming^10^ in mammalian cells.

At the core of the problem is the poor predictability of gene expression^9^ in engineered cells arising from the dependence of gene expression on the cellular context. In particular, the often overlooked dependence of exogenous genetic circuits on limited host resources that are shared with endogenous pathways frequently leads to unanticipated and counterintuitive circuit behaviors^11^. In bacterial cells, substantial progress towards increasing the predictability of gene expression has been made by showing that exogenous genetic material imposes a significant burden, resulting in decreased growth rates and degraded cellular performance^12^. This has been attributed to the diversion of the pool of resources available for gene expression^13,14^ towards transcription and translation of the newly introduced synthetic payloads. These observations prompted the development of models that consider gene expression in a resource-limited context^15–18^ and led to approaches for mitigating the impact of resource burden in bacteria^19,20^. Analogous studies in *S. Cerevisiae* showed that transcription and translation are limiting processes^21^. In mammalian cells, while performance shortcomings of synthetic circuits due to transactivator dosage and plasmid uptake variation^22^ have been observed, a deeper understanding of the problem of resource burden and methods for its mitigation are still missing. Moreover, competition for endogenous resources might have detrimental effects on basic and translational biology. For instance, in studies based on transient DNA expression, genes that are used to normalize the results might be subject to resource-dependent expression coupling (e.g protein levels measured by flow cytometry are usually normalized to the expression levels of the transfection marker, which is also used as a measure of transfection efficiency).

Here, we investigate the burden imposed by synthetic circuits on host cells (Fig. 1). Through the design of genetic constructs that allow us to uncouple transcription and translation processes, we separately study transcriptional and translational burden caused by cellular resource sharing. In particular, we engineer several regulatory circuits composed of a tunable load, called *X-tra* (eXtra Transgene), which we genetically express in the host cell in varying amounts. We then measure the impact of this tunable load on a ‘sensor’ gene, which we refer to as the *capacity monitor* (Fig. 1a). We demonstrate in different mammalian cell lines that the sharing of transcriptional and translational resources in the host cell can tightly couple otherwise independently co-expressed synthetic genes and lead to trade-offs in their expression (Fig. 1a). To enhance the predictability of synthetic devices in mammalian cells, we explicitly incorporate these load sharing effects in a general mathematical model in which we replace the rates of resource-dependent reactions with adjusted *effective* rates (Fig. 1b). We demonstrate that such a resource-aware model successfully recapitulates the observed, and sometimes unexpected, dose-responses of the various genetic circuits we tested. Additionally, we investigate the role of post-transcriptional regulators, like RNA-binding proteins (RBPs) and microRNAs (miRNAs) in mitigating the impact of burden-induced coupling and find that both are able to reallocate resources, making them candidates for use in burden-mitigation circuits. Using these observations, and guided by our modelling framework, we identify the *incoherent Feed-Forward Loop* (iFFL) as a network topology that is particularly effective at resource burden mitigation, and then we use endogenous and synthetic microRNA (miRNA) regulation to engineer iFFL-based, burden-mitigating synthetic circuits (Fig. 1c). While miRNA-based iFFL circuits have been previously constructed to buffer gene expression against noise^23^ and fluctuations in external inducer concentration^24,25^, in this study we demonstrate for the first time that they also act to rescue the expression level of genes of interest despite changes in available cellular resources due to the loading effects of transgene constructs (Fig. 1c).

**Fig. 1.**
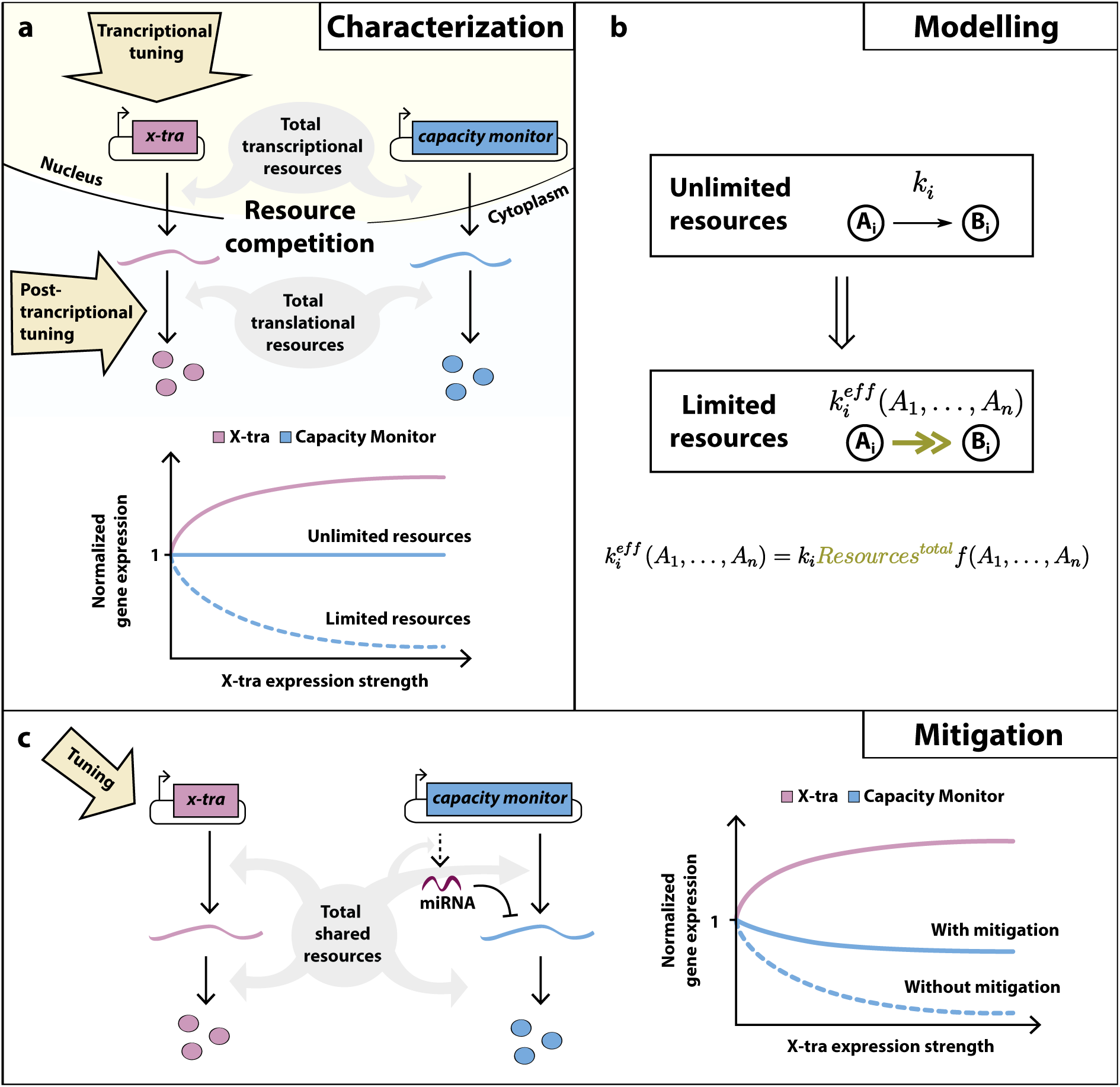
Graphical abstract. **(a) Characterization of gene expression burden.** Expression of independent exogenous genes impacts on host cellular resources. Thus, perturbations in one gene’s expression (hereby named *X-tra*) affect the expression of a second gene (hereby named *capacity monitor*). **(b) Modelling of gene expression in a resource limited environment.** Modelling of gene expression is generally performed under the assumption of unlimited resources. A simple framework enables the straightforward transformation of such a model to a system that incorporates resources explicitly. The transformation involves a simple function that scales the original reaction rate. **(c) Mitigation of gene expression burden.** A simple microRNA-based circuit motif is capable of mitigating the burden-induced coupling of *X-tra* and the *capacity monitor*.

To the best of our knowledge this is the first study to characterize the individual transcriptional and translational burden contributions induced by exogenous gene expression in mammalian cells, and to unveil mechanisms of resource reallocation for burden mitigation using post-transcriptional regulation. The study also pioneers the use of *endogenous* miRNAs as integral parts of synthetic circuits, which in our case consist of iFFL burden-mitigation devices. Together with a refined mathematical modelling framework that describes biochemical reactions in the context of finite resources, these findings pave the way to more realistic output predictions and optimal synthetic construct design in mammalian cells.

## Results

### Genetic circuits impose a burden on mammalian cells as they compete for limited shared cellular resources

We reasoned that due to competition for finite cellular resources, there is a threshold above which gene expression from synthetic circuits saturates. To test this, we co-transfected HEK293T cells with two constitutively expressed fluorescent proteins mCitrine and mRuby3 driven by EF1α promoters, in molar ratios ranging from 1:4 to 4:1, for a total of 50 ng (low) or 500 ng (high) of encoding plasmid (Fig. 2a). Model-based computer simulations of the expected response in the presence of competition for limited resources are presented (Fig. 2a, **left**). The equations used to generate these plots were derived from the modelling framework introduced later on in Fig. 4a (**Supplementary Note 1**). In accordance with the simulations, the total amount of 500 ng of encoding plasmids results in a dramatic drop in experimentally measured global gene expression as compared to 50 ng (Fig. 2a, **right**). Furthermore, in both experimental conditions mCitrine and mRuby3 fluorescence levels are negatively correlated; the higher the amount of expressed mCitrine, the lower that of mRuby3 and vice versa (Fig. 2a, **right**); this correlation was also more severe for 500 ng of transfected plasmid than for 50 ng.

**Fig. 2.**
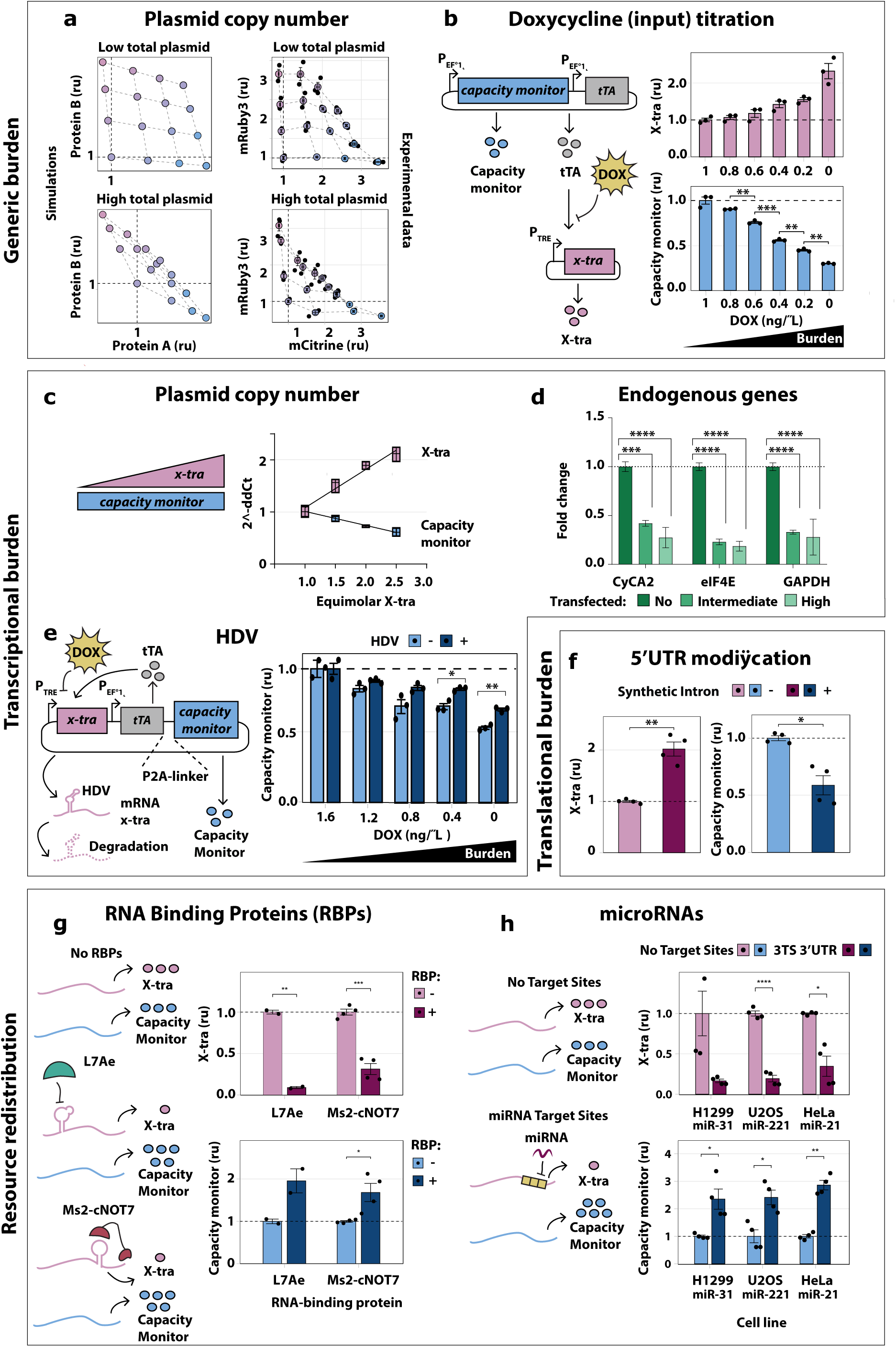
Burden imposed by genetic circuits in mammalian cells. **(a)** Left: model-guided simulations accounting for the limited resources (**Supplementary Note 1**) shows the correlation of independent genes’ expression. As the total plasmid amount increases, the total protein expression plateaus. Right: Titration of two plasmids constitutively expressing the fluorescent proteins mCitrine and mRuby3 from EF1α promoters in ratios from 1:4 to 4:1, for a total of 50 ng (Top right) or 500 ng of DNA (Bottom right). Computer simulations effectively recapitulate the saturation of gene expression that we observed for a total amount of 50 or 500 ng of plasmids. N=3 biological replicates. **(b)** Inhibition of tTA by Dox treatment further unveils the hidden coupling of expression levels among two genes. Two plasmids were co-transfected, one constitutively expressing *capacity monitor* and tTA from a strong constitutive promoter and the other expressing *X-tra* from a tTA responsive promoter. Data show that *capacity monitor* levels counterbalance the increase in *X-tra* expression. Flow cytometry data are normalized to the expression levels of each gene in absence of Dox. N=3 biological replicates. **(c)** Transcriptional burden. mRNA quantification of *X-tra* and a *capacity monitor* expressed at different molar ratios. As the *X-tra* increases, the mRNA levels of the *capacity monitor* decreases. N=4 biological replicates. qPCR analysis was performed 48 hours post transfection and data show fold change +/− SE. **(d)** Cells transfected with a plasmid encoding two fluorescent proteins expressed from a bidirectional promoter were sorted according to *high*, *intermediate* or *no* fluorescence **(Supplementary figure 8)**, and the mRNA was extracted. qPCR shows that the mRNA levels expressed from endogenous genes decrease in cells with *intermediate* and *high* fluorescence. N=3 biological replicates. N=3 biological replicates. Data show fold change +/− SE. **(e)** Increasing the expression of a translationally incompetent mRNA suggest transcriptional burden. *Capacity monitor* levels are higher in the presence of an active HDV ribozyme placed after the start codon of *X-tra* (which results in its rapid degradation of *X-tra*) than in the inactive form, suggesting a sequestration of transcriptional resources. N=3 biological replicates. **(f)** Translational burden. We included a synthetic intron in the 5’UTR of a transcript to increase its translation and to selectively observe translational burden. As expected the synthetic intron shows higher translational load of *X-tra* as compared to the control and leads to reduced *capacity monitor* levels. N=4 biological replicates. **(g)** Post-transcriptional regulators impact translational burden. The RBPs L7Ae and Ms2-cNOT7, downregulate *X-tra* expression by binding the 5’ and 3'UTR respectively. Data show that repressed *X-tra* expression leads to increased *capacity monitor* levels. N≥2 biological replicates. **(h)** Similar to (f) also miRNAs reallocate resources by downregulating targeted mRNA which include miRNA target sites (TS) in the 3’ UTR. We engineered *X-tra* transcripts with 3TS that respond to either miRNA miR-31, miR-221 or miR-21, endogenously expressed in H1299, U2OS and HeLa cells respectively. Data show that *capacity monitor* levels are higher when the *X-tra* is downregulated by miRNAs. N≥3 biological replicates. Flow cytometry data were acquired 48 hours post-transfection and is plotted +/− SE. SE: standard error. ru: relative units. unpaired T-test. p-value: ****<0.0001, ***<0.0005, **<0.005, *<0.05

**Fig. 3.**
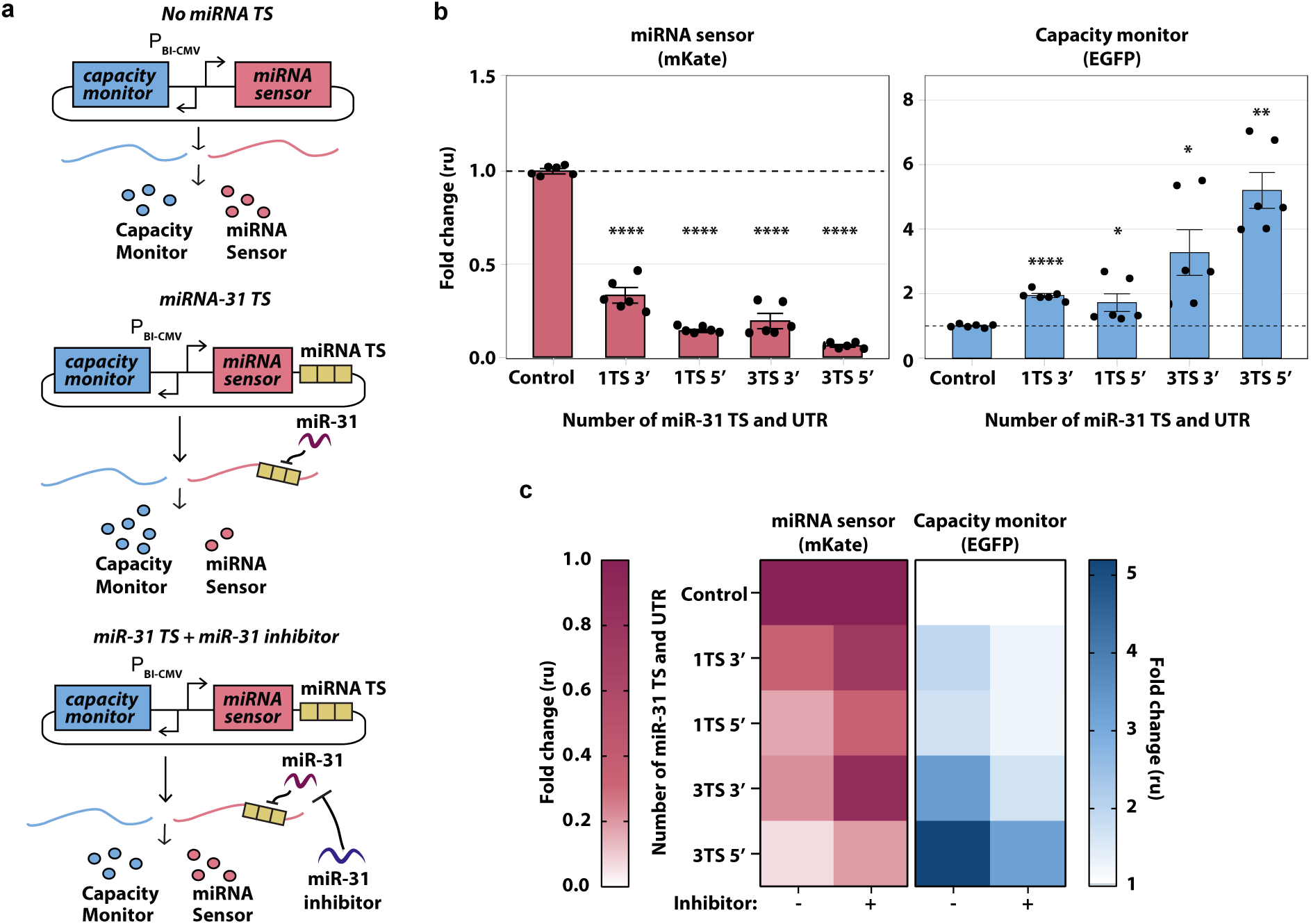
Impact of miRNA target sites number and location on burden. **(a)** Schematics of experimental design to infer miRNA-mediated cellular resources redistribution. EGFP (*capacity monitor*) and mKate (*miRNA sensor*) are encoded on the same bi-directional CMV promoter plasmid. 1 or 3 TS for miR-31 (TS) are added either in the 3’ or 5’UTR of mKate. Control: no miR31 TS. Hypothesis: in the absence of miR-31 regulation, *capacity monitor* and *miRNA sensor* are expressed to a certain level (left). In the presence of miR-31, lower *miRNA sensor* levels correlate with higher *capacity monitor* expression (middle). This condition is reversed by a miR-31 inhibitor (right). **(b)** Fold change of *miRNA sensor* and *capacity monitor* protein levels compared to control (set to 1). EGFP increases up to 5 fold with the strongest downregulation of mKate (3TS 5’UTR). Flow cytometry data were acquired 48 hours post-transfection and are plotted +/− SE. SE:standard error. ru: relative units. N=6 biological replicates, unpaired T-test. p-value: ****<0.0001, ***<0.0005, **<0.005, *<0.05. **(c)** When miR31 activity was impaired by a miR-31 inhibitor, the rescue of mKate expression corresponds to reduced EGFP levels, whereas both fluorescent proteins do not vary in the control. Flow cytometry data was acquired 48 hours post-transfection and is plotted as a heatmap of the fold change. The fold change is calculated as the ratio between the mean of 6 biological replicates and the corresponding mean in the control condition. Corresponding bar plots and statistical analysis are reported in **Supplementary Fig. 15**.

**Fig. 4.**
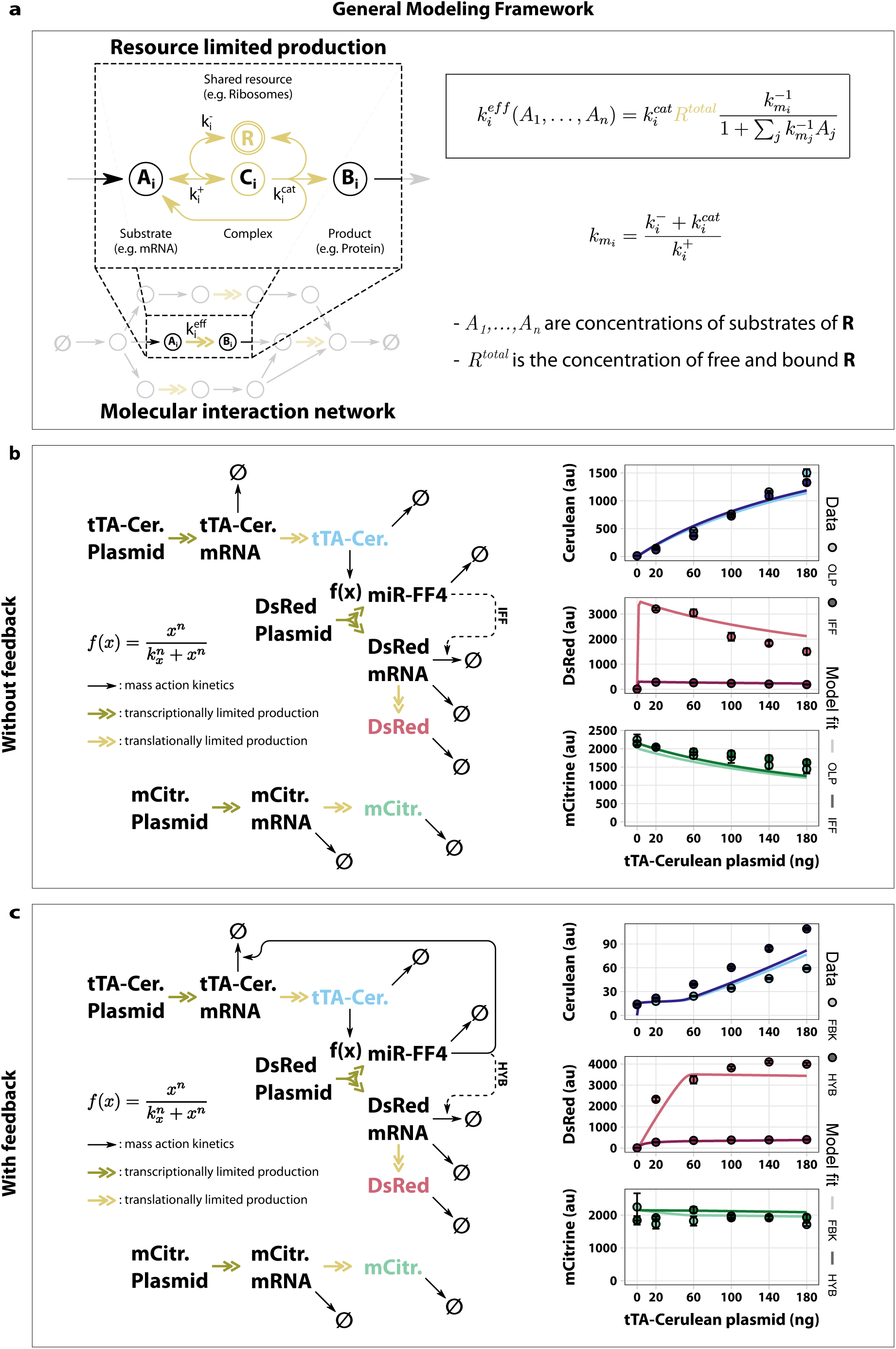
A mathematical network model that factors in burden-induced coupling reproduces counter-intuitive experimental observations. **(a) General framework for transforming molecular interaction network models.** Existing models of molecular interaction networks can be transformed to include shared limiting resources by substituting ki, the reaction rate of a resource limited production, with keffi. Shown above an exemplary resource limited production are the detailed interactions between the substrate and the shared resource. The substrate Ai binds and unbinds the shared resource R with rates k+i and k-i, respectively. Binding of the substrate Ai to the shared resource R forms a complex Ci which releases the substrate Ai, the shared resource R and the product Bi with rate kcati. Under quasi steady-state assumptions the equivalent reaction rate keffi can be expressed as shown on the right. For simplification the parameters k+i, k-i and kcati are lumped into kmi. **(b) Limited shared translational resources reproduces non-monotonous dose-response in open-loop and incoherent feedforward circuit topologies.** On the left a graphical representation of a model for both the open-loop (OLP) and incoherent feedforward (IFF) topologies from Lillacci *et al.22* is shown. Transcriptional activation is modelled by a Hill-type function as shown. The solid arrows denote reactions assumed to follow the law of mass action. The model incorporates translational resources as introduced in panel A. These reactions are depicted as double headed arrows. The model was fit to data obtained by transiently transfecting HEK293T cells with increasing amounts of plasmid encoding tTA-Cerulean. The data and the fit are shown on the right. For a full description of the model see **Supplementary Note 3**. **(c) Limited shared translational resources reproduces non-monotonous dose-response in feedback and hybrid circuit topologies.** The model shown on the left is the same as in panel B with an additional negative feedback from miR-FF4 to tTA-mRNA. These topologies correspond to the feedback (FBK) and hybrid (HYB) topologies from Lillacci *et al*.22 The activation of gene expression by tTA-Cerulean is modeled by a Hill-type function as shown in the center of the figure. Reactions with double headed arrows denote resource limited production reactions as introduced in panel A. Solid arrows are assumed to follow the law of mass action. The model was fit to experimental data obtained from transient transfections with increasing amounts of plasmid encoding tTA-Cerulean. A full description of the model equations can be found in Supplementary Note 3. A description of the models can be found in **Supplementary Note 3** and the parameter values obtained by fitting are summarized in **Supplementary Table 7**. Data were analysed 48 hours after transfection and is plotted +/− SE. SE: standard error. N=3 biological replicates.

We demonstrated that the negative correlation is cell-type and promoter independent by expressing in H1299, U2OS, HeLa, HEK293T and CHO-K1 cell lines, *X-tra* and *capacity monitor* both driven by the constitutive promoter CMV, in molar ratios ranging from 1:1 to 2.5:1, for a total of 500 ng of encoding plasmids (**Supplementary Fig. 1-4, 14a**). Further, by using a PGK promoter^26^ that has different expression strength compared to CMV in HEK293T and H1299 (**Supplementary Fig. 5a**), we observed analogous outcomes (**Supplementary Fig. 5b,c**). Finally, by combining different molar ratios of mCitrine and mRuby3 encoding plasmids driven by two promoters of different strengths (EF1α or EFS) a similar behavior to Fig. 2a was observed (**Supplementary Fig. 6**).

**Fig. 5.**
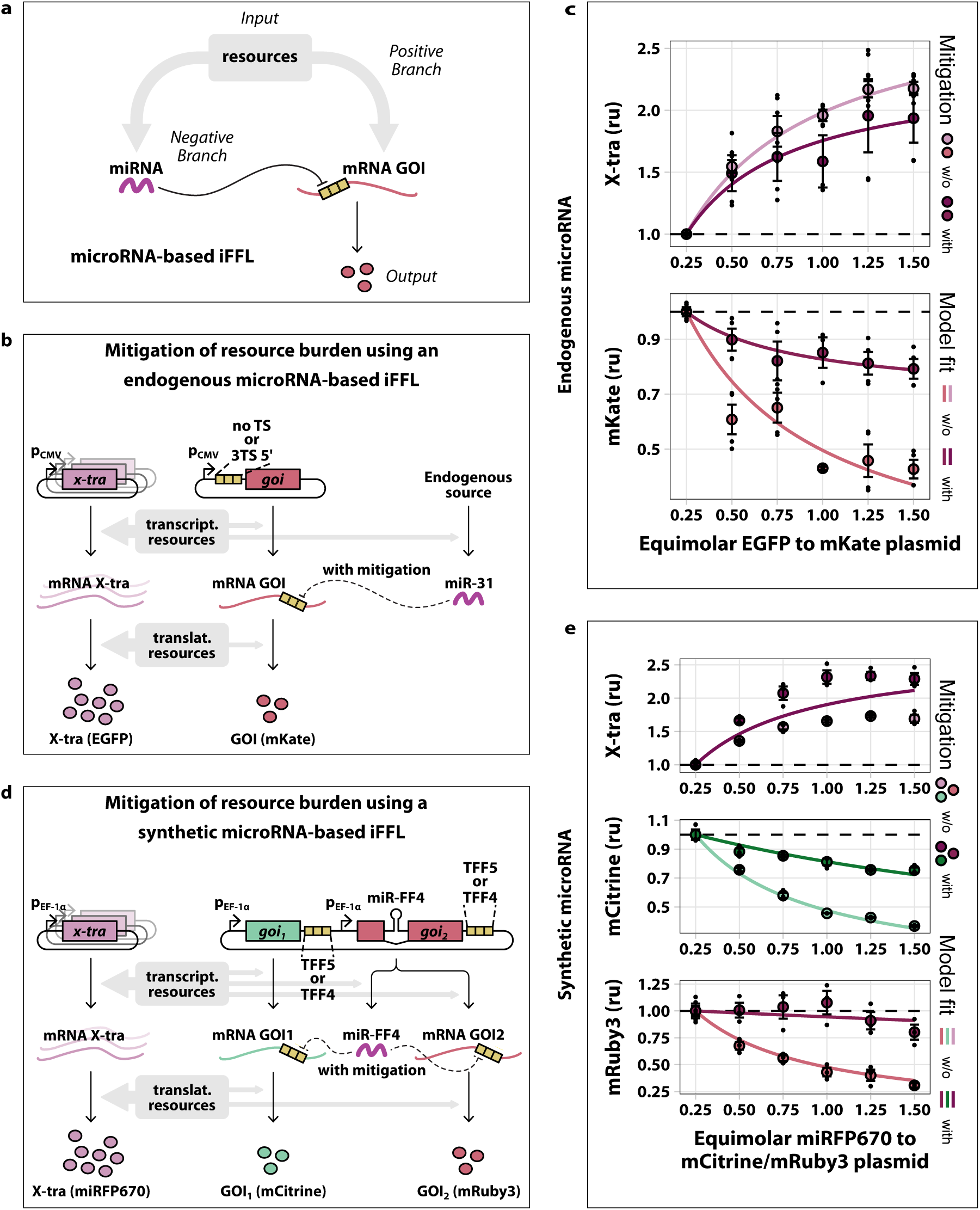
Mitigating the effects of resource limitation with microRNA-based iFFL. **(a)** The microRNA-based incoherent feedforward loop (iFFL) motif. **(b)** Schematic representation of the mitigation system based on endogenous microRNA. At high copies number of the *X-tra*, resources are drawn away from the production of the *GOI* and miR-31. By sensing the resource availability and repressing the *GOI* less when there are fewer resources, the miRNA reduces the effect of limited resources. **(c)** Two plasmids were co-transfected into H1299 cells which respectively express the *X-tra* gene (EGFP, panel b) and *GOI* (mKate, panel b) from strong constitutive CMV promoters. To limit the amount of resources available to the expression of mKate, the copy number of the X-tra plasmid was increased relative to the copy number of mKate. A mKate variant, which encodes three targets for the microRNA miR-31 in it’s 5’ UTR, mitigates effects due to resource sharing when compared to a variant lacking microRNA targets. The parameter values obtained by fitting are summarized in **Supplementary Table 8**. N=3 biological replicates. **(d)** Schematic of the mitigation system based on synthetic miRNA. In the presence of many copies of the *X-tra* gene, resources are drawn away from the production of both of the *GOIs* and the miR-FF4. Due to lower production of miR-FF4 the *GOIs* are less repressed. This compensates for the reduced availability of resources. **(e)** A plasmid encoding both the fluorescent protein mCitrine (*GOI*, panel d) and an intronically expressed microRNA (miR-FF4, panel d) under the control of strong constitutive promoters was co-transfected into HEK293T cells with increasing amounts of a plasmid expressing the *X-tra* gene (miRFP670, panel d) also under the control of a strong constitutive promoter. The impact of resource limitation on both *GOIs* was reduced when they contained three miR-FF4 targets in their 3’ UTRs compared to when they contained three mismatched miR-FF5 targets. The parameter values obtained by fitting are summarized in **Supplementary Table 9**. N=3 biological replicates. A description of the models can be found in **Supplementary Note 4**. Flow cytometry data were acquired 48h post-transfection and are plotted +/− SE. SE:standard error. ru: relative units.

As many synthetic circuits rely on tunable gene expression, we next tested resource competition on transcriptional inducible systems, by modulating *X-tra* repression with a Doxycycline (Dox)-repressed promoter (Fig. 2b) at different concentrations of Dox (from 0 to 1 μg/mL) while keeping *capacity monitor* amounts constant (Fig. 2b, **left**). Consistent with previous results, we observed that increased repression of *X-tra* corresponds to increased *capacity monitor* levels (Fig. 2b, **right**).

To exclude any experimental confounds as the source of our observations, we demonstrated that neither cell seeding nor nutrient supply had any apparent effect on the expression levels of the two genes, one of which was titrated whereas the second was held at a constant copy number (**Supplementary Fig. 7**).

These proof-of-concept experiments demonstrate that: *i)* gene expression in mammalian synthetic circuits is connected even in the absence of direct regulation, and *ii)* expression of exogenous genes is limited by cellular resource availability.

### Both transcriptional and translational resources are limiting

Since several different resource pools could be responsible for the observed effects described above, we set out to characterize the individual contributions of transcriptional and translational resource limitation to cellular burden. To evaluate potential limitations in transcriptional resources and the consequent gene competition for mRNA expression, we quantified mRNA levels in cells expressing *X-tra*/*capacity monitor* molar ratios from 1:1 to 2.5:1 in H1299 cells for a total of 500ng of plasmid DNA (corresponding protein data in **Supplementary Fig. 1**). We observed that as the *X-tra* mRNA increased, the *capacity monitor* mRNA levels decreased (Fig. 2c), supporting the hypothesis that shared transcriptional resources are indeed a limiting factor in mammalian synthetic gene co-expression.

To investigate whether the expression of endogenous genes is also affected by heterologous genetic payloads, we transfected H1299 cells with a plasmid encoding for EGFP and mKate under the control of a bidirectional promoter. We then sorted transfected cells according to *high* and *intermediate* levels of fluorescent markers, as well as *non-transfected* cells (absence of fluorescence) (**Supplementary Fig. 8**). We then quantified the mRNA levels of 3 endogenous genes (CyCA2, eIF4E, GAPDH, Fig. 2d). Notably, in transfected cells that express *high* and *intermediate* levels of EGFP and mKate, the expression of CyCA2, eIF4E and GAPDH decreases when compared to the *non-transfected* population. We also measured the mRNA levels of CyCA2, eIF4E and GAPDH in cells transfected with *X-tra*/*capacity monitor* molar ratios from 1:1 to 2:1 and observed a progressive, albeit not dramatic decrease with higher amounts of *X-tra* when compared to the 1:1 ratio (**Supplementary Fig. 9**). In this case cells were not sorted before mRNA extraction.

To provide further support to the observations on transcriptional burden on exogenous genes (Fig. 2c), we implemented a genetic circuit that can selectively overload the transcriptional resource pool without sequestering translational resources. The system is based on the self-cleaving hepatitis delta virus (HDV) ribozyme, which ensures that most of the transcribed mRNA is cleaved and thus destabilized (Fig. 2e, **left**). The circuit is composed of a single plasmid with two transcriptional units (TUs). One TU contains a tTA transcription factor co-expressed with the mRuby3 (*capacity monitor*) via the P2A peptide, driven by a constitutive promoter. The second TU includes the HDV-*X-tra* expression regulated by the TRE promoter. In this setup, Dox can be used to modulate the amount of burden imposed, similar to what was already shown in Fig. 2b.

We compared this circuit to a catalytically inactive mutant of the HDV ribozyme. As expected, we observed that when the HDV ribozyme is inactive, *X-tra* protein levels increase with decreasing amounts of Dox (**Supplementary Fig. 10, top pale pink bar**), whereas those of the *capacity monitor* decrease (Fig. 2e, **bottom pale blue bar**). In contrast, when the HDV ribozyme is active, *X-tra* expression is strongly reduced and only minorly increasing with lower Dox concentrations (**Supplementary Fig. 10, top dark purple bar**). Here, the *capacity monitor* levels decrease to a smaller extent than in the previous condition, supporting the observations in Fig. 2c that transcriptional resources are limited to a certain extent (Fig. 2e, **dark blue bar**). Interestingly, the expression levels of the *capacity monitor* with active HDV ribozyme are higher compared to the inactive mutant (**Supplementary Fig. 10, bottom dark blue bar**). We suggest that, assuming that the *X-tra* mRNA with an active HDV ribozyme is decapped and rapidly degraded, it is likely to sequester fewer translational resources, which should result in higher expression of the *capacity monitor*.

Transcriptional resource pool sharing is therefore at least partially responsible for the described gene expression trade-offs, and translational resources may represent an additional bottleneck to the overall expression of synthetic genes. We confirmed this hypothesis by adding a synthetic intron^27^ in the 5’UTR of the *X-tra* fluorescent protein (Fig. 2f, **top**). The synthetic intron enhances translation by augmenting mRNA export from the nucleus to the cytoplasm^27^ and therefore imposes specific translational load. Indeed, we observed higher expression of *X-tra* in H1299, HEK293T, HeLa, and U2OS cell lines in the presence of a synthetic intron, accompanied by lower *capacity monitor* levels (Fig. 2f, **Supplementary Fig. 11**), confirming that resources employed for translational regulation are also limiting. Thus our data collectively indicate that exogenous genes compete for resources both at the transcriptional and translational levels, overall imposing a gene expression burden on mammalian cells.

Since one of the goals in synthetic biology is output predictability, reproducibility and robustness, gene expression burden is a key issue to address. We reasoned that post-transcriptional and translational regulators, such as RBPs and miRNAs, may free up cellular resources^28^ by repressing target mRNA translation or inducing its degradation. If true, they could be exploited in more robust circuit topologies to reduce gene expression load, resulting in improved performance and predictability of engineered circuits. Therefore, we tested two RBPs, L7Ae and Ms2-cNOT7^29,30^ (Fig. 2g, **Supplementary Fig. 12-13**), as well as endogenous miRNAs, miR-31, miR-221 and miR-21 (Fig. 2h). For each system, a fluorescent protein encoding mRNA targeted by either RBPs or miRNAs (*X-tra*) was co-expressed with a second, constitutively expressed fluorescent readout (*capacity monitor*). L7Ae binds the 5’UTR of the *X-tra* mRNA inhibiting its translation, whereas Ms2 binds target sites in the 3’UTR of the *X-tra* transcript, allowing cNOT7 to cut the polyA tail to destabilize the target mRNA^29^. We tested the systems in U2OS, H1299, HEK293T and HeLa cell lines, and consistently observed that *X-tra* downregulation by RBPs results in increased levels of the *capacity monitor* (Fig. 2g, **Supplementary Fig. 12-13**).

miRNAs operate by either translation inhibition or mRNA degradation, according to complete^31^ or partial^32^ complementarity to the mRNA target. To evaluate the effect of miRNA regulation on cellular resources re-allocation, we placed three perfect complementary target sites (TS) in the 3’UTR of *X-tra*, which respond to the endogenous miR-31, miR-221, miR-21, highly expressed in H1299, U2OS, and HeLa cells respectively. The *capacity monitor* expression levels increased when the *X-tra* mRNA was downregulated by miRNAs, as compared to controls lacking miRNA target sites (Fig. 2h).

Due to their high productive capability^33^, CHO-K1 cells are the workhorses of the biopharmaceutical industry and have been extensively engineered to favor the expression of therapeutic proteins with desired post-translational modifications. We demonstrate that burden also affects these cells either by increasing *X-tra* gene amounts or enhancing its translation (**Supplementary Fig. 14a,b**). Redistribution of resources is also observed via regulation by the RBP L7Ae and the highly expressed endogenous miR-21 (**Supplementary Fig. 14c,d**).

These results confirm that post-transcriptional regulators can redistribute intracellular resources and importantly, that this phenomenon is cell-context independent. The extent of negative correlation between *X-tra* and *capacity monitor* expression, as well as the amount of repression by post-transcriptional regulators differs across cell lines; this could be the consequence of several factors, such as the relative abundance of transcriptional, post-transcriptional and translational resources.

A major advantage of miRNAs over RBPs is that they are endogenously expressed and cell-line specific. Thus, their expression does not impose an additional burden, and since several thousand endogenous miRNAs with different target sites are naturally present in mammalian cells ^34^, the design space is rather large, giving rise to a tremendous number of circuits that can be easily tailored to the cell/tissue of interest. Based on the results presented here, we envision that genetic circuits that mitigate resource competition via miRNAs may be designed for any mammalian cell line with a very broad set of potential applications.

### Characterizing the effect of miRNAs on resource distribution

We sought to characterize the correlation between miRNA-mediated downregulation and resource redistribution, by building a library of *miRNA sensors* for miR-31, which is endogenously expressed in H1299 lung cancer cells^35^. The *miRNA sensor* is composed of the fluorescent reporter mKate with or without miR-31 TS, encoded along with the *capacity monitor* (EGFP) on a single plasmid with a bidirectional promoter (Fig. 3a). The library includes 0, 1, or 3 fully complementary miR-TS in the 3’ or 5’UTR of mKate.

Similar to what was previously observed (Fig. 2g), when the *miRNA sensor’s* levels decrease as a consequence of miR-31 regulation, the expression of the *capacity monitor* increase*s*. The strongest repression was achieved with 3TS in the 5’UTR and was accompanied by corresponding higher capacity monitor levels (Fig. 3b). Conversely, when we rescued mKate expression by a miR-31 inhibitor (Fig. 3c **left, Supplementary Fig. 15 pink bars**), the *capacity monitor* levels decreased (Fig. 3c **right, Supplementary Fig. 15 dark blue bars**) demonstrating that miRNA sensor and *capacity monitor* levels are linked. Interestingly, the effect of the miRNA inhibitor was more pronounced with TS placed in the 3’UTR. Synthetic miRNA inhibitors bind to endogenous miRNAs in an irreversible manner^36^, but differences in their action (e.g. when target sites are placed in the 3’ *vs* 5’ UTR), as well as mechanistic insights into these differences, are still missing.

To confirm that miRNA-mediated resource redistribution is independent of experimental setting and plasmid design, we encoded the *miRNA sensor* and *capacity monitor* on two separate plasmids. Similar to previous results, *miRNA sensor* and *capacity monitor* were negatively correlated (**Supplementary Fig. 16a**), suggesting that cellular burden and miRNA-dependent resource reallocation are a common challenge and solution respectively. Downregulation of the *miRNA sensor* was also confirmed by qPCR (**Supplementary Fig. 16b**). Finally, when the *miR-31 sensor* was transfected in low miR-31 cell lines such as U2OS and HEK293T, neither the *miRNA sensor* nor the *capacity monitor* levels varied (**Supplementary Fig. 17**), further confirming the miRNA-dependent resource reallocation.

We showed in Fig. 2h that miRNA-dependent resource reallocation is observed across different cell lines, by expressing cell-specific *miRNA-sensors* which include 3TS in the 3’UTR. We then built a library of sensors with different numbers and locations of TS for miRNA-221 and −21 which are highly expressed in U2OS and HeLa cells respectively. We also confirmed here that *miRNA-sensor* and *capacity monitor* are inversely correlated, consistent with our observations in H1299 cells (**Supplementary Fig. 18-19**).

Overall these data show that miRNAs can be used to develop resource-aware plasmid-designs harbouring burden-mitigating circuit topologies, and that the number and location of TS can be tuned to achieve desired protein expression levels.

### A gene expression model that incorporates shared cellular resources recapitulates unexpected dose-response data

In order to provide a better understanding of our results, we developed a general resource-aware model, which offers a simple and convenient framework for extending existing models of biochemical reactions allowing them to incorporate the effects of shared limited resources. This framework follows ideas originally used to capture the competitive interaction of multiple inhibitors with an enzyme^37^ and has been applied to describe shared cellular resources in previous studies^15–18,38^.

Fig. 4a illustrates an overview of the framework. The main idea is to replace the rates of reactions that involve a shared resource with an *effective* reaction rate that captures the reduced availability of that resource due to the presence of competing genes. To create a distinction between regular reactions and resource-limited ones, we use double-headed reaction arrows to denote resource-limited reactions as illustrated at the bottom of Fig. 4a. This double-headed arrow summarizes the set of intermediate interactions shown in more detail at the top left of Fig. 4a. Here, the substrate A_i_ binds resource R with rate k^+^_i_ to form the complex C_i_. This reaction is also assumed to be reversible with rate k^-^_i_. With a rate k^cat^_i_ the complex gives rise to the product B_i_, while also freeing up both the substrate A_i_ and the resource R. We assume that the total amount of resource, R^total^, is conserved and remains constant at the time scale of the considered reactions. Considering all possible substrates that require resource R and assuming that C_i_ is in quasi-steady-state, the rate for resource-limited production can be expressed as k^eff^_i_, shown in the top right corner of Fig. 4a. k^eff^_i_ is a function of the total amount of resources and the current concentration of all substrates competing for this resource. This expression can be readily used to substitute all reaction rates that involve shared and limited resources.

In Lillacci *et al.*^22^, new synthetic control systems were introduced to enhance the robustness of gene expression. The same study also described a non-monotonic dose-response behavior in a simple inducible gene expression system. Such non-monotonic behavior is rather surprising and seems unintuitive, especially considering that conventional methods of modelling gene expression via Hill-type functions can only produce monotonically increasing (or decreasing) dose-response curves. Although Lillacci *et al.* proposed models for the different circuit topologies^22^, the effect of competition for limited resources was not considered. To demonstrate the effectiveness of our modelling framework, we extend these models to include limited resources and show that the resulting extended models recapitulate the experimental observations.

The four topologies considered in Lillacci *et al.*^22^ were split into two groups based on the presence of negative feedback from the fluorescent protein DsRed to the transcriptional activator (tTA). The first group consisted of the open-loop (OLP) and incoherent feedforward (IFF) topologies. In both these circuits, the constitutively expressed transcriptional transactivator, fused to the fluorescent protein Cerulean (tTA-Cer), activates the expression of the fluorescent protein DsRed. Furthermore, the gene of DsRed intronically encodes the synthetic microRNA FF4 (miR-FF4). In the IFF topology, the matched target of this microRNA is present in the 3’ untranslated region (UTR) of the DsRed gene. This target is replaced by a mismatched target for the microRNA FF5 in the OLP. These detailed interactions are depicted here in Fig. 4b **left side**. To observe potential shifts in the allocation of resources, we generated dose-response curves by increasing the amount of transfected tTA-Cer plasmid, while the other two plasmids, containing DsRed and the constitutively expressed fluorescent transfection reporter mCitrine, were held constant. As can be seen from the model fit, plotted as a solid line in the data graph, the extended model reproduces the non-monotonic behavior of the dose-responses (Fig. 4b, **right**).

The second group of topologies considered by Lillacci *et al.* consisted of the feedback (FBK) and the FBK+IFF hybrid (HYB) topologies. In addition to all the interactions described for the OLP and IFF circuits, the FBK and the HYB circuits possess miR-FF4 targets in the 3’ UTR of the tTA-Cer gene, which introduces negative feedback. Furthermore, the FBK and HYB differ from each other by the presence of a matched target for miR-FF4 in the HYB topology, which introduces incoherent feedforward and is replaced by a mismatched FF5 target in the FBK circuit. All the interactions are illustrated in detail in Fig. 4c, **left**. The dose-response curves for the two circuits were obtained as described above. Again, the fit of the extended model to the data captures its rather unexpected behavior (Fig. 4c, **right**).

Our simple framework adapts existing models of gene expression to include pools of shared and limited resources. We show that it can be used to provide an explanation for unintuitive dose-responses in tTA-based circuits. With this framework as a tool, we believe that performance issues attributed to gene expression burden can be addressed head-on in the design phase of circuit-building, thereby reducing the need for costly subsequent build-test-learn iterations.

### Mitigating cellular burden with Incoherent Feed-Forward Loop (iFFL) circuits

We implemented a strategy that exploits miRNA to reduce the indirect coupling between co-expressed genes. In particular, we took advantage of the fact that miRNA production also requires (pre-translational) cellular resources, therefore acting as a sensor for resource availability. Because of this, it is possible to reduce the coupling between genes co-expressed via a common resource pool by introducing miRNA-mediated repression of those genes (as long as the miRNA itself is also affected by the same resource pool). Since both the miRNA and the miRNA-repressed gene are affected by the availability of resources, miRNA-mediated repression implements an incoherent feedforward loop similar to previously published circuits^22,25,39^ (Fig. 5a). Interestingly, this iFFL-based circuit constitutes a biological implementation of the miRNA circuit proposed by Zechner *et al*^40^. In this setting, the miRNA can be interpreted as an estimator of its cellular context (e.g. amount of free resources) and acts to filter out this context, thereby minimizing its impact on the output of interest.

We explored this strategy for an endogenously expressed miRNA (Fig. 5b,c) and a synthetic miRNA encoded on a plasmid (Fig. 5d,e). More specifically, Fig. 5b describes a strategy that exploits endogenous miRNAs to reduce the coupling of a gene of interest (GOI) to the expression level of other genes, introduced by the limitation in resources. Implementation of this strategy only requires adding target sites of an endogenous miRNA to the 5’ UTR of the gene of interest (mKate). In our experimental setup, when the copy number of a second gene *(X-tra)* is increased, resources are drawn away from the expression of mKate and allocated to the expression of *X-tra*. The shift in resource allocation is expected to also affect the miR-31, which acts as a *capacity monitor*. This leads to a reduction in the repression of mKate, effectively compensating for the burden imposed by the co-expression of the *X-tra* gene.

To demonstrate this mitigation approach experimentally, we co-transfected H1299 cells with increasing amounts of EGFP (*X-tra*), along with a constant amount of mKate (*GOI*) that either includes (for mitigation) or omits (no mitigation) three miR-31-TS in the 5’UTR. As expected, the expression level of *X-tra* approached saturation as the plasmid copy number increased, both for the targeted and non-targeted *GOI* variants (Fig. 5c). In agreement with previous results, the expression of the non-targeted *GOI* strongly decreased with increased expression of *X-tra*. Conversely, the decrease in expression of the targeted *GOI* was only about a third of that of the non-targeted variant, indicating improved adaptation to changes in resource availability (Fig. 5c, **Supplementary Fig. 20**). This observation was also captured well by a model of the system that explicitly considered resources, as described in the previous section. Analogously, miR-221-iFFL circuits specific for U2OS (**Supplementary Fig. 18)** and HEK293T cells^41^ (**Supplementary Fig. 22**) show improved robustness to burden imposed by increasing exogenous gene load (**Supplementary Fig. 21, 23**).

Importantly, the delivery of genetic payloads also affects the expression of endogenous genes (CyCA2, elF4E and GAPDH), as shown in Fig. 2d. We then sought to compare the expression of the same endogenous genes in the presence or absence of miR-31 sensor in H1299 cells. After 48h from transfection of EGFP and mKate on a bidirectional plasmid, with mKate either including (miRNA sensor) or not (noTS) target sites for miR-31, we sorted cells according to high, intermediate or absence of fluorescence expression (**Supplementary Figure 24a**) and performed qPCR. Curiously, we observed that in cells transfected with miR-31-sensor, the decrease in the expression of the endogenous genes was much lower than in its absence (**Supplementary Figure 24c**). Furthermore, the expression of endogenous genes was inversely proportional to the levels of fluorescent proteins (**Supplementary Figure 24b**). Thus, the lower expression of endogenous genes due to the burden imposed by exogenous payloads is counteracted by the miR31-sensor.

Motivated by our desire to achieve portability across cell lines and multiple-output regulation are desired features of synthetic devices, we implemented and tested a synthetic miRNA-iFFL circuit that tunes three GOIs (Fig. 5d). Similar to the endogenous case, the genes-of-interest, mCitrine (*GOI_1_*) and mRuby3 (*GOI_2_*), encode target sites for the miRNA-FF4 in their 3’ UTRs. In contrast to endogenous miRNA expression, however, here the miRNA is expressed intronically from *GOI_2_*. In this way, the circuit forms a self-contained unit that can be easily transferred between cell types.

We co-transfected HEK293T cells with a plasmid encoding constitutively expressed miRFP670 (*X-tra*) and a plasmid composed of two transcriptional units (TU), each expressed under the constitutive promoter EF1α (Fig. 5d). The first TU encodes the mCitrine, whereas the second drives mRuby3. Furthermore, the 3’ UTR of mRuby3 contained either three target sites for the synthetic miRNA-FF4 or three mismatched miR-FF5 target sites (negative control). The miRNA-FF4 was intronically encoded in the mRuby3 gene. Identically to the endogenous case, the amount of *X-tra* plasmid was increased while keeping the *GOIs’* plasmid constant. Again, expression of *X-tra* increased and approached saturation with increasing molar amounts and consequently, the non-targeted variants of the *GOIs* decreased (**TFF5 in** Fig. 5e). Conversely, the expression of the targeted variants (**TFF4 in** Fig. 5e) decreased to a lesser extent than the non-targeted ones, analogously to what was observed for endogenous miRNAs, albeit with lower efficiency. Finally, to demonstrate the portability of the device we tested the approach in mouse embryonic stem cells (**Supplementary Fig. 25**). Here, adaptation to shifts in resource availability was similar to the endogenous miRNA-based regulation (Fig. 5d). Thus, we showed that also in entirely synthetic systems, adaptation to shifts in resource availability was achieved. To ensure that the observed mitigation was not caused by a higher tolerance to changes in availability at lower expression levels, we showed analytically using the described modelling framework that the normalized expression at lower levels was more sensitive to burden (**Supplementary Note 2**).

Overall, these results suggest that our approach can be used to mitigate resource-mediated coupling of gene expression despite cell-to-cell variability, demonstrating the portability and broad applicability of our findings. Our results demonstrate that iFFL circuits can mitigate burden from transgene expression in mammalian cells. Importantly, by using miRNA one can either opt for endogenous miRNAs to specifically tailor a circuit to a desired cell line, or create a portable circuit by using a synthetic miRNA such as miR-FF4.

## Discussion

Our study demonstrated for the first time that the sharing of limited cellular resources represent a general bottleneck for the predictability and performance of synthetic circuits in mammalian cells, with important consequences for mammalian synthetic biology, molecular biology, as well as for biotechnology applications. We showed that heterologous gene expression imposes a resource burden, independent of cell type (six different cell lines were tested) and of promoter strength (Fig. 2a,b, **Supplementary Fig. 1,5-6**), although it is more severe with high plasmid copy number (Fig. 2a).

Further, we presented a detailed characterization of the distinct contributions of transcriptional and translational burden by combining molecular biology techniques (Fig. 2c-d) with the engineering of simple genetic motifs that enabled the specific control of translation of the desired gene (Fig. 2e,f). Notably, we showed that endogenous and synthetic post-transcriptional and translational regulators such as RNA-binding proteins and miRNAs can effectively redistribute cellular resources (Fig. 2g,h-3), an observation we exploited to engineer resource-aware synthetic circuits that utilize post-transcriptional regulation to mitigate burden.

We showed that endogenous genes are also affected by the burden imposed by genetic payloads (Fig. 2d), whereas this effect was less pronounced in the presence of the miRNA sensor (Fig. 3a **middle, Supplementary Fig. 24c**). We speculate that this positive effect of miRNA sensor is attributed to the freeing up of translational resources, leading to the increase of proteins involved in the transcription of endogenous genes. At the same time, as already proposed in *Gambardella et al.*^42^, the downregulation of mKate by miR-31 may lead to a ‘queueing effect’ for the degradation of the other mRNAs, similar to what was shown with two independent proteins tagged for degradation by the proteasome^43^.

To get a deeper understanding of the mechanisms through which resource sharing dynamically couples gene expression and to explain resulting non-intuitive behavior of genetic circuits induced by such coupling in mammalian cells^22^, we described a modelling framework that captured the indirect interdependence of gene expression in a resource-limited context. Our framework for introducing resource-limited reactions into pre-existing models of synthetic circuits provides a straightforward way of obtaining resource-aware models and facilitates the design of circuits that are less prone to burden effects. For instance, the family of controllers analyzed in Fig. 4 exemplifies the non-intuitive behavior that might arise due to burden. Here, limiting resources led to a non-monotonic dose-response in a simple transcription factor and reporter pair. Our extended modelling framework successfully captured this unexpected behavior of the dose-responses (Fig. 4b), which could not be explained by traditional models. This same modelling framework suggested that an incoherent feedforward loop (iFFL) is a particularly well-suited circuit motif for mitigating burden effects. The incoherent feedforward loop (iFFL) itself is one of the core gene regulatory motifs in biology, and unsurprisingly it has served as inspiration for many synthetic genetic circuits that exploit its adaptation properties, particularly to input perturbations^22,25,39,44^. This together with our experimental observations on the effective role of miRNA in resource distribution (Fig. 2h, 3) led us to choose a specific implementation of this motif based on miRNAs for burden mitigation in mammalian cells (Fig. 5).

Given its importance in synthetic circuits, resource burden has recently been studied in *E. coli*^12,16,19,20^, where two different synthetic circuits have been used for burden mitigation. In Ceroni *et al*.^20^, the mitigation strategy aims to reduce the expression of the gene of interest to avoid burden, while the strategy in Huang *et al*.^19^ aims to maintain the expression of a gene of interest upon a reduction of resources, possibly at the expense of other genes. Both strategies rely on negative feedback, a regulatory circuit motif that is widely used in natural and synthetic circuits. While iFFL is also a common regulatory motif, iFFL circuits have not to date been utilized in burden mitigation, in spite of several distinct advantages over negative feedback circuits for burden mitigation. In particular, iFFL circuits are considerably simpler to implement and much easier to tune than negative feedback circuits, which usually require more components and can become dynamically unstable if not properly designed and tuned. In terms of dynamic response, iFFL circuits are also generally faster in rejecting disturbances like a sudden change in resource availability. Indeed iFFL regulation responds to the disturbance itself, while negative feedback begins to act only after the impact of the disturbance on the regulated output has been detected.

In mammalian cells, burden-mitigating circuits have not been proposed until *this* and another accompanying study by Jones *et al.*^45^ (Nature Biotechnology joint submission). While both studies independently reached the conclusion that iFFL motifs are ideally suited for burden mitigation in mammalian cells, the two studies follow complementary paths for the circuit implementation of this motif, with both implementations successfully demonstrated in multiple cell lines. In the Jones *et al.* study, an endoribonuclease-based iFFL is used to mitigate the effects of resource burden. While the use of RNA-binding proteins to implement iFFL circuits in mammalian cells is not new^46^, the use of endoRNAse for load mitigation in Jones *et al.* was not previously reported. One feature of RNA-binding protein iFFL circuits is that they do not rely on other factors like RISC for their function. Another feature of RNA binding protein mitigation implementations is that they respond directly to changes in translational resources, even though they were designed for transcriptional resource mitigation.

In this study we adopt a miRNA implementation of iFFL circuits for the purpose of burden mitigation. miRNA-iFFLs have been demonstrated to increase robustness to gene dosage variability^22,44^ and gene expression noise^25,39^, but their use for burden mitigation in any cell type is novel. After considering both RNA-binding proteins (L7Ae and Ms2-cNOT7) and miRNA for iFFL implementations (Fig. 2g,h), we opted for a miRNA-based one (both endogenous and synthetic) due to several considerations. Importantly, miRNA circuits do not use genetic components that derive from different organisms, circumventing potential toxicity and immunogenicity concerns that could limit their application in medical therapy^47,48^. Additionally, as we highlight next, miRNA-iFFLs exhibit high versatility, faster dynamics^49^, and other distinctive features that are important for biotechnological applications.

To date, endogenous miRNA have been used as inputs to synthetic cell type classifier circuits^50,51^ but their use as incoherent feedforward signals is unprecedented and provides important advantages over protein-based implementations. Specifically, iFFL circuits that exploit endogenous miRNAs do not require the expression of additional components (no additional translational burden is imposed), enable cell-type *specificity*^50,52^ and can enhance stabilization against perturbation by inserting target sites for one or more endogenous miRNAs (*scalability*). At the same time, synthetic miRNAs enable *portability* of circuits across different cell lines, and the flexibility at the sequence level allows scaling up to many orthogonally operating circuits. Additionally, synthetic miRNA-based iFFLs are compact and can be easily included in pre-existing genes as introns containing the miRNA scaffold (*minimal footprint* on the transcriptional resources of the cell), making them suitable for therapeutic applications such as gene therapy. For example, while protein-based iFFLs require incorporation of much longer sequences, miRNA-based iFFL could be used in AAV-mediated gene therapy because their smaller sequences well fit within the maximum cargo capacity of 4.5 kb of AAV vectors^53^.

Another critical feature of miRNAs is their *programmability*. The specificity of a miRNA can be easily engineered to target any synthetic or endogenous gene without the need to engineer the target itself^54–56^. This again has implications for medical applications, where the target might not be accessible for modification. One possible limitation here is that the use of the same miRNA to regulate multiple targets might lead to depletion of the miRNA, giving rise to an inevitable trade-off. This trade-off is observed for competing endogenous RNA (ceRNA), which are known to naturally regulate other RNAs by competing for miRNA-binding. The principle of ceRNA has been demonstrated synthetically on a gene circuit^57^. Finally, *tunability* of repression strength can be easily achieved both through the number and the placement of the targets (Fig. 3). Such tunability of repression can be used to enhance adaptation to variations in resource availability. It should be noted that stronger repression will yield lower expression levels of the gene of interest. This tradeoff is unavoidable, and inherent to all implementations of iFFL burden mitigation circuits, including endoRNase implementations.

In terms of burden mitigation, at first thought one might argue that miRNA-based iFFL would only mitigate coupling through shared resources on the transcriptional level since translational resources do not affect miRNA production. However, given that several proteins are involved in the maturation, transportation, and activity (e.g. RISC multi-protein) of miRNAs and these proteins are impacted by translational resource availability, this would indirectly allow miRNA-based iFFLs to also mitigate translational burden. In any case, translational burden mitigation may be easily incorporated *directly* by expressing the miRNA under the control of a synthetic or endogenous transcription factor. Implementation considerations aside, we point out that due to the differences and unique features of both miRNA and endoRNase implementations, we expect that in the future it will be desirable to combine them in single cells to create an even larger set of simultaneously operating burden-mitigating circuits.

Understanding the impact of resource availability will have important consequences for biological studies and for improved mammalian cell engineering. For example, for successful treatment, cell therapies that rely on the fine expression and secretion of therapeutic molecules can now be designed with resource-aware circuits. Our findings suggest that, when choosing a host cell line, one of the key factors to consider should be its transcriptional and translational capacity^21^, not only in terms of productivity, but also in terms of the ability of the cells to maintain their fitness over the production phase. In this respect, enhancing the insulation of synthetic genetic circuits from cellular metabolism is paramount. In our study, we implemented a system that could efficiently sustain the production of up to three (and probably more) output proteins (Fig. 5), reducing the unwanted coupling among their expression levels and with other cellular processes. However, it should be noted that our system was not intended to reach the highest expression levels, as its purpose was instead to allow the concurrent production of several output proteins in a predictable manner.

Finally, studies of biological functions that employ system perturbations by exogenous gene expression often lack accuracy and exhibit highly variable results due to less-than-optimal genetic circuit designs. This is critical when performing quantitative analyses, for which predictability and reproducibility are essential. Our study presents a portable design capable of enhancing the insulation of transgene expression and will thus contribute to the development of robust-by-design mammalian synthetic circuits, with important implications for basic science and applications in industrial biotechnology and medical therapy.

## Methods

### Cell culture

HEK293T, U2OS and HeLa cells (ATCC) used in this study were maintained in Dulbecco's modified Eagle medium (DMEM, Gibco); H1299 were maintained in Roswell Park Memorial Institute medium (RPMI, Gibco); CHO-K1 were maintained in Minimum Essential Medium α (α-MEM, Gibco). All media were supplemented with 10% FBS (Atlanta BIO), 1% penicillin/streptomycin/L-Glutamine (Sigma-Aldrich) and 1% non-essential amino acids (HyClone). HEK239T cells (ATCC, strain number CRL-3216) used for part of this study were maintained in DMEM (Sigma-Aldrich or Gibco) supplemented with 10% FBS (Sigma-Aldrich), 1X GlutaMAX (Gibco) and 1 mM Sodium Pyruvate (Gibco). E14 mouse embryonic stem (mES) cells were grown in DMEM (Gibco) supplemented with 15% FBS (PAN Biotech; specifically for ES cell culture), 1% penicillin/streptomycin (Sigma-Aldrich), 1% non-essential amino acids (Gibco), 2 mM L-Glutamine (GlutaMAX; Gibco), 0.1 mM beta-mercaptoethanol (Sigma-Aldrich) and 100 U/mL Leukemia inhibitory factor (LIF; Preprotech). At every passage the media was additionally supplemented with fresh CHIR99021 to 3 uM and PD0390125 to 1 uM to support naïve pluripotency (2i conditions^58^). All labware used was coated with 0.1% gelatin (prepared ourselves) prior to plating the ES cells. The cells were maintained at 37 °C and 5% CO2.

### Transfection

Transfections were carried out in 24-well plate for flow cytometry analysis or in a 12-well plate format for flow cytometry and qPCR analysis run on the same biological replicates (**Supplementary Table 1**). Transfections for Fig. 2d and **Supplementary Fig. 8** were carried out in 6 cm dishes. H1299, HeLa, U2OS, HEK293T and CHO-K1 cells were transfected with Lipofectamine® 3000 (ThermoFisher Scientific) according to manufacturer’s instructions and 300 ng total DNA (500 ng in Fig. 2c,d and **Supplementary Fig. 1-5,8,9**) in 24-well plates. DNA and transfection reagents were scaled up according to the Lipofectamine® 3000 manufacturer’s instructions. miR-31 inhibitor (Invitrogen™ *mir*Vana™ miRNA Inhibitors) was cotransfected using the same method as for DNA (Fig. 3c).

HEK293T cells used for experiments shown in Fig. 2a,b,e, 4 and 5e were plated approximately 24h before transfection at 62500 to 75000 cells per well in 24-well plates. The transfection solution was prepared using Polyethylenimine (PEI) “MAX” (Mw 40000; Polysciences, Inc.) in a 1:3 (ug DNA to ug PEI) ratio with a total of 500 ng of plasmid DNA per well. Both DNA and PEI were diluted in Opti-MEM I reduced serum media (Gibco) before being mixed and incubated for 25 minutes prior to addition to the cells. E14 mouse embryonic stem cells were transfected using Lipofectamine® 2000 (ThermoFisher Scientific) in a 1:3 (ug DNA to ug Lipofectamine® 2000) with 300 ng of plasmid DNA per well. The transfection was performed on cells in suspension immediately after plating at approximately 30000 cells per well. All wells were coated with 0.1% Gelatin before the addition of the cells.

### Flow cytometry and data analysis

H1299, HEK293T, U2OS, HeLa and CHO-K1 cells were analyzed with a BD Facsaria™ cell analyzer (BD Biosciences) or BD Celesta™ cell analyzer (BD Biosciences) using 488 and 561 lasers. For each sample >20000 singlet events were collected and fluorescence data were acquired with the following cytometer settings: 488 nm laser and 530/30 nm bandpass filter for EGFP, 561 nm laser and 610/20 nm filter for mKate. Cells transfected in 12-well plates were washed with DPBS, detached with 100 μL of Trypsin-EDTA (0.25%) and resuspended in 600 μL of DPBS (Thermo Fisher). 200 μL of cell suspension were used for flow cytometry and 400 μL for RNA extraction. HEK293T used for experiments shown in Fig. 2a,b,e, 4 and 5e cells were measured 48 hours after transfection on a BD LSRFortessa™ Special Order and Research Product (SORP) cell analyzer. mCitrine fluorescence was excited via a 488 nm laser and was detected through a 530/11 nm bandpass filter. mRuby3 was excited via 561 nm laser and measured through a 610/20 nm bandpass filter. miRFP670 was excited at 640 nm and measured through a 670/14 nm bandpass filter. E14 mES cells were measured 48 hours after transfection on a Beckman Coulter CytoFLEX S flow cytometer. mCitrine fluorescence was excited using a 488 nm laser and was detected through a 525/40+OD1 bandpass filter. mRuby3 was excited with 561 nm laser light and measured through a 610/20+OD1 bandpass filter. miRFP670 was excited at 638 nm and measured through a 660/10 bandpass filter. The cells were collected for measurement by washing with DBPS (Sigma-Aldrich or Gibco) and detaching in 70 to 180 μL of Accutase solution (Sigma-Aldrich). For each sample between 10000 to 200000 singlet events were collected. Fluorescence intensity in arbitrary units (au) was used as a measure of protein expression. For each experiment a compensation matrix was created using unstained (wild type cells), and single-color controls (mKate/mCherry only, EGFP only or mCitrine only, mRuby3 only, miRFP670 only). Live cell population and single cells were selected according to FCS/SSC parameters (**Supplementary Fig. 26, 27**). Data analysis was performed with Cytoflow or a custom R script. Data fitting was performed using Mathematica’s NonlinearModelFit function and the InteriorPoint method.

### Cell sorting

H1299 cells used for the experiment shown in Fig. 2d were trypsinized from 6 cm dishes and counted. They were then centrifuged at 500 g for 5 min and resuspended at a concentration of 5 mln/mL in sorting buffer (PBS 1x + 3 mM EDTA + 0.8 % Trypsin + 1 % FBS). Cells were sorted with a BD FACSMelody™ cell sorter according to their fluorescence levels (**Supplementary Fig. 8**). 150,000 cells per gate were collected.

### DNA cloning and plasmid construction

Plasmid vectors carrying gene cassettes were created using In-Fusion HD cloning kit (Clonetch), Gibson Assembly, via digestion and ligation or using the yeast toolkit (YTK)^59^ with custom parts for mammalian cells. Gibson Assembly master mixes were created from Taq DNA Ligase (NEB), Phusion High-Fidelity DNA Polymerase (NEB) and T5 Exonuclease (Epicentre) in 5X isothermal buffer (**Supplementary Table 6**). Ligation reactions were performed in 1:2-5 molar ratios of plasmid backbone:gene insert starting with 50 to 100 ng of vector backbone digested with selected restriction enzymes. Assemblies using the YTK were performed according to the original publication^59^. Newly created constructs were transformed into XL10-Gold or TOP10 *E. coli* strains.

For plasmids with miRNA target sites, the target sequences were selected using miRBase database (http://www.mirbase.org/) and are listed in **Supplementary Table 4**. List of oligos used to clone endogenous miRNAs TS are listed in **Supplementary Table 3**. All plasmids were confirmed by sequencing analysis and deposited to addgene.

To perform western blot analysis, a His-tag composed of 6 Histidine residues was inserted after the start codon of mKate encoding plasmids.

### mRNA extraction and reverse transcription

RNA extraction was performed with E.Z.N.A.® Total RNA Kit I (Omega Bio-tek). The protocol was followed according to manufacturer’s instructions and RNA was eluted in 30 μL of RNAse free water. RNA samples were conserved at −80°C.

PrimeScript RT Reagent Kit with gDNA Eraser – Perfect Real Time (Takara) was used according to manufacturer’s instructions. The protocol was performed on ice in a RNAse free environment to avoid RNA degradation. A negative control without PrimeScript RT Enzyme Mix I was always prepared to investigate genomic DNA contamination.

### qPCR

Fast SYBR Green Master Mix (ThermoFisher Scientific) was used to perform qPCR of cDNAs obtained from 500 ng of RNA and diluted 1:5. Samples were loaded in MicroAmp™ Fast Optical 96-Well Reaction Plate (0.1 mL) and the experiment was carried out with a CFX96 Touch Real-Time PCR Detection System (BioRad) machine. Each well contained 20 μL of final volume (7 μL SYBR Green Master Mix, 10 μL ddHO, 1 μL of each primer, 1 μL of template). Also, a control without template (blank) was set. Primers were designed to amplify a region of 60-200 bp (**Supplementary table 5**) and with a temperature of annealing between 50 and 65 °C 72. Data were analysed using the Comparative Ct Method according to Applied Biosystems Protocols.

## Acknowledgements

We thank Dr. Gabiele Lillacci for sharing the plasmids expressing the synthetic circuit topologies as well as the plasmid expressing mCitrine, Dr. Maaike Welling for sharing the mES E14 cell line. We thank Daniela Perna and Luca Giorgio Wanderlingh for their technical support. GB.S. gratefully acknowledges support from his UK EPSRC Fellowship for Growth in Synthetic Biology (EP/M002187/1) and his UK Royal Academy of Engineering Chair in Emerging Technologies for Engineering Biology.

## Author contributions

T.F., F.C., GB.S., M.K. and V.S. conceived the project. F.C. designed and performed experiments and performed data analysis of endogenous miRNAs, RBPs, transcriptional and translational burden, endogenous genes’ expression. T.F. designed and performed experiments and performed data analysis of synthetic miRNA, generic and transcriptional burden (HDV). T.F. and J.G. developed the mathematical model. F.T. and F.C. performed and analyzed iFFL experiments with endogenous miRNAs. M.K. and V.S. supervised the experimental work and secured funding. M.K. and GB.S. supervised the computational work. T.F., F.C., GB.S., M.K., and V.S. wrote the manuscript. All authors edited the manuscript.

## Data availability

All relevant data are included as Source Data and/or are available from the corresponding author on reasonable request. Plasmid sequences are deposited on AddGene and GenBank under the accession codes specified in **Supplementary Table 2**. Strains and plasmids used in this study are available from the corresponding author on reasonable request.

## Code availability

The code used for automated analysis and fitting is available on reasonable requests from the corresponding authors.

