## Supplementary Information for "Characterization, modelling and mitigation of gene expression burden in mammalian cells"

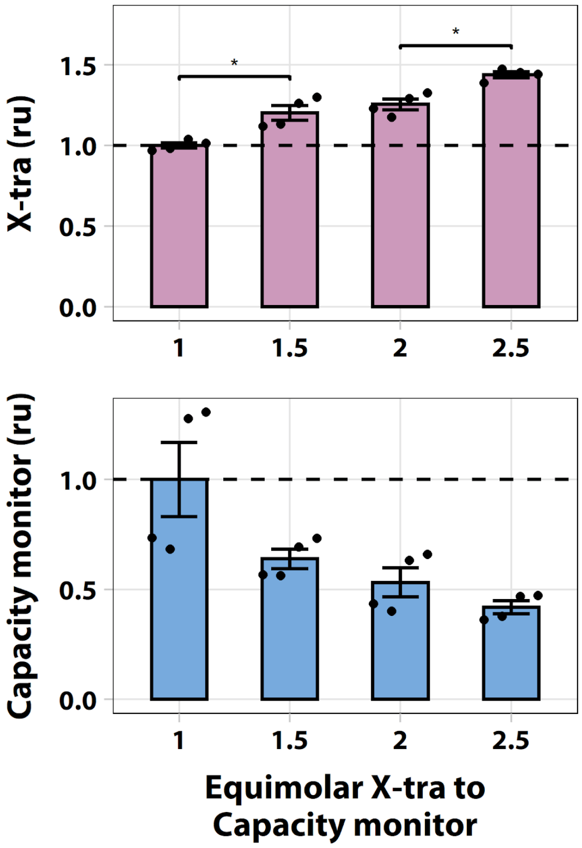

**Supplementary Figure 1. Titration of *X-tra* gene in H1299 cells.** Flow cytometry results of H1299 cells co-transfected with fixed amount of *capacity monitor* (mKate) and increasing *X-tra* (EGFP) at molar ratio from 1:1 to 1:2.5, both under CMV promoter regulation. Data correspond to samples collected for qPCR of **Fig. 2c** and were acquired 48 hours post-transfection. Data were normalized to mean fluorescence values at the plasmid molar ratio of 1.00 +/- SE. SE:standard error. ru: relative units. N=4 biological replicates. unpaired T-test. p-value: * < 0.05.

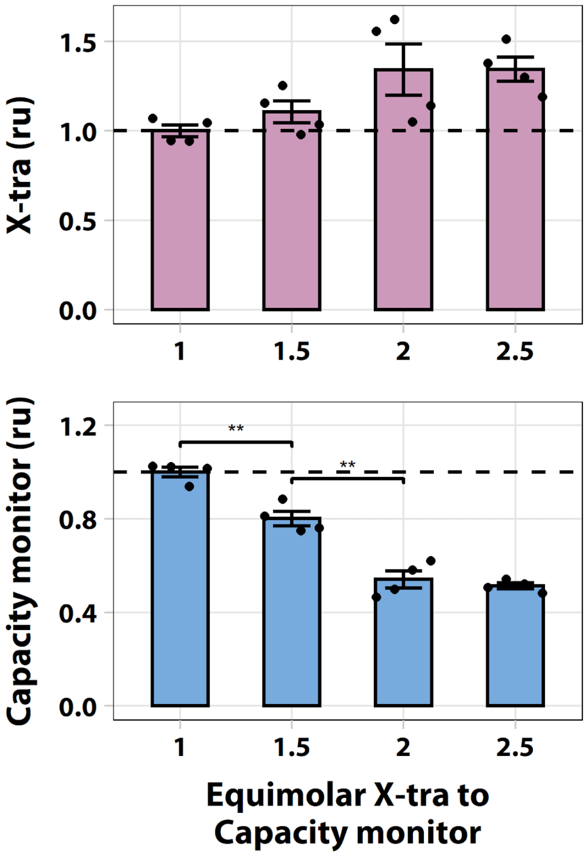

**Supplementary Figure 2. Titration of *X-tra* gene in U2OS cells.** Flow cytometry results of U2OS cells co-transfected with fixed amount of *capacity monitor* (mKate) and increasing *X-tra* (EGFP) at molar ratio from 1:1 to 1:2.5, both under CMV promoter regulation. Flow cytometry data were acquired 48 hours post-transfection. Data show the mean fluorescence normalized to its value at a plasmid molar ratio of 1. Error bars represent the standard error SE. ru: relative units. N=4 biological replicates. unpaired T-test. p-value: ** < 0.005.

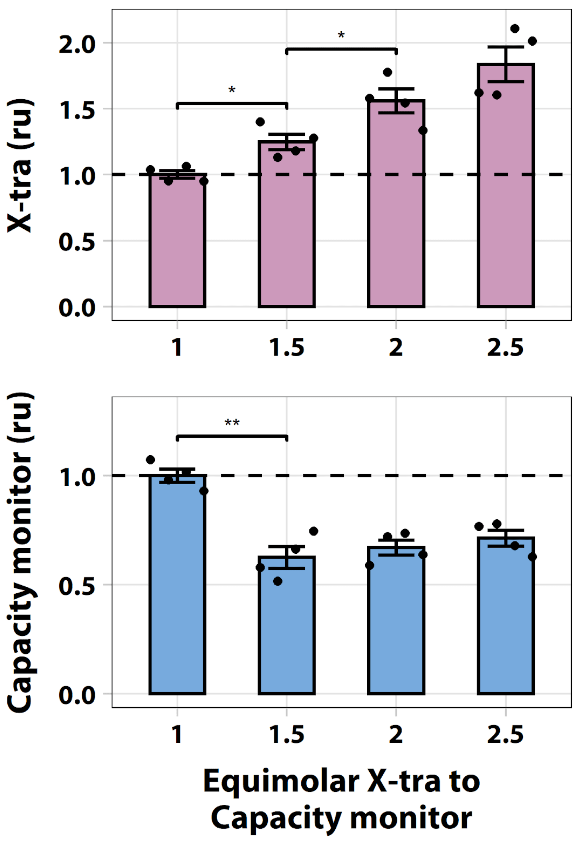

**Supplementary Figure 3. Titration of *X-tra* gene in HeLa cells.** Flow cytometry results of HeLa cells co-transfected with fixed amount of *capacity monitor* (mKate) and increasing *X-tra* (EGFP) at molar ratio from 1:1 to 1:2.5, both under CMV promoter regulation. Flow cytometry data were acquired 48 hours post-transfection. Data show the mean fluorescence normalized to its value at a plasmid molar ratio of 1. Error bars represent the standard error. ru: relative units. N=4 biological replicates. unpaired T-test. p-value: ** < 0.005, * < 0.05.

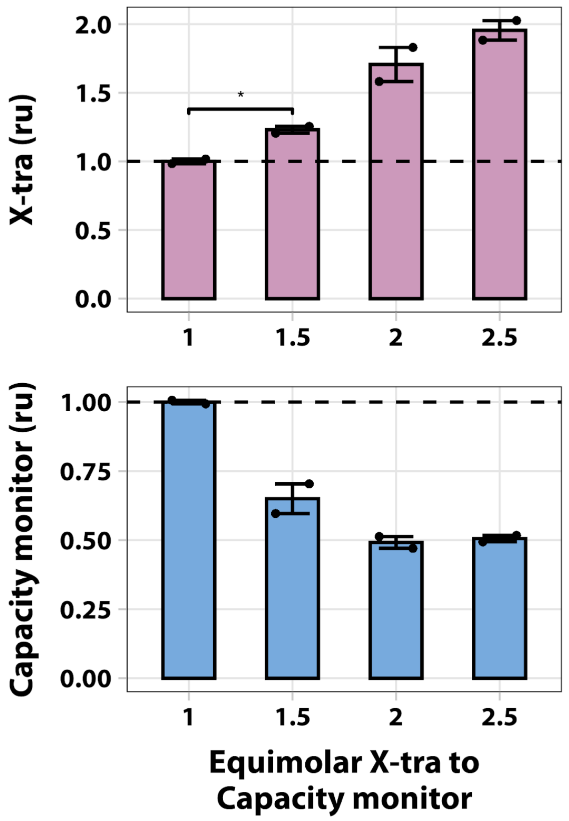

**Supplementary Figure 4. Titration of *X-tra* gene in HEK293T cells.** Flow cytometry results of HEK293T cells co-transfected with fixed amount of mKate (*capacity monitor*) and increasing amount of EGFP (*X-tra*) at molar ratio from 1:1 to 1:2.5, both under CMV promoter regulation. Flow cytometry data were acquired 48 hours post-transfection. Data show the mean fluorescence normalized to its value at a plasmid molar ratio of 1. Error bars represent the standard error SE. ru: relative units. N=2 biological replicates. unpaired T-test. p-value: * < 0.05.

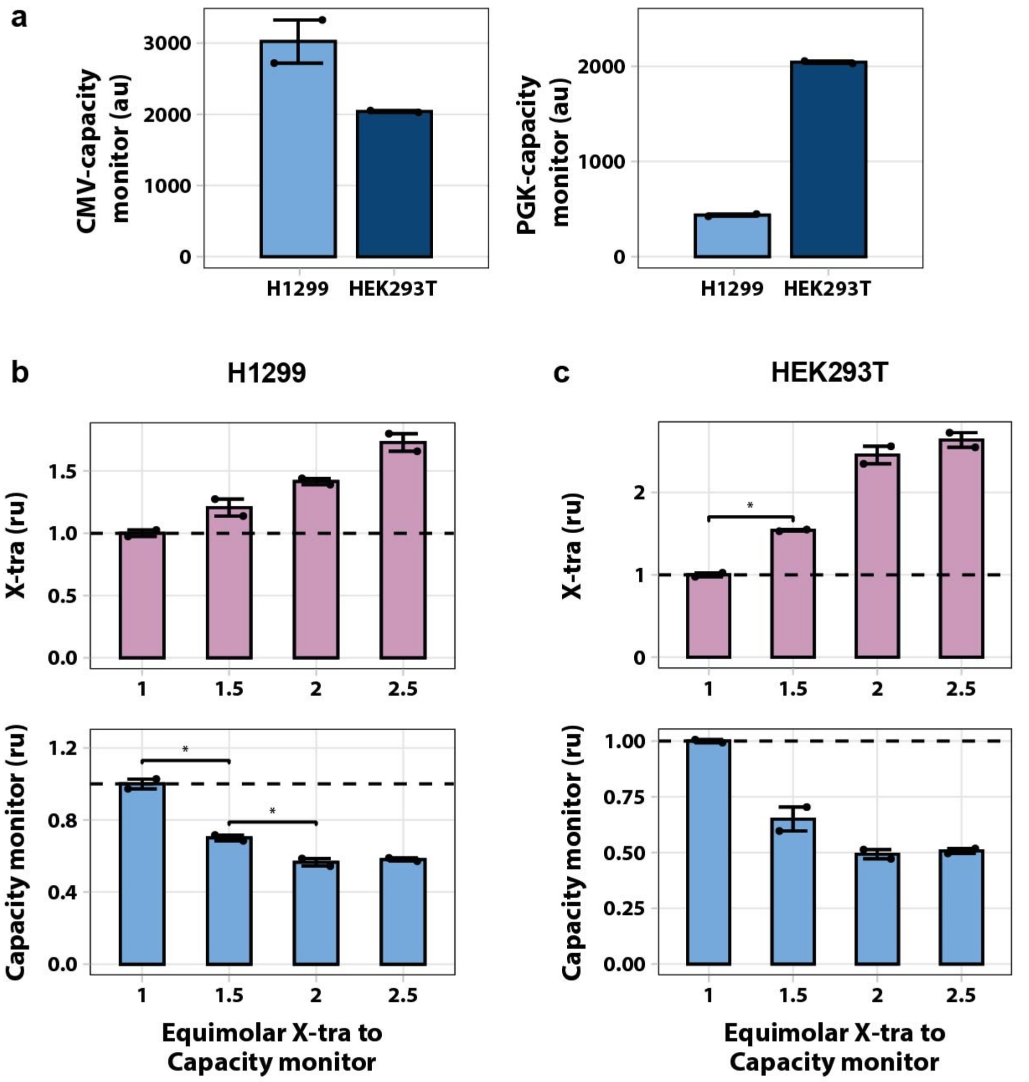

**Supplementary Figure 5. Relation of *X-tra* and *capacity monitor expression* in H1299 and HEK293T cell lines using PGK promoter. (a)** Levels of fluorescence driven by the same promoter (CMV or PGK) differ across cell lines. Data show absolute units of capacity monitor detected by flow cytometry in 1:1 molar ratio transfection. **(b)** Flow cytometry results of H1299 cells co-transfected with fixed amount of mKate (*capacity monitor*) under PGK promoter regulation and increasing amount of EGFP (*X-tra*) under CMV promoter regulation (molar ratio from 1:1 to 1:2.5). **(c)** Flow cytometry results of HEK293T cells co-transfected with fixed amount of mKate (*capacity monitor*) under PGK promoter regulation and increasing amount of EGFP (*X-tra*) under CMV promoter regulation. Data were acquired 48h post-transfection. In **(b)** and **(c)** data show the mean fluorescence normalized to its value at a plasmid molar ratio of 1. Error bars represent the standard error, SE. au: arbitrary units. ru: relative units. N=2 biological replicates. unpaired T-test. p-value: * < 0.05.

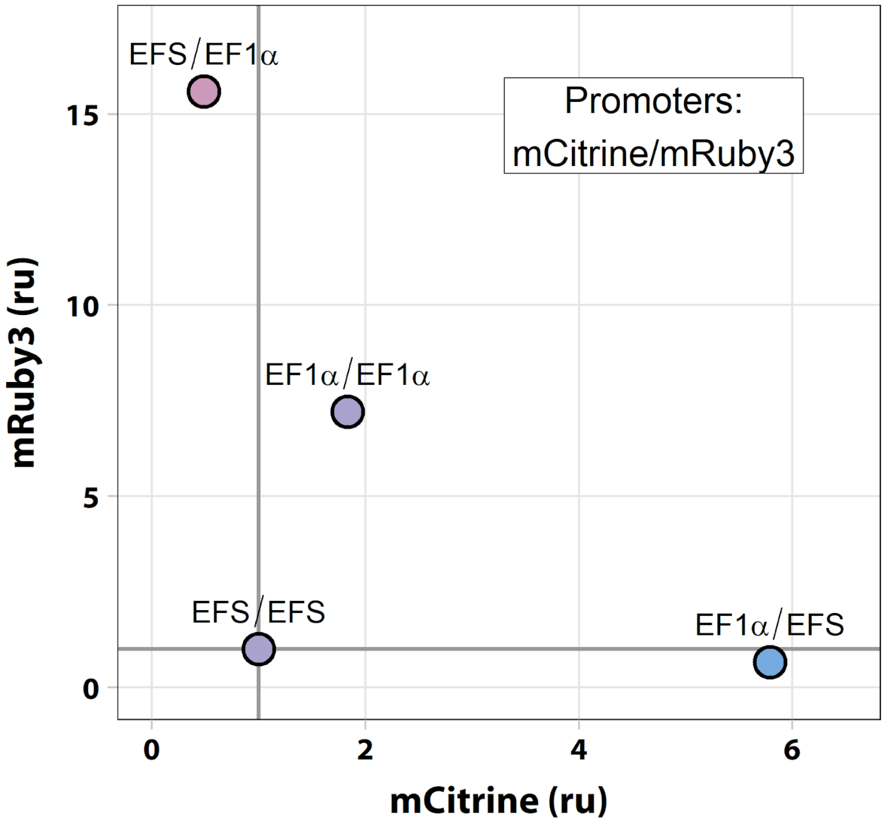

**Supplementary Figure 6. Promoter expression strength indirectly affects expression of co-transfected genes.** Plasmids expressing the fluorescent protein mCitrine and mRuby3 from a strong (EF-1α) or a medium strength EF-1α short (EFS) promoter were co-transfected in several molar ratio combinations. The expression levels for both mCitrine and mRuby3 were normalized by the data obtained from the weakest promoters pair (EFS/EFS). Similar to **Fig. 2a**, there is a negative correlation between the expression strength of one protein and the promoter strength of the other gene. Of note, when strong promoters drive both proteins, the global expression levels drop as already suggested by **Fig. 2a**. Data was acquired 48 hours after transfection and is plotted as fluorescence normalized to the EFS/EFS sample. ru: relative units.

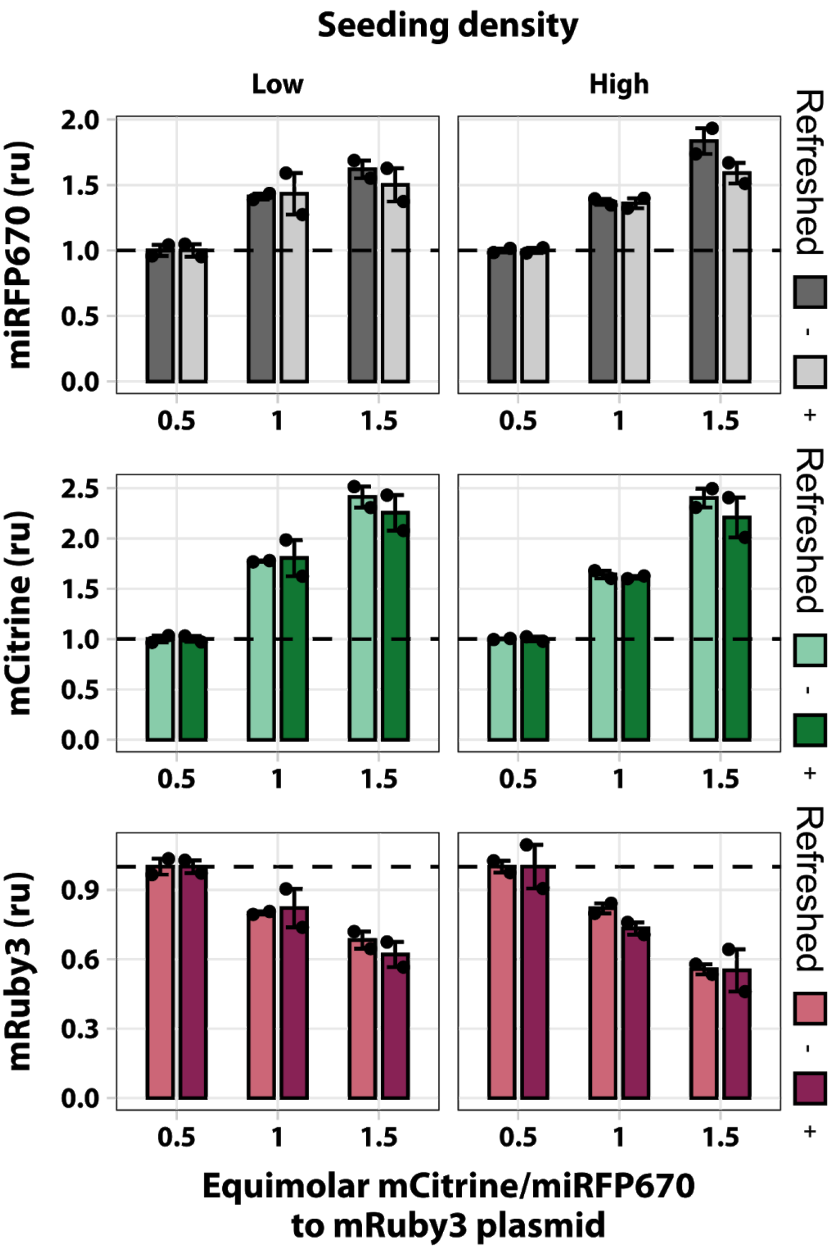

**Supplementary Figure 7. Assessing nutrient starvation and cell seeding density as potential impacts on limited resources.** In this experimental setting, cells were co-transfected with two plasmids. The first plasmid, which was provided at incremental levels, is composed of two transcriptional units (TU), one consisting of a strong promoter driving the expression of mCitrine (hEF1a), the other driving the expression of miRFP670 under a weak promoter (SV40). The second plasmid encodes for mRuby3 under a strong constitutive promoter (hEF1a).
HEK293T cells were seeded at $5e^4$ (low) and $7.5e^4$ (high) cells/well to assess seeding density effects. To investigate the effects of nutrient starvation, we refreshed the medium in two out of four wells per condition. Data were collected 48 hours post transfection and represent the mean fluorescence intensity of the three fluorescent proteins normalized to the 0.5 equimolar ratio condition. Error bars represent the standard error, SE. N=2 biological replicates.

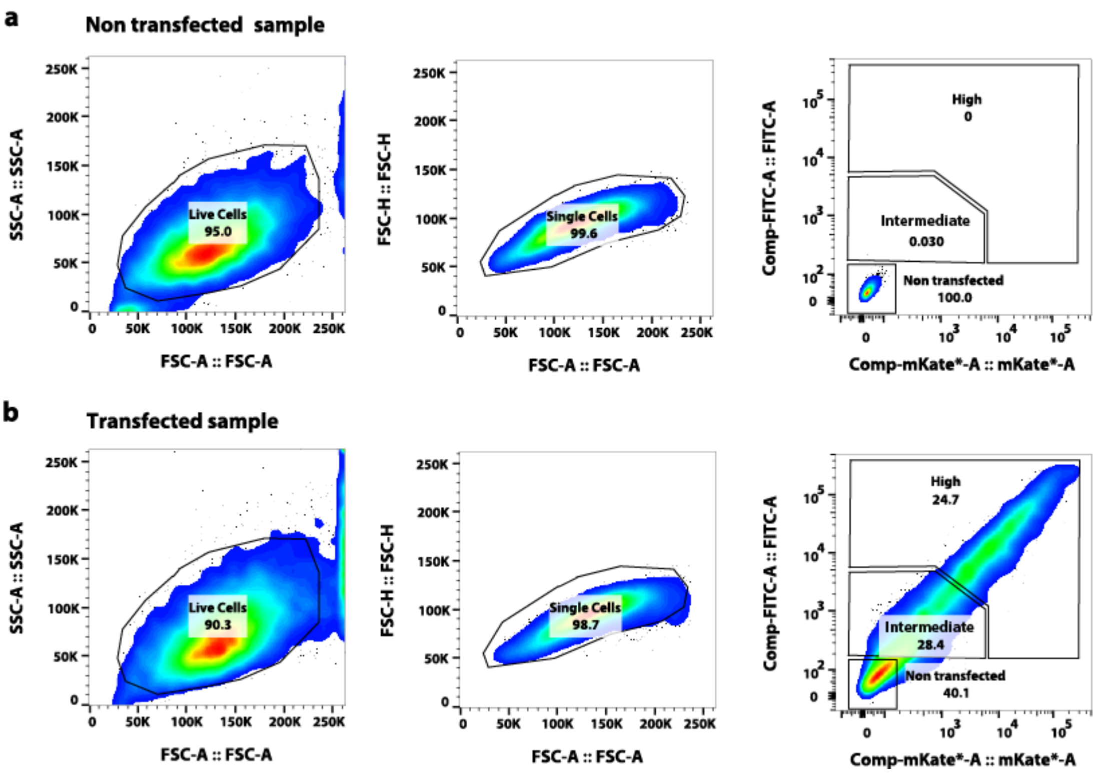

**Supplementary Figure 8. Sorting strategy.** H1299 cells were transfected with a plasmid encoding the fluorescent proteins EGFP and mKate, expressed from a bidirectional promoter. Cells were sorted by fluorescence intensity 48 hours post-transfection to collect *non-transfected*, *intermediate* and *high transfected* cells from the same transfection plate. **(a, b)** First, gates to select live and single cells were determined (left and middle plots). Then, the threshold for fluorescent intensity was set using a *non-transfected* sample as reference **(a, right)**. The two additional gates to collect *intermediate* and *high transfected* cells were created as shown in the plots **(a, b)** on the right.

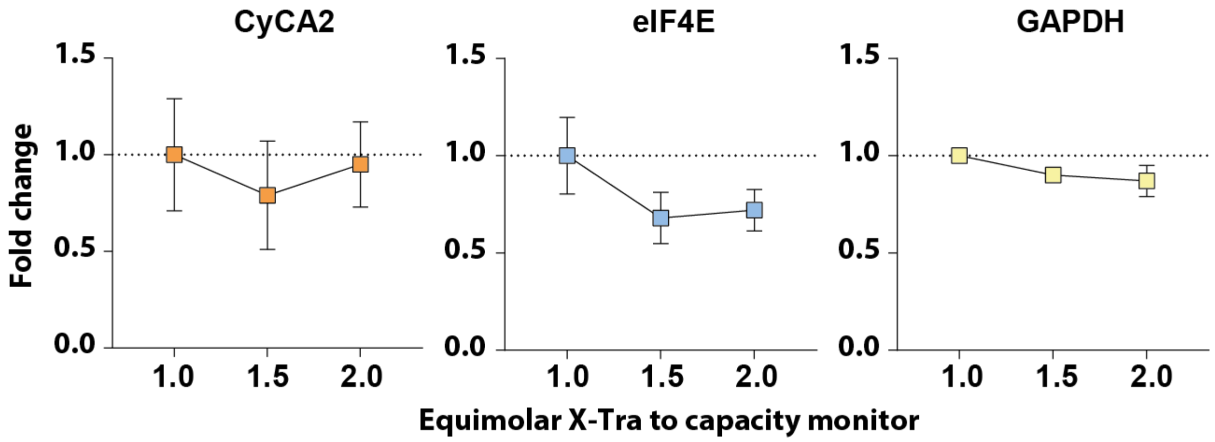

**Supplementary Figure 9. Effect of *X-tra* titration on endogenous genes.** We measured CyCA2, eIF4E and GAPDH mRNA levels by qPCR in the samples shown in **Fig. 2c** at 1.0, 1.5 and 2.0 molar ratios. Data are normalized to the equimolar ratio of 1.0. Error bars represent the standard error, SE. N=4 biological samples for CyCA2 and GAPDH. N=2 biological samples for eIF4E.

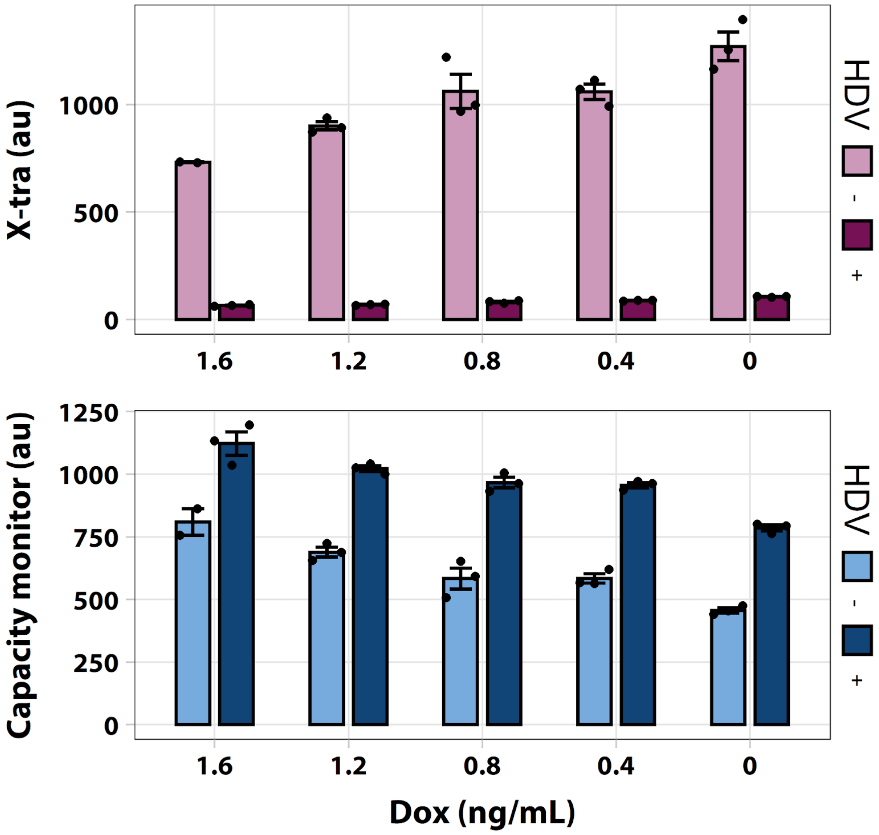

**Supplementary Figure 10. Fluorescence data of Fig. 2d shown in arbitrary units.** In this experimental settings Dox represses *X-tra* transcription. Thus, the lower Dox, the higher the *X-tra* levels, and as a consequence, the lower the *capacity monitor* levels. The HDV-dependent mRNA decapping and degradation of *X-tra* should consume less translational resources, which is consistent with the higher expression of the *capacity monitor* (dark blue bars) as compared to the inactive mutant (pale blue bars). Data was acquired 48 hours after transfection and is plotted as mean fluorescence intensity +/- SE. SE: standard error. N≥2 biological replicates.

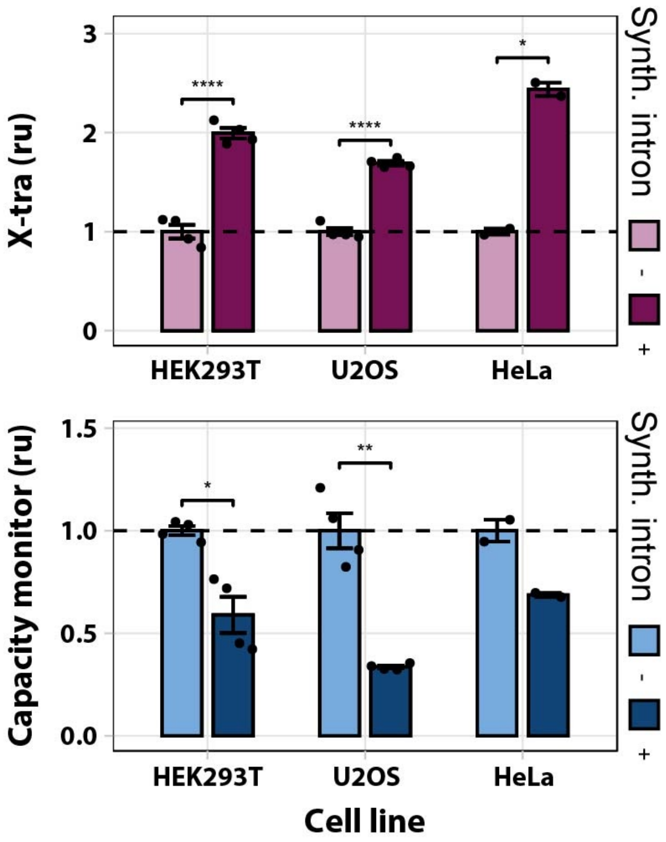

**Supplementary Figure 11. Translational resource limitation in different cell lines.** Flow cytometry results of HeLa, U2OS and HEK293T cells co-transfected with mKate (*X-tra*) with or without a synthetic intron, and EGFP (*capacity monitor*). Data show that when mKate expression is enhanced by the synthetic intron, EGFP levels decrease in all cell lines. Data were acquired 48 hours post-transfection and normalized to fluorescence values in the absence of the intron. Error bars represent the standard error, SE. ru: relative units. N≥2 biological replicates. unpaired T-test. p-value: **** < 0.0001, ** < 0.005,* < 0.05.

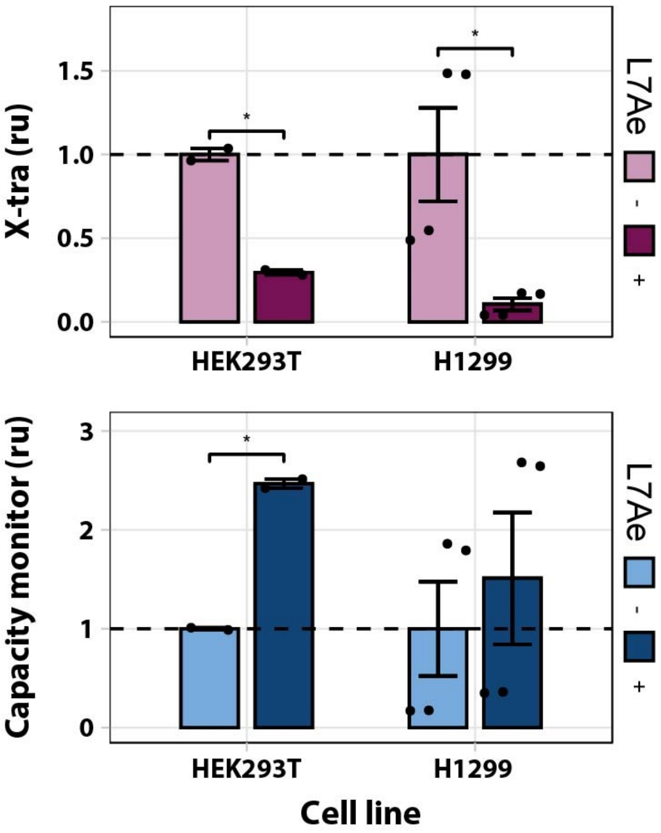

**Supplementary Figure 12. L7Ae-mediated resource re-allocation in mammalian cell lines. (a)** Flow cytometry results of HEK293T and H1299 cells co-transfected with 2kturn-EGFP (*X-tra*) and mKate (*capacity monitor*) in presence or absence of L7Ae. Data show that when *X-tra* is down-regulated, the *capacity monitor* levels increase. Data were acquired 48 hours post-transfection. Plots represent normalization of fluorescence values to the condition without L7Ae. Error bars represent the standard error, SE. ru: relative units. N≥2 biological replicates. unpaired T-test.p-value: * < 0.05.

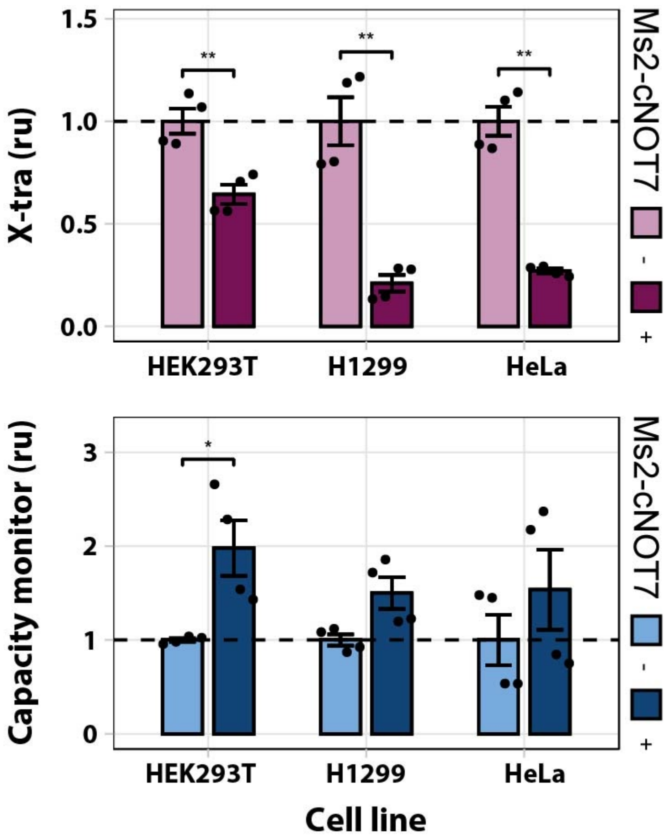

**Supplementary Figure 13. Ms2-cNOT7-mediated resource re-allocation in mammalian cell lines.** Flow cytometry results of HeLa, U2OS and HEK293T cells co-transfected with EGFP-8x-Ms2 target sites (*X-tra*) and mKate (*capacity monitor*) in presence and absence of Ms2-cNOT7. Data show that when *X-tra* is down-regulated, the *capacity monitor* levels increase. Data were acquired 48 hours post-transfection. Plots represent mean fluorescence intensity normalization to the condition without Ms2-cNOT7. Error bars represent the standard error. ru: relative units. N=4 biological replicates. unpaired T-test.p-value: ** < 0.01, * < 0.05.

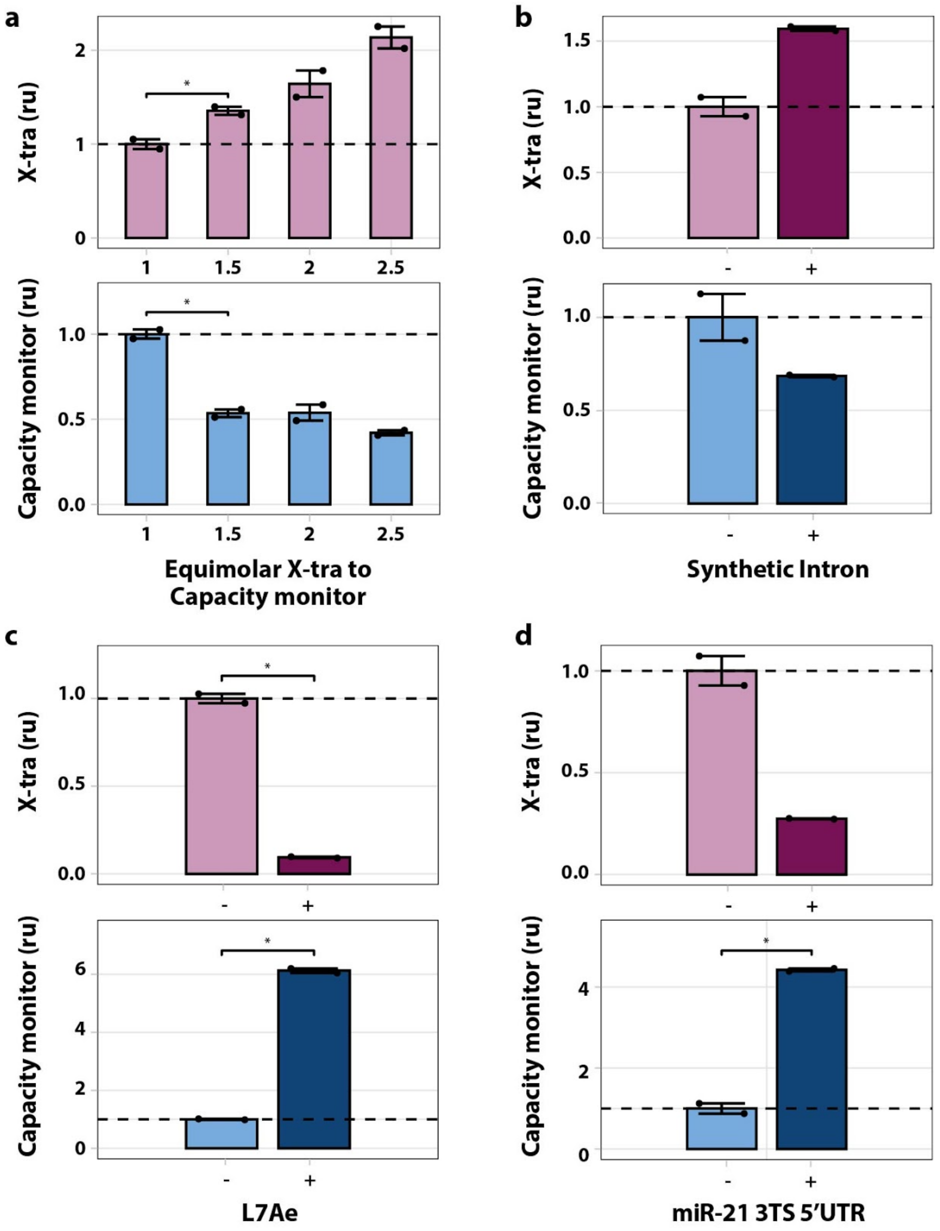

**Supplementary Figure 14. Gene expression burden in CHO-K1 cells. (a)** Flow cytometry results of CHO-K1 cells co-transfected with fixed amount of *capacity monitor* and increasing amount of *X-tra* (1:1 to 1:2.5 molar ratio), both under CMV promoter regulation. Data show the mean fluorescence normalized to its value at a plasmid molar ratio of 1. **(b)** Flow cytometry results of CHO-K1 cells co-transfected with *X-tra* (mKate) which includes or not a synthetic intron in the 5'UTR, and *capacity monitor* (EGFP). Data show that when mKate expression is enhanced by the synthetic intron, EGFP levels decrease. Data are normalized to fluorescence values in the absence of the intron. **(c)** Flow cytometry results of CHO-K1 cells co-transfected with 2kturn-EGFP (*X-tra*) and mKate (*capacity monitor*) in presence or absence of L7Ae. Data show that when *X-tra* is down-regulated, the *capacity monitor* levels increase. Plot represents normalization of fluorescence values to the condition without L7Ae. **(d)** Flow cytometry results of CHO-K1 cells co-transfected with mKate (*X-tra*) that includes or not miR-21 target sites in the 5'UTR, and EGFP (*capacity monitor*). Data show that *capacity monitor* levels are higher when the *X-tra* is downregulated by miR-21. Plot represents normalization of fluorescence values to the no target site condition. Data were acquired 48 hours post-transfection. Error bars represent the standard error. ru: relative units. N=2 biological replicates.

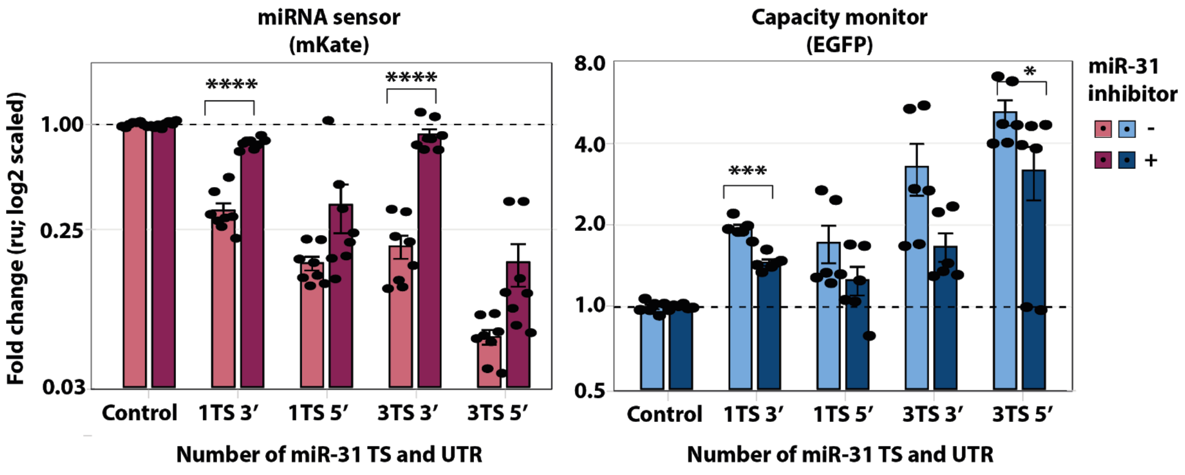

**Supplementary Figure 15. Inhibition of miR-31 in H1299 cells.** miR-31 activity impaired by a miR-31 inhibitor, leads to the rescue of *miRNA sensor* (mKate) expression in transfected H1299 cells. As a consequence, *capacity monitor* (EGFP) levels decrease. Both fluorescent proteins do not vary in the control. Data are expressed in logarithmic base 2 scale. Flow cytometry data were acquired 48 hours post-transfection and are plotted +/- SE. SE:standard error. ru: relative units. N=6 biological replicates, unpaired T-test. p-value: ****<0.0001, ***<0.0005, *<0.05.

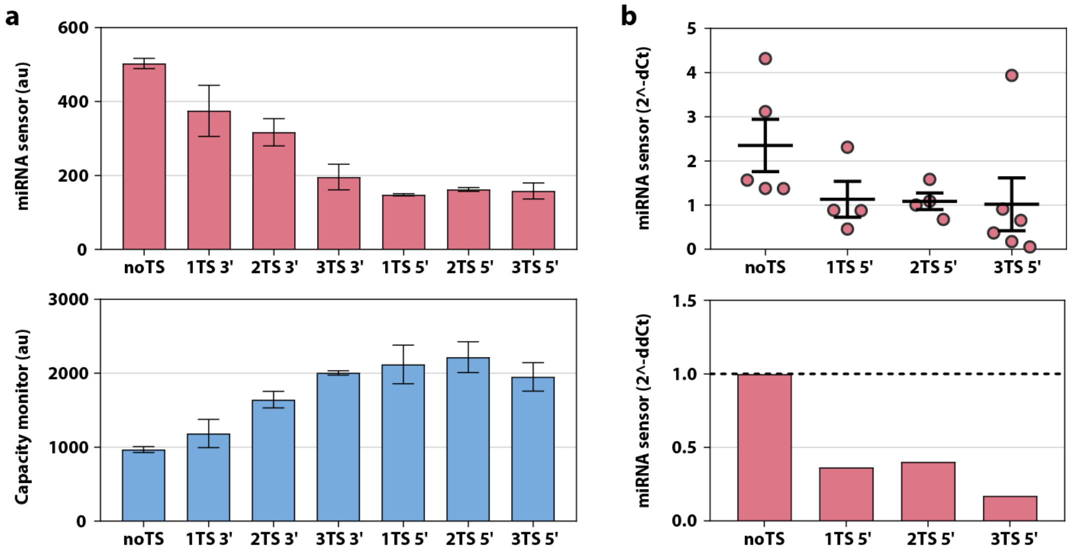

**Supplementary Figure 16. miR-31 sensor in H1299. (a)** Flow cytometry results of mKate-miR31-TS (*miRNA sensor*) co-transfected with EGFP (*capacity monitor*) in H1299 cells show that downregulation of *miRNA sensor* expression leads to an increase in *capacity monitor* levels. Data were acquired 48h post transfection +/- SE. SE:standard error. au: arbitrary units. N≥2 biological replicates. **(b)** qPCR measurement confirms lower mRNA levels of *miRNA sensor*. Top, scattered dot plot of 2^-dCt values. Bottom, bar plot of the fold change measured with the 2^-ddCt method[8].

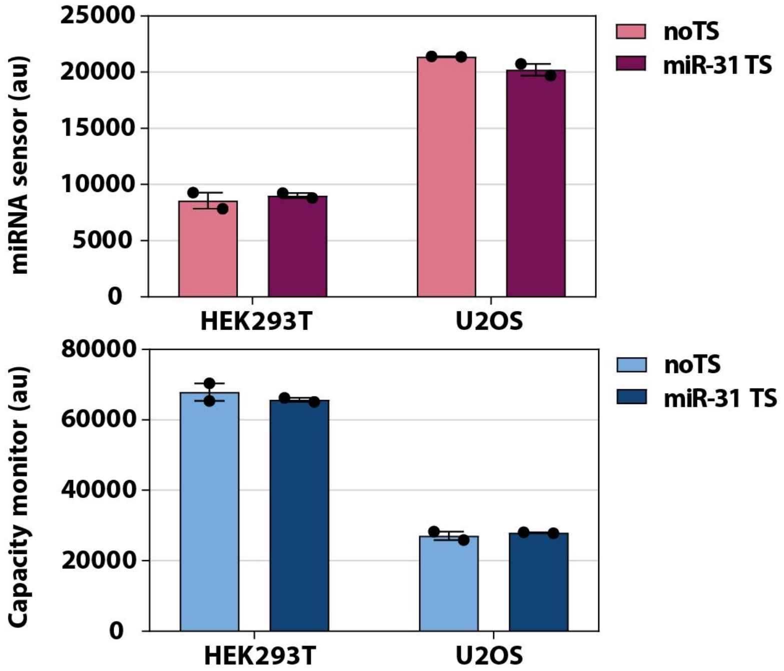

**Supplementary Figure 17. The increase of *capacity monitor* levels is a consequence of miRNA regulation.** U2OS and HEK293T cells were co-transfected with the 4-TS-3'UTR miR-31 sensor (*miRNA sensor*) and EGFP (*capacity monitor*). Both cell lines do not exhibit high expression of miR31, therefore *miRNA sensor* levels should not change. Data show that both *miRNA sensor* and the *capacity monitor* levels are comparable with and without miR-31 TS, indicating that the higher capacity monitor levels are indeed a consequence of miRNA activity. Data were acquired 48h post-transfection +/- SE. SE:standard error. au: arbitrary units. N=2 biological replicates.

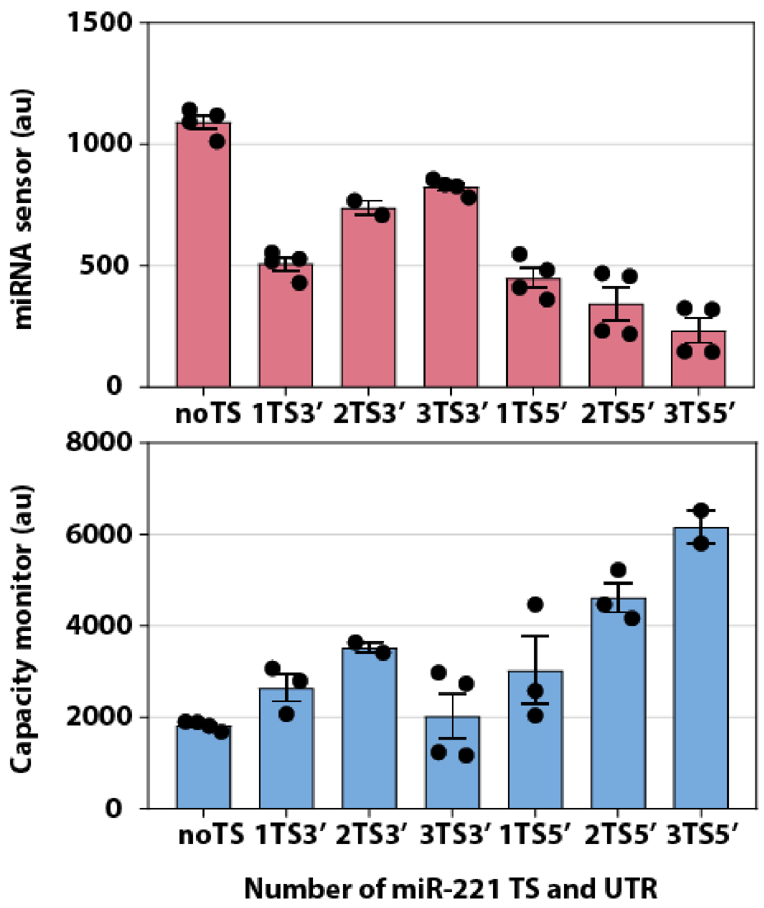

**Supplementary Figure 18. miRNA-mediated resource re-allocation in the U2OS cell line.** The *miRNA sensor* responds to miR-221, which is highly expressed in U2OS cells. Flow cytometry results of a co-transfection of mKate-miR221-TS (*miRNA sensor)* and EGFP (*capacity monitor*) in U2OS cells show the negative correlation of the two genes. Data were acquired 48h post-transfection and are plotted +/- SE. SE:standard error. au: arbitrary units. N≥2 biological replicates.

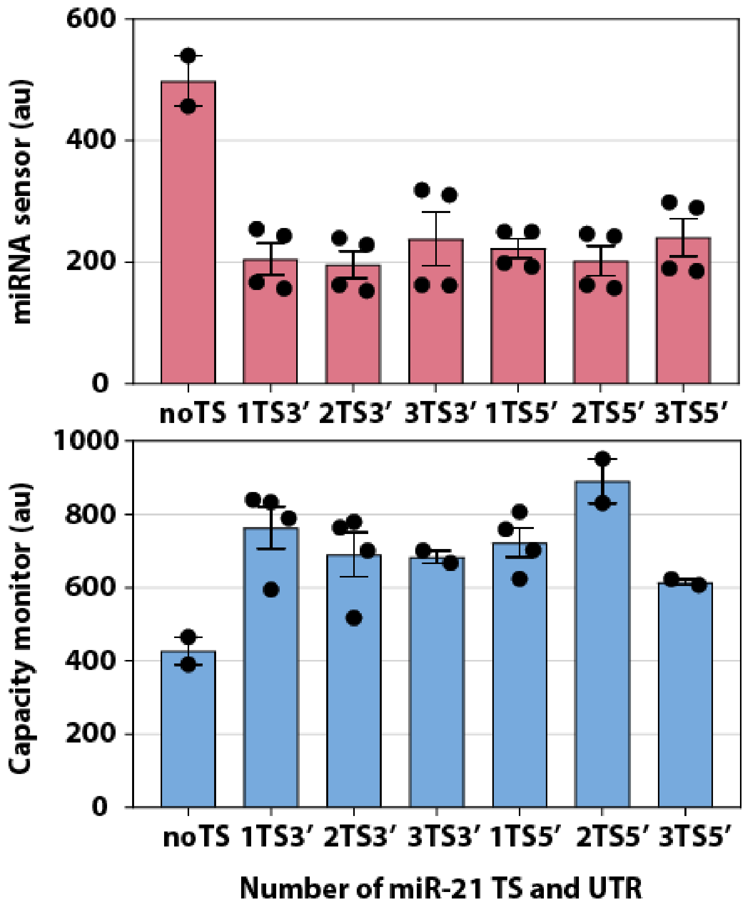

**Supplementary Figure 19. miRNA-mediated resource re-allocation in the HeLa cell line.** The *miRNA sensor* gene was designed for miR-21, which is highly expressed in HeLa cells. Flow cytometry results from a co-transfection of mKate-miR21-TS (*miRNA sensor)* and EGFP (*capacity monitor*) in HeLa cells. Interestingly, mKate downregulation seems to saturate already at 1TS3'. This may be due to the absolute levels of miR21 in this cell line. Data were acquired 48h post transfection and are plotted +/- SE. SE: standard error. au: arbitrary units. N≥2 biological replicates.

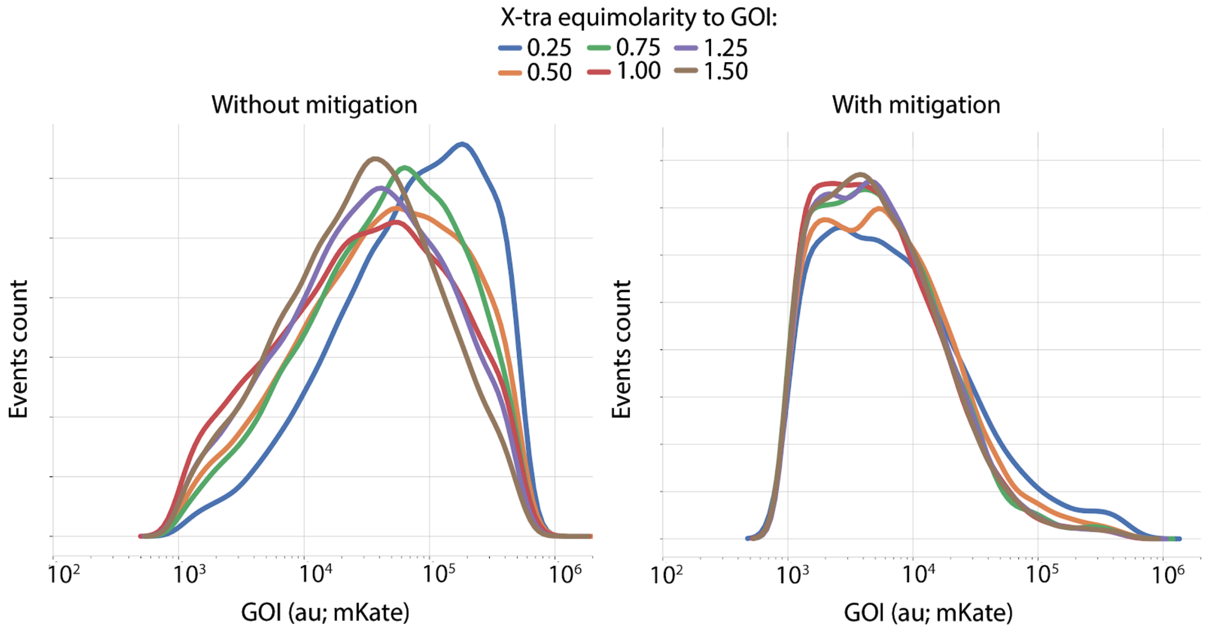

**Supplementary Figure 20. GOI fluorescence distribution in co-transfected cells population**. We compared the tolerance of mKate to increasing levels of *X-tra* gene in the absence or presence of an iFFL in which mKate includes miR-31 TS in the 5'UTR. The iFFL mitigation of resource competition is reflected by smaller shifts in GOI fluorescence at different equimolarities (right side). Data were acquired 48h post-transfection.

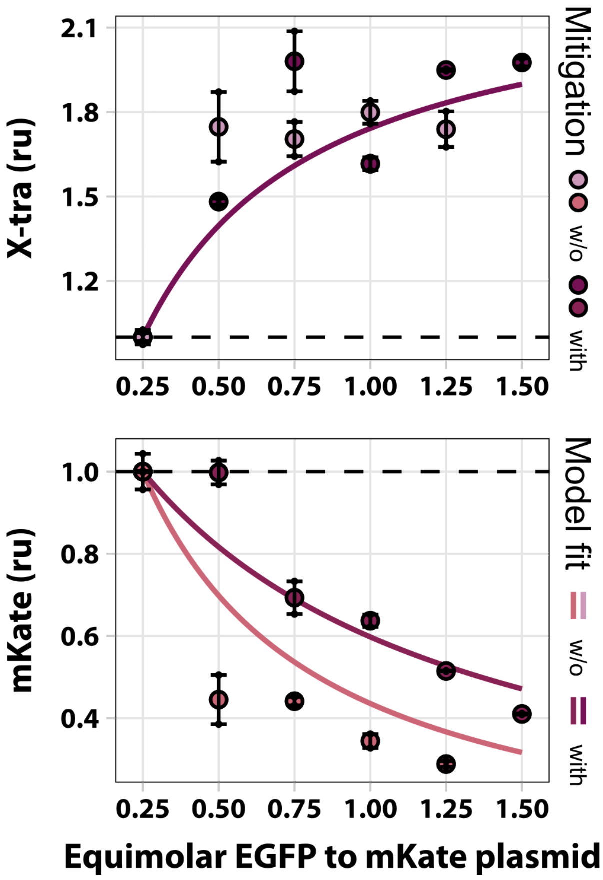

**Supplementary Figure 21. The miR-221-based iFFL improves tolerance to exogenous gene load in U2OS cells**. An iFFL whereby mKate includes miR-221 TS in the 5'UTR is less affected by the increased amount of the *X-tra* gene, as compared to the expression in the absence of miR-221 regulation. The model was unable to capture the differences in expression between the two conditions in the X-tra response due to the variability in the data. Therefore, the two lines plotted are exactly the same and it appears as if only one was plotted. Experimental data are normalized to the lowest equimolar ratio. The parameter values obtained by fitting are summarized in **Supplementary Table 10**. Data were acquired 48h post-transfection and are plotted +/- SE. SE:standard error. ru: relative units. N=2 biological replicates.

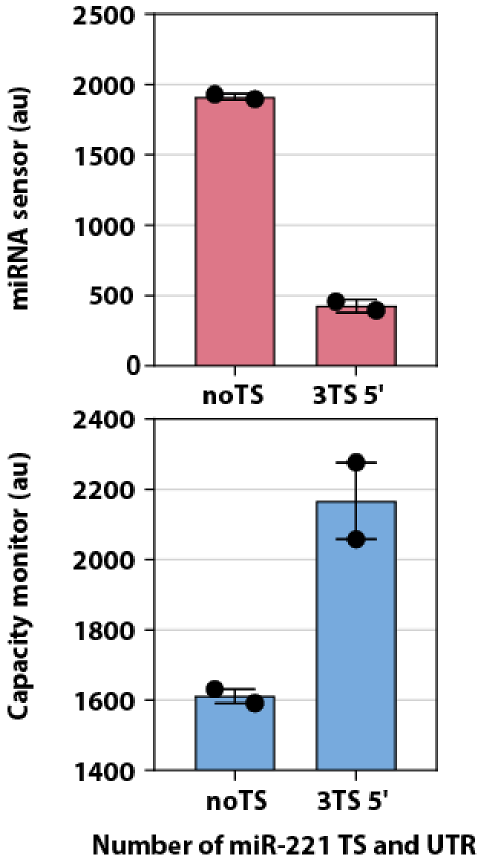

**Supplementary Figure 22. miRNA-mediated resource re-allocation in the HEK293T cell line.** The *miRNA sensor* gene was designed for miR-221, which is highly expressed in HEK293T cells. Flow cytometry results from a co-transfection of mKate-3xmiR221_5'UTR-TS (*miRNA sensor)* and EGFP (*capacity monitor*) in HEK293T cells. Data were acquired 48h post-transfection and are plotted +/- SE. SE: standard error. au: arbitrary units. N=2 biological replicates.

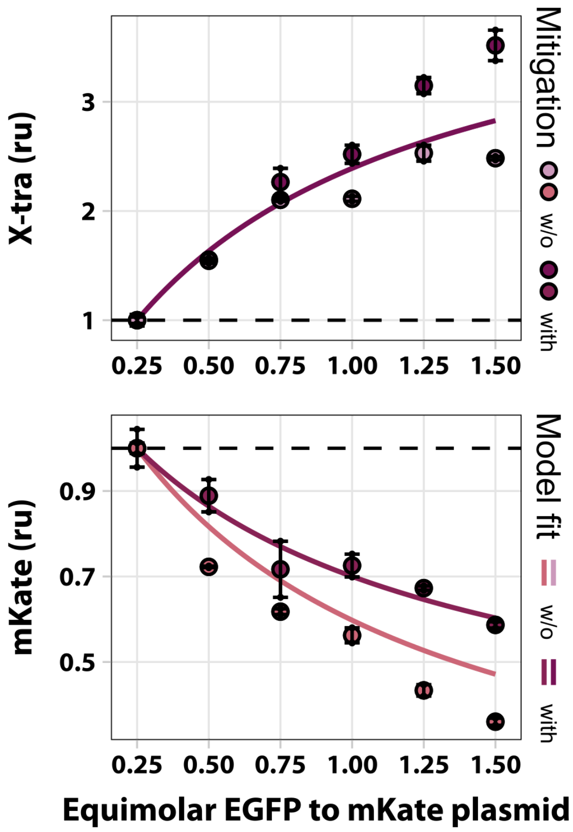

**Supplementary Figure 23. The miR-221-based iFFL improves tolerance to exogenous gene load in HEK293T cells**. We compared the tolerance of mKate to increasing levels of *x-tra* gene in the absence or presence of an iFFL whereby mKate includes miR-221 TS in the 5'UTR. The iFFL mitigates the effects of resource competition. The model was unable to capture the differences in expression between the two conditions in the X-tra response. Therefore, the two lines plotted are exactly the same and it appears as if only one was plotted. The parameter values obtained by fitting are summarized in **Supplementary Table 11**. Experimental data are normalized to the lowest equimolar ratio. Data were acquired 48h post-transfection and are plotted +/- SE. SE:standard error. ru: relative units. N=2 biological replicates.

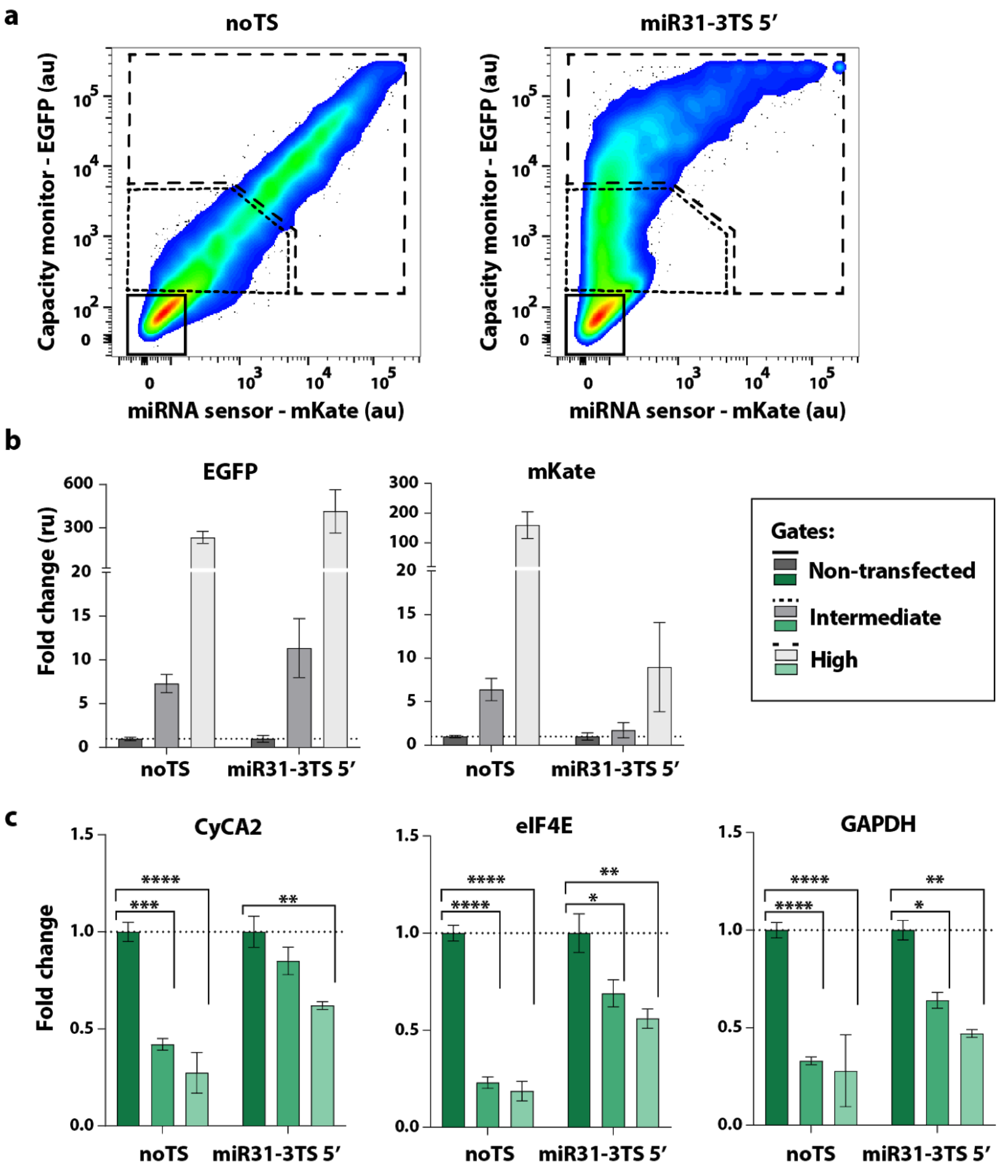

**Supplementary Figure 24. Impact of transient plasmid transfection on endogenous genes in noTS and miR31-sensor samples. (a)** H1299 cells were transfected with a bidirectional promoter plasmid encoding the fluorescent proteins EGFP (*capacity monitor)* and mKate (miRNA sensor), without **(noTS, left)** or with TS for miR-31 **(miR31-sensor, right)**. Cells were sorted by fluorescence intensity 48 hours after transfection to collect *non-transfected*, *intermediate transfected* and *high transfected* cells from the same transfection plate. **(b)** Protein levels of EGFP and mKate in sorted populations of noTS and miR31-sensor transfected cells. Consistent with the gates, fluorescence intensity increases in *intermediate* and *high transfected* cells when compared to *non-transfected* cells. In agreement with data shown in **Fig 2h and 3b,c**, EGFP fluorescence is higher in miR31-sensor samples, while mKate is lower. Data are normalized to the fluorescence in the *non-transfected* population. **(c)** mRNA levels of CyCA2, eIF4E and GAPDH in sorted samples. All three endogenous genes decrease in *intermediate* and *high transfected* cells as compared to *non-transfected* cells. However, in cells transfected with the miR31-sensor circuit the decrease of expression is lower. mRNA levels are normalized to the *non-transfected* population. Data were collected 48 hours after transfection and are represented as mean +/- SE. SE: standard error. ru: relative units. unpaired T-test. p-value: ****<0.0001, ***<0.0005, **<0.005, *<0.05. N≥3 biological replicates.

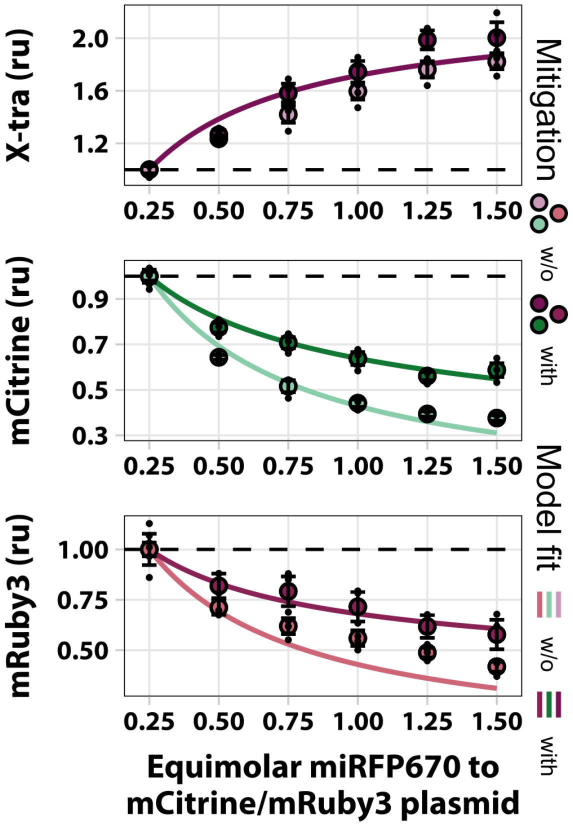

**Supplementary Figure 25. The iFFL architecture improves tolerance to increase gene load in a 3-output system.** Mouse embryonic stem cells were transfected with the miRNA mitigation iFFL shown in **Fig. 5d**. Light and dark colors represent gene expression levels in the absence or presence of mitigation. The solid lines show a model that includes resources, fit to the experimental data. Experimental data are normalized to the lowest equimolar ratio. The parameter values obtained by fitting are summarized in **Supplementary Table 12**. All data were acquired 48h post transfection and are plotted +/- SE. SE: standard error. ru: relative units. N=3 biological replicates.

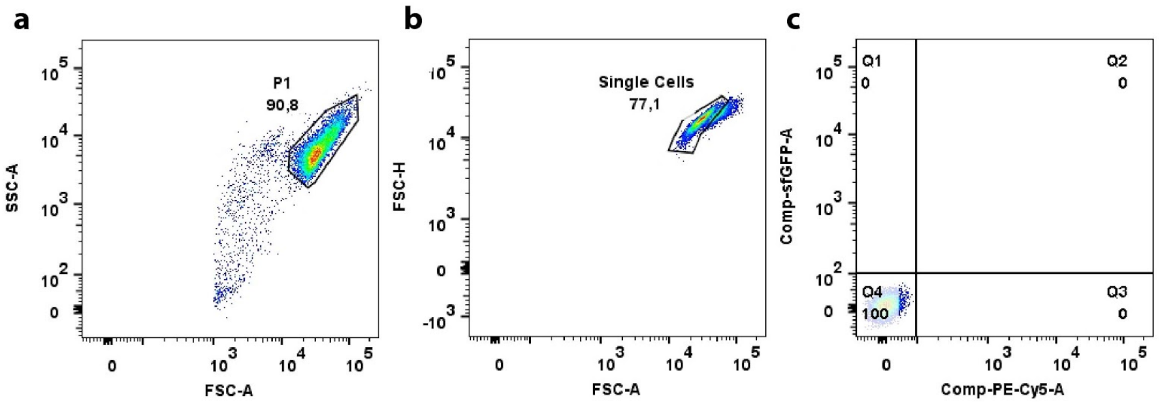

**Supplementary Figure 26. FACS gating strategy. (a)** The recorded events were gated in the FSC-A vs SSC-A channels to select the living cells population (P1). **(b)** The P1 was then gated in the FSC-A vs FSC-H channels to select the single cell population. **(c)** For each experiment a sample of non-transfected cells was used to set the positive threshold for each fluorescence. Cells selected following this pipeline were then analyzed.

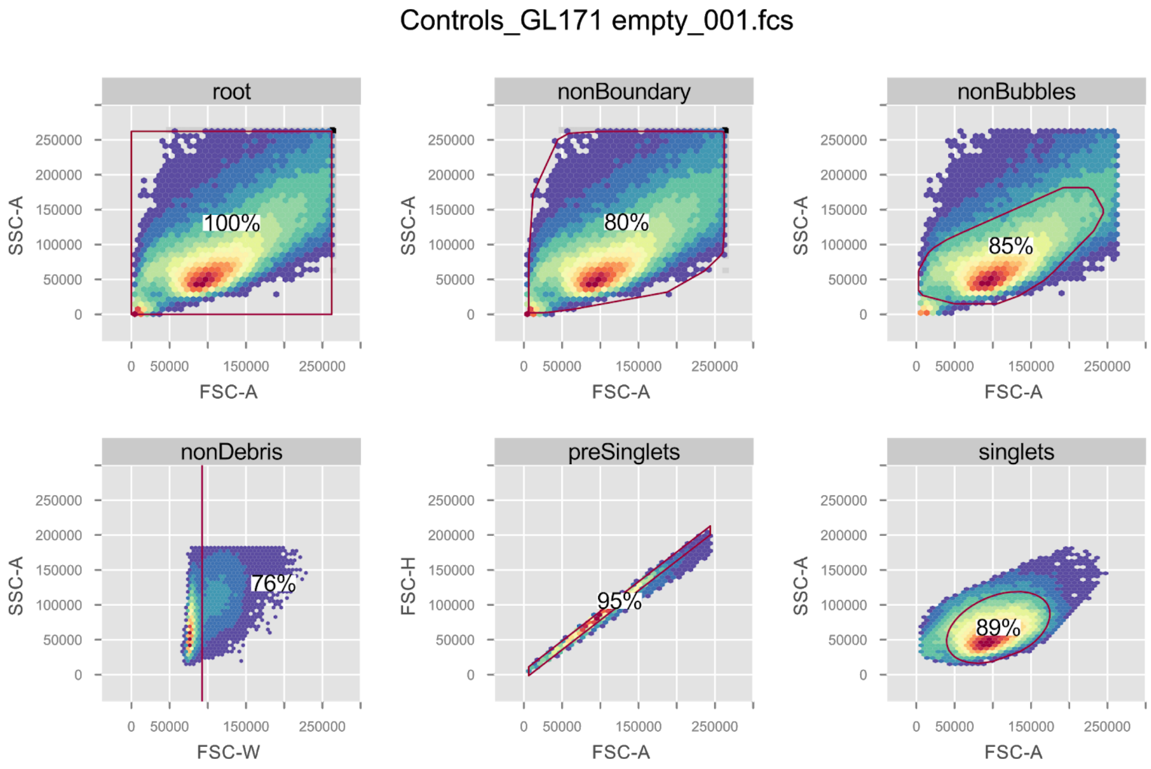

**Supplementary Figure 27. Alternative FACS gating strategy.** The plots show the hierarchical gating strategy implemented in a custom R script. The hierarchy progresses from left to right, top to bottom. **Top left:** The first gate removes events that potentially lie on the boundary of the detectable values. **Top middle:** This gate facilitates the subsequent gating by removing potential bubbles that were recorded. **Top right:** Here, a custom density-based gating strategy is employed to select for the living cell population and remove debris. **Bottom left:** In this gate the tail of the distribution in the FSC-W channel representing the bulk of the doublet event is removed. **Bottom middle:** The singlet population is further refined by gating in the FSC-H vs FSC-A channels. **Bottom right:** The resulting singlets are further refined by applying an ellipse gate around the point of highest density.

**Supplementary Note 1.** Simple model of plasmid copy number induced competition for limited resources.

To derive the equations used in Figure 1A of the main text the following model was used:

$$\dot{M}_i = k_{M_i}^{eff}(G_1, \ldots, G_n)G_i - \delta_{M_i} M_i$$

$$\dot{P}_i = k_{P_i}^{eff}(M_1, \ldots, M_n)M_i - \delta_{P_i} P_i$$

Here, we distinguish between two potential pools of shared limited resources. One for mRNA production and one for protein production. The mRNA species are denoted by the subscript letter $M$ and the protein species are denoted by the subscript letter $P$. The copy number of the genes are denoted by $G_i$ and are assumed to be constant. The degradation rates of each species is represented by a $\delta$ subscripted with the respective species' name. Using the definition

$$k_{B_i}^{eff}(A_1, \ldots, A_n) := k_{B_i}^{cat} R^{total} \frac{k_{m_{B_i}}^{-1}}{1 + \sum_{j=1}^{n} k_{m_{B_j}}^{-1} A_j}$$

the equations can be written as

$$\dot{M}_i = k_{M_i}^{cat} R_M^{total} \frac{k_{m_{M_i}}^{-1}}{1 + \sum_{j=1}^{n} k_{m_{M_j}}^{-1} G_j} G_i - \delta_{M_i} M_i$$

$$\dot{P}_i = k_{P_i}^{cat} R_P^{total} \frac{k_{m_{P_i}}^{-1}}{1 + \sum_{j=1}^{n} k_{m_{P_j}}^{-1} M_j} M_i - \delta_{P_i} P_i$$

We solve for the steady state expressions of each of the species by setting the left-hand side of the equations to zero. After simplifying we obtain the following expressions:

$$M_i^* = \frac{k_{M_i}^{cat} R_M^{total}}{\delta_{M_i}} \frac{k_{m_{M_i}}^{-1} G_i}{1 + \sum_{j=1}^{n} k_{m_{M_j}}^{-1} G_j}$$

$$P_i^* = \frac{k_{M_i}^{cat} R_M^{total}}{\delta_{M_i}} \frac{k_{P_i}^{cat} R_P^{total}}{\delta_{P_i}} \frac{k_{m_{M_i}}^{-1} k_{m_{P_i}}^{-1} G_i}{1 + \sum_{j=1}^{n} k_{m_{M_j}}^{-1} \left(1 + k_{m_{P_j}}^{-1} \frac{k_{M_j}^{cat} R_M^{total}}{\delta_{M_j}}\right) G_j}$$

The normalized expressions as shown in Figure 2A are further defined as $\hat{P}_i^* = \frac{P_i^*}{P_i^*|_{G_k = G_{k_0}}}$ with $k \neq i$, which gives

$$\hat{P}_i^* = \frac{1 + k_{m_{M_k}}^{-1} \left(1 + k_{m_{P_k}}^{-1} \frac{k_{M_k}^{cat} R_M^{total}}{\delta_{M_k}}\right) G_{k_0} + \sum_{j \neq k}^{n} k_{m_{M_j}}^{-1} \left(1 + k_{m_{P_j}}^{-1} \frac{k_{M_j}^{cat} R_M^{total}}{\delta_{M_j}}\right) G_j}{1 + k_{m_{M_k}}^{-1} \left(1 + k_{m_{P_k}}^{-1} \frac{k_{M_k}^{cat} R_M^{total}}{\delta_{M_k}}\right) G_k + \sum_{j \neq k}^{n} k_{m_{M_j}}^{-1} \left(1 + k_{m_{P_j}}^{-1} \frac{k_{M_j}^{cat} R_M^{total}}{\delta_{M_j}}\right) G_j}$$

Specifically for Figure 2A $n$ was set to 2.

**Supplementary Note 2.** Normalized gene expression of low absolute expression levels are more sensitive to reduced availability of resources.

To show that normalized expression is more sensitive to burden at low expression levels we take the general term for the protein levels derived in Supplementary Note 1.

$$P_i^* = \frac{k_{M_i}^{cat} R_M^{total}}{\delta_{M_i}} \frac{k_{P_i}^{cat} R_P^{total}}{\delta_{P_i}} \frac{k_{m_{M_i}}^{-1} k_{m_{P_i}}^{-1} G_i}{1 + \sum_{j=1}^{n} k_{m_{M_j}}^{-1} \left(1 + k_{m_{P_j}}^{-1} \frac{k_{M_j}^{cat} R_M^{total}}{\delta_{M_j}}\right) G_j}$$

To simplify the above term we lump parameters by setting

$$\beta_{M_i} := k_{m_{P_i}}^{-1} \frac{k_{M_i}^{cat} R_M^{total}}{\delta_{M_i}}$$

$$\alpha_{P_i} := \frac{k_{P_i}^{cat} R_P^{total}}{\delta_{P_i}}$$

Which gives

$$P_i^* = \frac{\alpha_{P_i} \beta_{M_i} k_{m_{M_i}}^{-1} G_i}{1 + \sum_{j=1}^{n} k_{m_{M_j}}^{-1} \left(1 + \beta_{M_j}\right) G_j}$$

From this term we can write the expression level normalized with respect to the expression level at $G_{k0}$ as

$$\hat{P}_i^* = \frac{P_i^*}{P_i^*|_{G_k = G_{k0}}} = \frac{1 + k_{m_{M_k}}^{-1} \left(1 + \beta_{M_k}\right) G_{k0} + \sum_{j \neq k}^{n} k_{m_{M_j}}^{-1} \left(1 + \beta_{M_j}\right) G_j}{1 + k_{m_{M_k}}^{-1} \left(1 + \beta_{M_k}\right) G_k + \sum_{j \neq k}^{n} k_{m_{M_j}}^{-1} \left(1 + \beta_{M_j}\right) G_j}$$

with $k \neq i$.

To test whether the normalized expression for a gene expressing at low levels is less sensitive to reduced resource availability than its high expressing counterpart we wish to evaluate the following inequality $\hat{P}_i^* < \hat{P}_i^*|_{G_i = \bar{G}_i}$ where the low expression is given by the assumption that $\bar{G}_i < G_i$. To simplify this inequality we write it as the equivalent inequality given by $\mathrm{sgn}(\hat{P}_i^* - \hat{P}_i^*|_{G_i = \bar{G}_i}) < 0$ where $\mathrm{sgn}$ denotes the sign function. Plugging in the expression for $\hat{P}_i^*$ and $\hat{P}_i^*|_{G_i = \bar{G}_i}$ gives

$$\mathrm{sgn}\left( \frac{1 + k_{m_{M_k}}^{-1} \left(1 + \beta_{M_k}\right) G_{k_0} + \sum_{j \neq k}^{n} k_{m_{M_j}}^{-1} \left(1 + \beta_{M_j}\right) G_j}{1 + k_{m_{M_k}}^{-1} \left(1 + \beta_{M_k}\right) G_k + \sum_{j \neq k}^{n} k_{m_{M_j}}^{-1} \left(1 + \beta_{M_j}\right) G_j} - \frac{1 + k_{m_{M_i}}^{-1} \left(1 + \beta_{M_i}\right) \bar{G}_i + k_{m_{M_k}}^{-1} \left(1 + \beta_{M_k}\right) G_{k_0} + \sum_{j \neq \{i,k\}}^{n} k_{m_{M_j}}^{-1} \left(1 + \beta_{M_j}\right) G_j}{1 + k_{m_{M_i}}^{-1} \left(1 + \beta_{M_i}\right) \bar{G}_i + k_{m_{M_k}}^{-1} \left(1 + \beta_{M_k}\right) G_k + \sum_{j \neq \{i,k\}}^{n} k_{m_{M_j}}^{-1} \left(1 + \beta_{M_j}\right) G_j} \right) < 0$$

Given that all parameters are positive and we demand that $G_k > G_{k0}$ the term on the left hand side of the inequality above reduces to

$$\mathrm{sgn}(G_i - \bar{G}_i) < 0$$

Given our initial assumption that $\bar{G}_i < G_i$ and the requirement that both are positive real numbers we find that the claim stated in the inequality is false because the sign function evaluates to 1.

Therefore, the model shows that the normalized expression of a gene expressing at low absolute levels will be more affected by resource availability than its high expressing counterpart.

**Supplementary Note 3.** Model for the topologies from Lillacci et al. [1].

The model for the four topologies can be given by a system of ordinary differential equations, where setting the individual repression rates $\eta_{M_1}$ and/or $\eta_{M_2}$ of the microRNA to zero specifies the topology. More specifically, $\eta_{M_1} = 0$ and $\eta_{M_2} = 0$ is the open-loop (OLP) topology, $\eta_{M_1} = 0$ and $\eta_{M_2} > 0$ the incoherent feedforward (IFF) topology, $\eta_{M_1} > 0$ and $\eta_{M_2} = 0$ is the feedback (FBK) topology and $\eta_{M_1} > 0$ and $\eta_{M_2} > 0$ is the hybrid (HYB) topology.

$$\dot{M}_1 = k_{M_1}^{eff}(G_1, G_2 f(P_1), G_3) G_1 - \left(\delta_{M_1} + \eta_{M_1} m\right) M_1$$

$$\dot{M}_2 = k_{M_2}^{eff}(G_1, G_2 f(P_1), G_3) G_2 f(P_1) - \left(\delta_{M_2} + \eta_{M_2} m\right) M_2$$

$$\dot{M}_3 = k_{M_3}^{eff}(G_1, G_2 f(P_1), G_3) G_3 - \delta_{M_3} M_3$$

$$\dot{m} = k_m^{eff}(G_1, G_2 f(P_1), G_3) G_2 f(P_1) - \delta_m m$$

$$\dot{P}_1 = k_{P_1}^{eff}(M_1, M_2, M_3) M_1 - \delta_{P_1} P_1$$

$$\dot{P}_2 = k_{P_2}^{eff}(M_1, M_2, M_3) M_2 - \delta_{P_2} P_2$$

$$\dot{P}_1 = k_{P_3}^{eff}(M_1, M_2, M_3) M_3 - \delta_{P_3} P_3$$

In this system of equations the species $M_1$, $M_2$ and $M_3$ correspond to tTA-Cer. mRNA, DsRed mRNA and mCitr. mRNA respectively as shown in Figure 4B and C. $P_1$ denotes the transcriptional activator tTA-Cer., $P_2$ denotes the fluorescent protein DsRed and $P_3$ denotes the fluorescent protein mCitrine. $m$ denotes miR-FF4 expressed from the same gene as DsRed. Furthermore, $G_1$, $G_2$ and $G_3$ correspond to the plasmid copy number of tTA-Cer., DsRed and mCitr. Respectively. The rates beginning with a $\delta$ denote the degradation rates of the species written in the subscript. The rates beginning with $\eta$ correspond to the repression rates of the microRNA FF4. Lastly, the transcriptional activation was modeled by a hill-type function $f(x) := \frac{x^h}{\kappa^h + x^h}$. The steady states for the protein species used for fitting can be written as

$$P_1^* = \frac{\alpha_{P_1^*} \frac{\beta_{M_1} G_1 k_{m_{M_1}}^{-1}}{1+G_1 k_{m_{M_1}}^{-1}+\gamma_{M_2} f(P_1^*)\left(1+\theta_{M_1}\right)+\gamma_{M_3}}}{1 + \frac{\beta_{M_1} G_1 k_{m_{M_1}}^{-1}}{1+G_1 k_{m_{M_1}}^{-1}+\gamma_{M_2} f(P_1^*)\left(1+\theta_{M_1}\right)+\gamma_{M_3}} + \frac{\beta_{M_2}\gamma_{M_2} f(P_1^*)}{1+G_1 k_{m_{M_1}}^{-1}+\gamma_{M_2} f(P_1^*)\left(1+\theta_{M_2}\right)+\gamma_{M_3}} + \frac{\beta_{M_3}\gamma_{M_3}}{1+G_1 k_{m_{M_1}}^{-1}+\gamma_{M_2} f(P_1^*)+\gamma_{M_3}}}$$

$$P_2^* = \frac{\alpha_{P_2} \frac{\beta_{M_2}\gamma_{M_2} f(P_1^*)}{1+G_1 k_{m_{M_1}}^{-1}+\gamma_{M_2} f(P_1^*)\left(1+\theta_{M_2}\right)+\gamma_{M_3}}}{1 + \frac{\beta_{M_1} G_1 k_{m_{M_1}}^{-1}}{1+G_1 k_{m_{M_1}}^{-1}+\gamma_{M_2} f(P_1^*)\left(1+\theta_{M_1}\right)+\gamma_{M_3}} + \frac{\beta_{M_2}\gamma_{M_2} f(P_1^*)}{1+G_1 k_{m_{M_1}}^{-1}+\gamma_{M_2} f(P_1^*)\left(1+\theta_{M_2}\right)+\gamma_{M_3}} + \frac{\beta_{M_3}\gamma_{M_3}}{1+G_1 k_{m_{M_1}}^{-1}+\gamma_{M_2} f(P_1^*)+\gamma_{M_3}}}$$

$$P_3^* = \frac{\alpha_{P_3} \frac{\beta_{M_3}\gamma_{M_3}}{1+G_1 k_{m_{M_1}}^{-1}+\gamma_{M_2} f(P_1^*)+\gamma_{M_3}}}{1 + \frac{\beta_{M_1} G_1 k_{m_{M_1}}^{-1}}{1+G_1 k_{m_{M_1}}^{-1}+\gamma_{M_2} f(P_1^*)\left(1+\theta_{M_1}\right)+\gamma_{M_3}} + \frac{\beta_{M_2}\gamma_{M_2} f(P_1^*)}{1+G_1 k_{m_{M_1}}^{-1}+\gamma_{M_2} f(P_1^*)\left(1+\theta_{M_2}\right)+\gamma_{M_3}} + \frac{\beta_{M_3}\gamma_{M_3}}{1+G_1 k_{m_{M_1}}^{-1}+\gamma_{M_2} f(P_1^*)+\gamma_{M_3}}}$$

Here, $\alpha_{P_i}$ and $\beta_{M_i}$ are the same as defined in Supplementary Note 2. Additionally, $\gamma_{M_i} := k_{m_{M_i}}^{-1} G_i$ for $i \in \{2,3\}$ and $\theta_{M_i} := \frac{\eta_{M_i}}{\delta_{M_i}} \frac{k_{M_2}^{cat} R_M^{total}}{\delta_m}$ were introduced. The equations were fit in the implicit form shown because a closed form solution could not be obtained.

**Supplementary Note 4.** Models for endogenous microRNA-based iFFL and synthetic microRNA-based iFFL circuits.

**Endogenous microRNA-based iFFL:**

The system of equations used to derive the steady state expressions is given by

$$\dot{M}_1 = k_{M_1}^{eff}(G_1, G_2, G_m)G_1 - \delta_{M_1}M_1$$

$$\dot{M}_2 = k_{M_2}^{eff}(G_1, G_2, G_m)G_2 - (\delta_{M_2} + \eta_{M_2}m)\, M_2$$

$$\dot{m} = k_{m}^{eff}(G_1, G_2, G_m)G_m - \delta_m m$$

$$\dot{P}_1 = k_{P_1}^{eff}(M_1, M_2)M_1 - \delta_{P_1}P_1$$

$$\dot{P}_2 = k_{P_2}^{eff}(M_1, M_2)M_2 - \delta_{P_2}P_2$$

Here, $M_1$ and $M_2$ represent the mRNA species for the fluorescent proteins EGFP (*X-tra*) and mKate (*GOI*) respectively. $P_1$ and $P_2$ correspond to the proteins themselves and $m$ denotes the microRNA miR-31. As in Supplementary Note 1 and 3, the degradation rates are shown as $\delta$ subscripted with the species they correspond to and the repression rates are shown as $\eta$ subscripted with the respective species. $G_1$ and $G_2$ are the plasmid copy number of the EGFP and mKate plasmids respectively and $G_m$ is the copy number of the microRNA on the genome. The steady states for the protein species can be written as

$$P_1^* = \frac{\alpha_{P_1}\frac{\beta_{M_1}G_1k_{m_{M_1}}^{-1}}{1+G_1k_{m_{M_1}}^{-1}+\gamma_{M_2}+\gamma_m}}{1+\frac{\beta_{M_1}G_1k_{m_{M_1}}^{-1}}{1+G_1k_{m_{M_1}}^{-1}+\gamma_{M_2}+\gamma_m}+\frac{\beta_{M_2}\gamma_{M_2}}{1+G_1k_{m_{M_1}}^{-1}+\gamma_{M_2}+\gamma_m\left(1+\theta_{M_2}\right)}}$$

$$P_2^* = \frac{\alpha_{P_2}\frac{\beta_{M_2}\gamma_{M_2}}{1+G_1k_{m_{M_1}}^{-1}+\gamma_{M_2}+\gamma_m\left(1+\theta_{M_2}\right)}}{1+\frac{\beta_{M_1}G_1k_{m_{M_1}}^{-1}}{1+G_1k_{m_{M_1}}^{-1}+\gamma_{M_2}+\gamma_m}+\frac{\beta_{M_2}\gamma_{M_2}}{1+G_1k_{m_{M_1}}^{-1}+\gamma_{M_2}+\gamma_m\left(1+\theta_{M_2}\right)}}$$

For fitting, the expressions were normalized the same way as introduced in Supplementary Note 1 and 2 which yields the expressions

$$\hat{P}_1^* = \frac{G_1\left(1+G_{1_0}k_{m_{M_1}}^{-1}+\gamma_{M_2}+\gamma_m\right)}{G_{1_0}\left(1+G_1k_{m_{M_1}}^{-1}+\gamma_{M_2}+\gamma_m\right)}\frac{1+\frac{\beta_{M_1}G_{1_0}k_{m_{M_1}}^{-1}}{1+G_{1_0}k_{m_{M_1}}^{-1}+\gamma_{M_2}+\gamma_m}+\frac{\beta_{M_2}\gamma_{M_2}}{1+G_{1_0}k_{m_{M_1}}^{-1}+\gamma_{M_2}+\gamma_m\left(1+\theta_{M_2}\right)}}{1+\frac{\beta_{M_1}G_1k_{m_{M_1}}^{-1}}{1+G_1k_{m_{M_1}}^{-1}+\gamma_{M_2}+\gamma_m}+\frac{\beta_{M_2}\gamma_{M_2}}{1+G_1k_{m_{M_1}}^{-1}+\gamma_{M_2}+\gamma_m\left(1+\theta_{M_2}\right)}}$$

$$\hat{P}_2^* = \frac{1+G_{1_0}k_{m_{M_1}}^{-1}+\gamma_{M_2}+\gamma_m\left(1+\theta_{M_2}\right)}{1+G_1k_{m_{M_1}}^{-1}+\gamma_{M_2}+\gamma_m\left(1+\theta_{M_2}\right)}\frac{1+\frac{\beta_{M_1}G_{1_0}k_{m_{M_1}}^{-1}}{1+G_{1_0}k_{m_{M_1}}^{-1}+\gamma_{M_2}+\gamma_m}+\frac{\beta_{M_2}\gamma_{M_2}}{1+G_{1_0}k_{m_{M_1}}^{-1}+\gamma_{M_2}+\gamma_m\left(1+\theta_{M_2}\right)}}{1+\frac{\beta_{M_1}G_1k_{m_{M_1}}^{-1}}{1+G_1k_{m_{M_1}}^{-1}+\gamma_{M_2}+\gamma_m}+\frac{\beta_{M_2}\gamma_{M_2}}{1+G_1k_{m_{M_1}}^{-1}+\gamma_{M_2}+\gamma_m\left(1+\theta_{M_2}\right)}}$$

**Synthetic microRNA-based iFFL:**

The system of equations used to obtain the steady state expressions is given by

$$\dot{M}_1 = k_{M_1}^{eff}(G_1, G_2)G_1 - \delta_{M_1} M_1$$

$$\dot{M}_2 = k_{M_2}^{eff}(G_1, G_2)G_2 - \left(\delta_{M_2} + \eta_{M_2} m\right) M_2$$

$$\dot{M}_3 = k_{M_3}^{eff}(G_1, G_2)G_2 - \left(\delta_{M_3} + \eta_{M_3} m\right) M_3$$

$$\dot{m} = k_{m}^{eff}(G_1, G_2)G_2 - \delta_m m$$

$$\dot{P}_1 = k_{P_1}^{eff}(M_1, M_2, M_3)M_1 - \delta_{P_1} P_1$$

$$\dot{P}_2 = k_{P_2}^{eff}(M_1, M_2, M_3)M_2 - \delta_{P_2} P_2$$

$$\dot{P}_1 = k_{P_3}^{eff}(M_1, M_2, M_3)M_3 - \delta_{P_3} P_3$$

Here, $M_1$, $M_2$ and $M_3$ denote the mRNA species of the fluorescent proteins miRFP670 (*X-tra*), mCitrine (*GOI$_1$*) and mRuby3 (*GOI$_2$*) respectively. Similarly, $P_1$, $P_2$ and $P_3$ denote their protein species and $m$ represents the microRNA FF4. $G_1$ corresponds to the plasmid which encodes miRFP670 and $G_2$ corresponds to the plasmid which encodes both the transcriptional unit of mCitrine and the transcriptional unit of mRuby3 and the microRNA. Again, rates beginning with $\delta$ describe degradation rates, while rates beginning with $\eta$ denote the repression by the microRNA. Compared to the endogenous system, the microRNA FF4 is produced from the same gene as mRuby3 and therefore we model their production rates in a similar manner. The steady states for the protein species can be obtained to be

$$P_1^* = \frac{\alpha_{P_1} \frac{\beta_{M_1} G_1 k_{m_{M_1}}^{-1}}{1+G_1 k_{m_{M_1}}^{-1} + \gamma_{M_2} + \gamma_{M_3}}}{1 + \frac{\beta_{M_1} G_1 k_{m_{M_1}}^{-1}}{1+G_1 k_{m_{M_1}}^{-1} + \gamma_{M_2} + \gamma_{M_3}} + \frac{\beta_{M_2}\gamma_{M_2}}{1+G_1 k_{m_{M_1}}^{-1} + \gamma_{M_2} + \gamma_{M_3}\left(1+\theta_{M_2}\right)} + \frac{\beta_{M_3}\gamma_{M_3}}{1+G_1 k_{m_{M_1}}^{-1} + \gamma_{M_2} + \gamma_{M_3}\left(1+\theta_{M_3}\right)}}$$

$$P_2^* = \frac{\alpha_{P_2} \frac{\beta_{M_2}\gamma_{M_2}}{1+G_1 k_{m_{M_1}}^{-1} + \gamma_{M_2} + \gamma_{M_3}\left(1+\theta_{M_2}\right)}}{1 + \frac{\beta_{M_1} G_1 k_{m_{M_1}}^{-1}}{1+G_1 k_{m_{M_1}}^{-1} + \gamma_{M_2} + \gamma_{M_3}} + \frac{\beta_{M_2}\gamma_{M_2}}{1+G_1 k_{m_{M_1}}^{-1} + \gamma_{M_2} + \gamma_{M_3}\left(1+\theta_{M_2}\right)} + \frac{\beta_{M_3}\gamma_{M_3}}{1+G_1 k_{m_{M_1}}^{-1} + \gamma_{M_2} + \gamma_{M_3}\left(1+\theta_{M_3}\right)}}$$

$$P_3^* = \frac{\alpha_{P_3} \frac{\beta_{M_3}\gamma_{M_3}}{1+G_1 k_{m_{M_1}}^{-1} + \gamma_{M_2} + \gamma_{M_3}\left(1+\theta_{M_3}\right)}}{1 + \frac{\beta_{M_1} G_1 k_{m_{M_1}}^{-1}}{1+G_1 k_{m_{M_1}}^{-1} + \gamma_{M_2} + \gamma_{M_3}} + \frac{\beta_{M_2}\gamma_{M_2}}{1+G_1 k_{m_{M_1}}^{-1} + \gamma_{M_2} + \gamma_{M_3}\left(1+\theta_{M_2}\right)} + \frac{\beta_{M_3}\gamma_{M_3}}{1+G_1 k_{m_{M_1}}^{-1} + \gamma_{M_2} + \gamma_{M_3}\left(1+\theta_{M_3}\right)}}$$

For fitting we again use the expression normalized to the first titration of miRFP670.

$$\hat{P}_1^* = \frac{G_1\left(1 + G_{1_0} k_{m_{M_1}}^{-1} + \gamma_{M_2} + \gamma_{M_3}\right)}{G_{1_0}\left(1 + G_1 k_{m_{M_1}}^{-1} + \gamma_{M_2} + \gamma_{M_3}\right)} \frac{1 + \frac{\beta_{M_1} G_{1_0} k_{m_{M_1}}^{-1}}{1+G_{1_0} k_{m_{M_1}}^{-1} + \gamma_{M_2} + \gamma_{M_3}} + \frac{\beta_{M_2}\gamma_{M_2}}{1+G_{1_0} k_{m_{M_1}}^{-1} + \gamma_{M_2} + \gamma_{M_3}\left(1+\theta_{M_2}\right)} + \frac{\beta_{M_3}\gamma_{M_3}}{1+G_{1_0} k_{m_{M_1}}^{-1} + \gamma_{M_2} + \gamma_{M_3}\left(1+\theta_{M_3}\right)}}{1 + \frac{\beta_{M_1} G_1 k_{m_{M_1}}^{-1}}{1+G_1 k_{m_{M_1}}^{-1} + \gamma_{M_2} + \gamma_{M_3}} + \frac{\beta_{M_2}\gamma_{M_2}}{1+G_1 k_{m_{M_1}}^{-1} + \gamma_{M_2} + \gamma_{M_3}\left(1+\theta_{M_2}\right)} + \frac{\beta_{M_3}\gamma_{M_3}}{1+G_1 k_{m_{M_1}}^{-1} + \gamma_{M_2} + \gamma_{M_3}\left(1+\theta_{M_3}\right)}}$$

$$\hat{P}_2^* = \frac{1 + G_{1_0} k_{m_{M_1}}^{-1} + \gamma_{M_2} + \gamma_{M_3}\left(1+\theta_{M_2}\right)}{1 + G_1 k_{m_{M_1}}^{-1} + \gamma_{M_2} + \gamma_{M_3}\left(1+\theta_{M_2}\right)} \frac{1 + \frac{\beta_{M_1} G_{1_0} k_{m_{M_1}}^{-1}}{1+G_{1_0} k_{m_{M_1}}^{-1}+\gamma_{M_2}+\gamma_{M_3}} + \frac{\beta_{M_2}\gamma_{M_2}}{1+G_{1_0} k_{m_{M_1}}^{-1}+\gamma_{M_2}+\gamma_{M_3}\left(1+\theta_{M_2}\right)} + \frac{\beta_{M_3}\gamma_{M_3}}{1+G_{1_0} k_{m_{M_1}}^{-1}+\gamma_{M_2}+\gamma_{M_3}\left(1+\theta_{M_3}\right)}}{1 + \frac{\beta_{M_1} G_1 k_{m_{M_1}}^{-1}}{1+G_1 k_{m_{M_1}}^{-1}+\gamma_{M_2}+\gamma_{M_3}} + \frac{\beta_{M_2}\gamma_{M_2}}{1+G_1 k_{m_{M_1}}^{-1}+\gamma_{M_2}+\gamma_{M_3}\left(1+\theta_{M_2}\right)} + \frac{\beta_{M_3}\gamma_{M_3}}{1+G_1 k_{m_{M_1}}^{-1}+\gamma_{M_2}+\gamma_{M_3}\left(1+\theta_{M_3}\right)}}$$

$$\hat{P}_1^* = \frac{1 + G_{1_0} k_{m_{M_1}}^{-1} + \gamma_{M_2} + \gamma_{M_3}\left(1+\theta_{M_3}\right)}{1 + G_1 k_{m_{M_1}}^{-1} + \gamma_{M_2} + \gamma_{M_3}\left(1+\theta_{M_3}\right)} \frac{1 + \frac{\beta_{M_1} G_{1_0} k_{m_{M_1}}^{-1}}{1+G_{1_0} k_{m_{M_1}}^{-1}+\gamma_{M_2}+\gamma_{M_3}} + \frac{\beta_{M_2}\gamma_{M_2}}{1+G_{1_0} k_{m_{M_1}}^{-1}+\gamma_{M_2}+\gamma_{M_3}\left(1+\theta_{M_2}\right)} + \frac{\beta_{M_3}\gamma_{M_3}}{1+G_{1_0} k_{m_{M_1}}^{-1}+\gamma_{M_2}+\gamma_{M_3}\left(1+\theta_{M_3}\right)}}{1 + \frac{\beta_{M_1} G_1 k_{m_{M_1}}^{-1}}{1+G_1 k_{m_{M_1}}^{-1}+\gamma_{M_2}+\gamma_{M_3}} + \frac{\beta_{M_2}\gamma_{M_2}}{1+G_1 k_{m_{M_1}}^{-1}+\gamma_{M_2}+\gamma_{M_3}\left(1+\theta_{M_2}\right)} + \frac{\beta_{M_3}\gamma_{M_3}}{1+G_1 k_{m_{M_1}}^{-1}+\gamma_{M_2}+\gamma_{M_3}\left(1+\theta_{M_3}\right)}}$$

**Supplementary Table 1.** Transfection tables for all experiments in this study.

**Figure 2a**

| 500 ng total | pGLM49 | pTTF72 | pGLM171 |
|---|---|---|---|
| 1:1 | 62.5 ng | 62.5 ng | 375 ng |
| 1:2 | 62.5 ng | 125 ng | 312.5 ng |
| 1:3 | 62.5 ng | 187.5 ng | 250 ng |
| 1:4 | 62.5 ng | 250 ng | 187.5 ng |
| 2:1 | 125 ng | 62.5 ng | 312.5 ng |
| 2:2 | 125 ng | 125 ng | 250 ng |
| 2:3 | 125 ng | 187.5 ng | 187.5 ng |
| 2:4 | 125 ng | 250 ng | 125 ng |
| 3:1 | 187.5 ng | 62.5 ng | 250 ng |
| 3:2 | 187.5 ng | 125 ng | 187.5 ng |
| 3:3 | 187.5 ng | 187.5 ng | 125 ng |
| 3:4 | 187.5 ng | 250 ng | 62.5 ng |
| 4:1 | 250 ng | 62.5 ng | 187.5 ng |
| 4:2 | 250 ng | 125 ng | 125 ng |
| 4:3 | 250 ng | 187.5 ng | 62.5 ng |
| 4:4 | 250 ng | 250 ng | 0 ng |
| Reagent/cells | | | |
| Optimem | to 50 µL | | |
| PEI | 1.5 µL | | |
| HEK293T | 62500 | | |

| 50 ng total | pGLM49 | pTTF72 | pGLM171 |
|---|---|---|---|
| 1:1 | 6.25 ng | 6.25 ng | 487.5 ng |
| 1:2 | 6.25 ng | 12.5 ng | 481.25 ng |
| 1:3 | 6.25 ng | 18.75 ng | 475 ng |
| 1:4 | 62.5 ng | 25 ng | 468.75 ng |
| 2:1 | 12.5 ng | 6.25 ng | 481.25 ng |
| 2:2 | 12.5 ng | 12.5 ng | 475 ng |
| 2:3 | 12.5 ng | 18.75 ng | 468.75 ng |
| 2:4 | 12.5 ng | 25 ng | 462.5 ng |

| | | | |
|---|---|---|---|
| 3:1 | 18.75 ng | 6.25 ng | 475 ng |
| 3:2 | 18.75 ng | 12.5 ng | 468.75 ng |
| 3:3 | 18.75 ng | 18.75 ng | 462.5 ng |
| 3:4 | 18.75 ng | 25 ng | 456.25 ng |
| 4:1 | 25 ng | 6.25 ng | 468.75 ng |
| 4:2 | 25 ng | 12.5 ng | 462.5 ng |
| 4:3 | 25 ng | 18.75 ng | 456.25 ng |
| 4:4 | 25 ng | 25 ng | 450 ng |
| Reagent/cells | | | |
| Optimem | to 50 µL | | |
| PEI | 1.5 µL | | |
| HEK293T | 62500 | | |

**Supplementary figure 6**

| | pTTF57 | pTTF181 | pGLM49 | pTTF72 | pGLM171 |
|---|---|---|---|---|---|
| EFS/EFS | 174.9 ng | 174.8 ng | 0 ng | 0 ng | 96.8 ng |
| EF1a/EFS | 0 ng | 174.8 ng | 215.0 ng | 0 ng | 56.7 ng |
| EFS/EF1a | 174.9 ng | 0 ng | 0 ng | 215.5 ng | 56.0 ng |
| EF1a/EF1a | 0 ng | 0 ng | 215.0 ng | 215.5 ng | 15.9 ng |
| Reagent/cells | | | | | |
| Optimem | to 50 µL | | | | |
| PEI | 1.5 µL | | | | |
| HEK293T | 50000/75000 | | | | |

**Figure 2b**

| | pGLM49 | pTTF72 | pGLM171 |
|---|---|---|---|
| 1:1 | 62.5 ng | 62.5 ng | 375 ng |
| Reagent/cells | | | |
| Optimem | to 50 µL | | |
| PEI | 1.5 µL | | |
| HEK293T | 62500 | | |

**Supplementary figure 7**

| | pTTF194 | pTTF72 | pGLM171 |
|---|---|---|---|
| 2:4 | 112.7 ng | 156.1 ng | 229.2 ng |
| 4:4 | 225.3 ng | 156.1 ng | 114.6 ng |
| 6:4 | 338 ng | 156.1 ng | 0 ng |
| Reagent/cells | | | |
| Optimem | to 50 µL | | |
| PEI | 1.5 µL | | |
| HEK293T | 50000/75000 | | |

**Figure 2c and supplementary figure 1, 2, 3, 4, 8, 14a**

| | pL-A1 | pBI-G | pEMPTY |
|---|---|---|---|
| 1.0 | 120 ng | 120 ng | 360 ng |
| 1.5 | 120 ng | 180 ng | 200 ng |
| 2.0 | 120 ng | 240 ng | 140 ng |
| 2.5 | 120 ng | 300 ng | 80 ng |
| Reagent/cells | | | |
| Optimem | 50 µL | | |
| Lipofectemine 3000 | 0.75 µL | | |
| P3000 | 1 µL | | |
| H1299 | 150000 | | |
| U2OS | 200000 | | |
| HeLa | 200000 | | |
| HEK293T | 150000 | | |
| CHO-K1 | 150000 | | |

**Supplementary figure 5**

| | ai274 | pBI-G | pEMPTY |
|---|---|---|---|
| 1.0 | 120 ng | 120 ng | 360 ng |
| 1.5 | 120 ng | 180 ng | 200 ng |
| 2.0 | 120 ng | 240 ng | 140 ng |
| 2.5 | 120 ng | 300 ng | 80 ng |
| Reagent/cells | | | |
| Optimem | 50 µL | | |

| | |
|---|---|
| Lipofectemine 3000 | 0.75 µL |
| P3000 | 1 µL |
| H1299 | 150000 |
| HEK | 150000 |

**Figure 2d and supplementary figure 9, 24**

| | pBI-F3G | pBI-H3G |
|---|---|---|
| noTS | 500 ng | |
| miR31 iFFL | | 500 ng |
| Reagent/cells | | |
| Optimem | 250 µL | |
| Lipofectemine 3000 | 11 µL | |
| P3000 | 8.25 µL | |
| H1299 | 1650000 | |

**Figure 2e and supplementary figure 10**

| | pTTF218 | pTTF219 |
|---|---|---|
| HDV (-) | 0 ng | 500 ng |
| HDV (+) | 500 ng | 0 ng |
| Reagent/cells | | |
| Optimem | to 50 µL | |
| PEI | 1.5 µL | |
| HEK293T | 70000 | |

**Figure 2f and supplementary figure 11, 14b**

| | pL-A1 | Oipron | pBI-G | pEMPTY |
|---|---|---|---|---|
| (-) synthetic intron | 50 ng | | 50 ng | 200 ng |
| (+) synthtic intron | | 50 ng | 50 ng | 200 ng |
| Reagent/cells | | | | |
| Optimem | 50 µL | | | |
| Lipofectemine 3000 | 0.75 µL | | | |
| P3000 | 1 µL | | | |
| H1299 | 150000 | | | |
| HEK293T | 150000 | | | |

| | |
|---|---|
| U2OS | 200000 |
| HeLa | 200000 |
| CHO-K1 | 150000 |

**Figure 2g and supplementary figure 12, 13, 14c**

| | pL-S1 | pL-C1 | p125 | pL-R1 | pL-A1 | pEMPTY |
|---|---|---|---|---|---|---|
| L7Ae control | 50 ng | | | | 50 ng | 200 ng |
| L7Ae | 50 ng | | 50 ng | | 50 ng | 150 ng |
| Ms2-cNOT7 control | | 50 ng | | | 50 ng | 200 ng |
| Ms2-cNOT7 | | 50 ng | | 50 ng | 50 ng | 150 ng |
| Reagent/cells | | | | | | |
| Optimem | 50 µL | | | | | |
| Lipofectemine 3000 | 0.75 µL | | | | | |
| P3000 | 1 µL | | | | | |
| H1299 | 150000 | | | | | |
| HEK | 150000 | | | | | |
| U2OS | 200000 | | | | | |
| HeLa | 200000 | | | | | |
| CHO-K1 | 150000 | | | | | |

**Figure 2h**

| | pL-A1 | pH-7 | pH-22 | pH-14 | pL-S1 | pEMPTY |
|---|---|---|---|---|---|---|
| H1299 control | 50 ng | | | | 50 ng | 200 ng |
| H1299 reppressed | | 50 ng | | | 50 ng | 200 ng |
| U2OS control | 50 ng | | | | 50 ng | 200 ng |
| U2OS repressed | | | 50 ng | | 50 ng | 200 ng |
| HeLa control | 50 ng | | | | 50 ng | 200 ng |
| HeLa repressed | | | | 50 ng | 50 ng | 200 ng |
| Reagent/cells | | | | | | |
| Optimem | 50 µL | | | | | |
| Lipofectemine 2000 | 1.5 µL | | | | | |
| P3000 | 1 µL | | | | | |
| H1299 | 150000 | | | | | |
| HeLa | 200000 | | | | | |

| | |
|---|---|
| U2-OS | 200000 |

**Supplementary figure 14d**

| | pL-A1 | pH-10 | pL-S1 | pEMPTY |
|---|---|---|---|---|
| noTS | 50 ng | | 50 ng | 200 ng |
| miR-21 TS | | 50 ng | 50 ng | 200 ng |
| Reagent/cells | | | | |
| Optimem | 50 µL | | | |
| Lipofectemine 3000 | 0.75 µL | | | |
| P3000 | 1 µL | | | |
| CHO-K1 | 150000 | | | |

**Figure 3b,c and supplementary figure 15**

| | pBI-F3G | pBI-H1G/pBI-H3G/pBI-H5G/pBI-H7G | pEMPTY |
|---|---|---|---|
| Control | 100 ng | | 200 ng |
| miR-31 TS | | 100 ng | 200 ng |
| miR-31 TS + inhibitor (20pmol) | | 100 ng | 200 ng |
| Reagent/cells | | | |
| Optimem | 50 µL | | |
| Lipofectemine 3000 | 0.75 µL | | |
| P3000 | 1 µL | | |
| H1299 | 150000 | | |

**Supplementary figure 16**

| | pL-A1 | pH-1/pH-2/pH-3/pH-5/pH-6/pH-7 | pL-S1 | pEMPTY |
|---|---|---|---|---|
| noTS | 50 ng | | 50 ng | 200 ng |
| miR-31 TS | | 50 ng | 50 ng | 200 ng |
| Reagent/cells | | | | |
| Optimem | 50 µL | | | |
| Lipofectemine 3000 | 0.75 µL | | | |
| P3000 | 1 µL | | | |
| H1299 | 150000 | | | |

**Supplementary figure 17**

| | pL-A1 | pH-31 | pL-S1 | pEMPTY |
|---|---|---|---|---|
| noTS | 50 ng | | 50 ng | 200 ng |
| miR-31 TS | | 50 ng | 50 ng | 200 ng |
| Reagent/cells | | | | |
| Optimem | 50 µL | | | |
| Lipofectemine 3000 | 0.75 µL | | | |
| P3000 | 1 µL | | | |
| HEK293T | 150000 | | | |
| U2-OS | 200000 | | | |

**Supplementary figure 18, 19**

| **SF 16** | pL-A1 | pH-16/pH-17/pH-18/pH-20/pH-21/pH-22 | pL-S1 | pEMPTY |
|---|---|---|---|---|
| noTS | 50 ng | | 50 ng | 200 ng |
| miR-221 TS | | 50 ng | 50 ng | 200 ng |
| **SF 17** | pL-A1 | pH-8/pH-9/pH-10/pH-12/pH-13/pH-14 | pL-S1 | pEMPTY |
| noTS | 50 ng | | 50 ng | 200 ng |
| miR-21 TS | | 50 ng | 50 ng | 200 ng |
| Reagent/cells | | | | |
| Optimem | 50 µL | | | |
| Lipofectemine 3000 | 0.75 µL | | | |
| HeLa | 200000 | | | |
| U2-OS | 200000 | | | |

**Figure 4**

| | pGLM49 | pGLM91 | pGLM92 | pGLM102 | pGLM103 | pGLM171 |
|---|---|---|---|---|---|---|
| OLP 0 ng | 140 ng | 0 ng | 180 ng | 0 ng | 0 ng | 180 ng |
| OLP 20 ng | 140 ng | 0 ng | 180 ng | 0 ng | 20 ng | 140 ng |
| OLP 60 ng | 140 ng | 0 ng | 180 ng | 0 ng | 60 ng | 100 ng |
| OLP 100 ng | 140 ng | 0 ng | 180 ng | 0 ng | 100 ng | 60 ng |
| OLP 140 ng | 140 ng | 0 ng | 180 ng | 0 ng | 140 ng | 20 ng |
| OLP 180 ng | 140 ng | 0 ng | 180 ng | 0 ng | 180 ng | 0 ng |
| IFF 0 ng | 140 ng | 180 ng | 0 ng | 0 ng | 0 ng | 180 ng |

| | | | | | | |
|---|---|---|---|---|---|---|
| IFF 20 ng | 140 ng | 180 ng | 0 ng | 0 ng | 20 ng | 140 ng |
| IFF 60 ng | 140 ng | 180 ng | 0 ng | 0 ng | 60 ng | 100 ng |
| IFF 100 ng | 140 ng | 180 ng | 0 ng | 0 ng | 100 ng | 60 ng |
| IFF 140 ng | 140 ng | 180 ng | 0 ng | 0 ng | 140 ng | 20 ng |
| IFF 180 ng | 140 ng | 180 ng | 0 ng | 0 ng | 180 ng | 0 ng |
| FBK 0 ng | 140 ng | 0 ng | 180 ng | 0 ng | 0 ng | 180 ng |
| FBK 20 ng | 140 ng | 0 ng | 180 ng | 20 ng | 0 ng | 140 ng |
| FBK 60 ng | 140 ng | 0 ng | 180 ng | 60 ng | 0 ng | 100 ng |
| FBK 100 ng | 140 ng | 0 ng | 180 ng | 100 ng | 0 ng | 60 ng |
| FBK 140 ng | 140 ng | 0 ng | 180 ng | 140 ng | 0 ng | 20 ng |
| FBK 180 ng | 140 ng | 0 ng | 180 ng | 180 ng | 0 ng | 0 ng |
| HYB 0 ng | 140 ng | 180 ng | 0 ng | 0 ng | 0 ng | 180 ng |
| HYB 20 ng | 140 ng | 180 ng | 0 ng | 20 ng | 0 ng | 140 ng |
| HYB 60 ng | 140 ng | 180 ng | 0 ng | 60 ng | 0 ng | 100 ng |
| HYB 100 ng | 140 ng | 180 ng | 0 ng | 100 ng | 0 ng | 60 ng |
| HYB 140 ng | 140 ng | 180 ng | 0 ng | 140 ng | 0 ng | 20 ng |
| HYB 180 ng | 140 ng | 180 ng | 0 ng | 180 ng | 0 ng | 0 ng |
| Reagent/cells | | | | | | |
| Optimem | to 50 µL | | | | | |
| PEI | 1.5 µL | | | | | |
| HEK293T | 62500 | | | | | |

**Figure 5c and supplementary figure 20, 21, 23**

| | pL-A1 | p-H3/p-H18 | pBI-G | pEMPTY |
|---|---|---|---|---|
| 0.25 noTS | 120 ng | | 30 ng | 150 ng |
| 0.25 miR-TS | | 120 ng | 30 ng | 150 ng |
| 0.50 noTS | 120 ng | | 60 ng | 120 ng |
| 0.50 miR-TS | | 120 ng | 60 ng | 120 ng |
| 0.75 noTS | 120 ng | | 90 ng | 90 ng |
| 0.75 miR-TS | | 120 ng | 90 ng | 90 ng |
| 1.00 noTS | 120 ng | | 120 ng | 60 ng |
| 1.00 miR-TS | | 120 ng | 120 ng | 60 ng |
| 1.25 noTS | 120 ng | | 150 ng | 30 ng |
| 1.25 miR-TS | | 120 ng | 150 ng | 30 ng |

| | | | |
|---|---|---|---|
| 1.50 noTS | 120 ng | | 180 ng |
| 1.50 miR-TS | | 120 ng | 180 ng |
| Reagent/cells | | | |
| Optimem | 50 µL | | |
| Lipofectemine 3000 | 0.75 µL | | |
| P3000 | 1 µL | | |
| H1299 | 150000 | | |
| U2OS | 200000 | | |
| HEK293T | 150000 | | |

**Supplementary figure 22**

| | pL-A1 | pH-18 | pL-S1 | pEMPTY |
|---|---|---|---|---|
| noTS | 50 ng | | 50 ng | 200 ng |
| miR-221 TS | | 50 ng | 50 ng | 200 ng |
| Reagent/cells | | | | |
| Optimem | 50 µL | | | |
| Lipofctemine 3000 | 0.75 µL | | | |
| P3000 | 1 µL | | | |
| HEK | 150000 | | | |

**Figure 5d**

| | pTTF220 | pTTF223 | pTTF138 | pGLM171 |
|---|---|---|---|---|
| 0.25 mCit.-3xTFF5/mRub.-3xTFF5 | 155.1 ng | 0 ng | 24.2 ng | 120.7 ng |
| 0.50 mCit.-3xTFF5/mRub.-3xTFF5 | 155.1 ng | 0 ng | 48.3 ng | 96.6 ng |
| 0.75 mCit.-3xTFF5/mRub.-3xTFF5 | 155.1 ng | 0 ng | 72.5 ng | 72.4 ng |
| 1.00 mCit.-3xTFF5/mRub.-3xTFF5 | 155.1 ng | 0 ng | 96.6 ng | 48.3 ng |
| 1.25 mCit.-3xTFF5/mRub.-3xTFF5 | 155.1 ng | 0 ng | 120.8 ng | 24.1 ng |
| 1.50 mCit.-3xTFF5/mRub.-3xTFF5 | 155.1 ng | 0 ng | 144.9 ng | 0 ng |
| 0.25 mCit.-3xTFF4/mRub.-3xTFF4 | 0 ng | 155.1 ng | 24.2 ng | 120.7 ng |
| 0.50 mCit.-3xTFF4/mRub.-3xTFF4 | 0 ng | 155.1 ng | 48.3 ng | 96.6 ng |
| 0.75 mCit.-3xTFF4/mRub.-3xTFF4 | 0 ng | 155.1 ng | 72.5 ng | 72.4 ng |
| 1.00 mCit.-3xTFF4/mRub.-3xTFF4 | 0 ng | 155.1 ng | 96.6 ng | 48.3 ng |
| 1.25 mCit.-3xTFF4/mRub.-3xTFF4 | 0 ng | 155.1 ng | 120.8 ng | 24.1 ng |
| 1.50 mCit.-3xTFF4/mRub.-3xTFF4 | 0 ng | 155.1 ng | 144.9 ng | 0 ng |

| Reagent/cells | |
| --- | --- |
| Optimem | to 50 µL |
| Lipofectamine 2000 | 0.6 µL |
| mES E14 | 70000 |

**Supplementary figure 25**

| | pTTF220 | pTTF223 | pTTF138 | pGLM171 |
| --- | --- | --- | --- | --- |
| 0.25 mCit.-3xTFF5/mRub.-3xTFF5 | 258.4 ng | 0 ng | 40.3 ng | 201.3 ng |
| 0.50 mCit.-3xTFF5/mRub.-3xTFF5 | 258.4 ng | 0 ng | 80.5 ng | 161.1 ng |
| 0.75 mCit.-3xTFF5/mRub.-3xTFF5 | 258.4 ng | 0 ng | 120.8 ng | 120.8 ng |
| 1.00 mCit.-3xTFF5/mRub.-3xTFF5 | 258.4 ng | 0 ng | 161.0 ng | 80.6 ng |
| 1.25 mCit.-3xTFF5/mRub.-3xTFF5 | 258.4 ng | 0 ng | 201.3 ng | 40.3 ng |
| 1.50 mCit.-3xTFF5/mRub.-3xTFF5 | 258.4 ng | 0 ng | 241.6 ng | 0 ng |
| 0.25 mCit.-3xTFF4/mRub.-3xTFF4 | 0 ng | 258.4 ng | 40.3 ng | 201.3 ng |
| 0.50 mCit.-3xTFF4/mRub.-3xTFF4 | 0 ng | 258.4 ng | 80.5 ng | 161.1 ng |
| 0.75 mCit.-3xTFF4/mRub.-3xTFF4 | 0 ng | 258.4 ng | 120.8 ng | 120.8 ng |
| 1.00 mCit.-3xTFF4/mRub.-3xTFF4 | 0 ng | 258.4 ng | 161.0 ng | 80.6 ng |
| 1.25 mCit.-3xTFF4/mRub.-3xTFF4 | 0 ng | 258.4 ng | 201.3 ng | 40.3 ng |
| 1.50 mCit.-3xTFF4/mRub.-3xTFF4 | 0 ng | 258.4 ng | 241.6 ng | 0 ng |
| Reagent/cells | | | | |
| Optimem | to 50 µL | | | |
| PEI | 1.5 µL | | | |
| HEK293T | 70000 | | | |

**Supplementary Table 2. List of the plasmids used in this study. Plasmids sequences are available on GenBank.**

| Fig. | Short plasmid name | Full plasmid name | Parts from | GenBank accession code |
|---|---|---|---|---|
| 2c-e-f-g, 5b & SF1, 2, 3, 4, 8, 11, 12, 13, 14, 16, 17, 18, 19, 20, 21, 23 | pL-A1 | pT-GTW6-CMV-mKate | [2] | MH107777 |
| SF5 | ai274 | pT-PGK-mCherry | [3] | |
| 2f & SF11, 14b | Oipron | pT-GTW6-CMV-SImKate | [4,5] | |
| 2c-e, 5b & SF1, 2, 3, 4, 5, 8, 11, 14a-b, 20, 21, 23 | pBI-G | pBI-CMV1_EGFP | Clontech 631630 | |
| 2f-g & SF12, 13, 14c-d, 12, 13, 14c-d, 16, 17, 18, 19, 22 | pL-S1 | pBoxCDGC_2xKMet_EGFP | [6] | |
| 2g & SF13 | pL-C1 | pBoxCDGCmut_KMetEGFP-8xMS2-pA | [6] | MH883358 |
| 2g & SF12, 14c | p125 | pT-GTW6-CMV-L7AescFv35 | [6] | MH883336 |
| 2g & SF13 | pL-R1 | pT-GTW6-CMV-MS2- CNOT7 | [6] | MH883359 |
| SF16 | pH-1 | pT-GTW6-CMV-mKate_1xmiR31TS5' | [5] | |
| SF16 | pH-2 | pT-GTW6-CMV-mKate_2xmiR31TS5' | [5] | |
| 5c & SF16 | pH-3 | pT-GTW6-CMV-mKate_3xmiR31TS5' | [5] | |
| SF16 | pH-5 | pT-GTW6-CMV-mKate_1xmiR31TS3' | [5] | |
| SF16 | pH-6 | pT-GTW6-CMV-mKate_2xmiR31TS3' | [5] | |

| | | | |
|---|---|---|---|
| 2h & SF16 | pH-7 | pT-GTW6-CMV-mKate_3xmiR31TS3' | 5 |
| SF17 | pH-31 | pT-GTW6-CMV-mKate_4xmiR31TS3' | 5 |
| 2d, 3 & SF9, 15, 24 | pBI-F3G | pBI-CMV1_EGFP_mKate | Clontech 631630 |
| 3 & SF15 | pBI-H1G | pBI-CMV1_EGFP_mKate_1xmiR31TS5' | Clontech 631630 |
| 3 & SF9, 15, 24 | pBI-H3G | pBI-CMV1_EGFP_mKate_3xmiR31TS5' | Clontech 631630 |
| 3 & SF15 | pBI-H5G | pBI-CMV1_EGFP_mKate_1xmiR31TS3' | Clontech 631630 |
| 3 | pBI-H7G | pBI-CMV1_EGFP_mKate_3xmiR31TS3' | Clontech 631630 |
| SF18 | pH-8 | pT-GTW6-CMV-mKate_1xmiR21TS5' | 5 |
| SF18 | pH-9 | pT-GTW6-CMV-mKate_2xmiR21TS5' | 5 |
| SF18 | pH-10 | pT-GTW6-CMV-mKate_3xmiR21TS5' | 5 |
| SF18 | pH-12 | pT-GTW6-CMV-mKate_1xmiR21TS3' | 5 |
| SF18 | pH-13 | pT-GTW6-CMV-mKate_2xmiR21TS3' | 5 |
| 2h & SF19 | pH-14 | pT-GTW6-CMV-mKate_3xmiR21TS3' | 5 |
| SF18 | pH-16 | pT-GTW6-CMV-mKate_1xmiR221TS5' | 5 |
| SF18 | pH-17 | pT-GTW6-CMV-mKate_2xmiR221TS5' | 5 |
| SF18, 120, 21, 22, 23 | pH-18 | pT-GTW6-CMV-mKate_3xmiR221TS5' | 5 |

| | | | |
|---|---|---|---|
| SF18 | pH-20 | pT-GTW6-CMV-mKate_1xmiR221 TS3' | [5] |
| SF18 | pH-21 | pT-GTW6-CMV-mKate_2xmiR221 TS3' | [5] |
| 2h & SF18 | pH-22 | pT-GTW6-CMV-mKate_3xmiR221 TS3' | [5] |
| 2a, 4 & SF6 | pGLM49 | INS-bGHpA-$P_{EF1a}$-mCitrine-SV40pA-INS | [1] |
| 2a, SF6 & SF7 | pTT72 | INS-bGHpA-$P_{EF1a}$-mRuby3-SV40pA-INS | |
| 2a, 4, 5e & SF25 | pGLM171 | AmpR-INS-bGHpA-SV40pA-INS-pUCori | |
| 2b | pTTF84 | INS-bGHpA-synpA-$P_{EF1a}$-tTA2::Cerulean-SV40pA-INS-bGHpA-$P_{EF1a}$-mCitrine-SV40pA | |
| 2b | pTTF145 | INS-bGHpA-mRuby3-$P_{biTRE}$-miRFP670-SV40pA-INS | |
| 2e & SF10 | pTTF218 | $P_{TRE}$-(HDV)mCitrine-SV40pA-INS-bGHpA-$P_{EF1a}$-tTA2::P2A::mRuby3-SV40pA-$P_{SV40}$-puΔtk-SV40pA | |
| 2e & SF10 | pTTF219 | $P_{TRE}$-(dHDV)mCitrine-SV40pA-INS-bGHpA-$P_{EF1a}$-tTA2::P2A::mRuby3-SV40pA-$P_{SV40}$-puΔtk-SV40pA | |
| SF6 | pTTF57 | INS-bGHpA-$P_{EFS}$-mCitrine-SV40pA-INS | |
| SF6 | pTTF181 | INS-bGHpA-$P_{EFS}$-mRuby3-SV40pA-INS | |
| 4 | pGLM91 | INS-bGHpA-$P_{TRE}$-DsRed(FF4)-TFF4x3-SV40pA-INS | [1] |
| 4 | pGLM92 | INS-bGHpA-$P_{TRE}$-DsRed(FF4)-TFF5x3-SV40pA-INS | |
| 4b | pGLM103 | $P_{SV40}$-PuroR-SV40pA-INS-bGHpA-$P_{EF1a}$-tTA::Cerulean-TFF5x3-SV40pA-INS | [1] |

| | | |
|---|---|---|
| 4c | pGLM102 | $P_{SV40}$-PuroR-SV40pA-INS-bGHpA-$P_{EF1a}$-tTA::Cerulean-TFF4x3-SV40pA-INS |
| 5e & SF25 | pTTF138 | INS-bGHpA-$P_{EF1a}$-miRFP670-SV40pA-INS |
| 5e & SF25 | pTTF220 | $P_{SV40}$-PuroR-SV40pA-INS-bGHpA-$P_{EF1a}$-mCitrine-TFF5x3-SV40pA-INS-$P_{EF1a}$-mRuby3(FF4)-TFF5x3-SV40pA-INS |
| 5e & SF25 | pTTF223 | $P_{SV40}$-PuroR-SV40pA-INS-bGHpA-$P_{EF1a}$-mCitrine-TFF4x3-SV40pA-INS-$P_{EF1a}$-mRuby3(FF4)-TFF4x3-SV40pA-INS |
| SF7 | pTTF194 | INS-bGHpA-$P_{EF1a}$-mCitrine-SV40pA-INS-$P_{SV40}$-miRFP670-SV40pA-INS |

**Supplementary Table 3. List of the primers and oligos used to generate miRNA target sites.**

| Construct | Primer name | Primer Sequence |
|---|---|---|
| mKate_1xmiR31TS5' | Fw1 | GATCCAGCTATGCCAGCATCTTGCCTG |
| mKate_1xmiR31TS5' | Rv1 | CTAGCAGGCAAGATGCTGGCATAGCTG |
| mKate_2xmiR31TS5' | Fw2 | GATCCAGCTATGCCAGCATCTTGCCTAGCTATGCCAGCATCTTGCCTG |
| mKate_2xmiR31TS5' | Rv2 | CTAGCAGGCAAGATGCTGGCATAGCTAGGCAAGATGCTGGCATAGCTG |
| mKate_3xmiR31TS5' | Fw3 | GATCCAGCTATGCCAGCATCTTGCCTAGCTATGCCAGCATCT |
| mKate_3xmiR31TS5' | Rv3 | TAGCTAGGCAAGATGCTGGCATAGCTG |
| mKate_3xmiR31TS5' | Fw4 | TGCCTAGCTATGCCAGCATCTTGCCTG |
| mKate_3xmiR31TS5' | Rv4 | CTAGCAGGCAAGATGCTGGCATAGCTAGGCAAGATGCTGGCA |
| mKate_1xmiR31TS3' | Fw5 | AGCTTAGCTATGCCAGCATCTTGCCTTTAAT |
| mKate_1xmiR31TS3' | Rv5 | TAAAGGCAAGATGCTGGCATAGCTA |
| mKate_2xmiR31TS3' | Fw6 | AGCTTAGCTATGCCAGCATCTTGCCTAGCTATGCCAGCATCTTGCCTTTAAT |
| mKate_2xmiR31TS3' | Rv6 | TAAAGGCAAGATGCTGGCATAGCTAGGCAAGATGCTGGCATAGCTA |
| mKate_3xmiR31TS3' | Fw7 | AGCTTAGCTATGCCAGCATCTTGCCTAGCTATGCCAGCATCT |
| mKate_3xmiR31TS3' | Rv7 | TAGCTAGGCAAGATGCTGGCATAGCTA |
| mKate_3xmiR31TS3' | Fw8 | TGCCTAGCTATGCCAGCATCTTGCCTTTAAT |
| mKate_3xmiR31TS3' | Rw8 | TAAAGGCAAGATGCTGGCATAGCTAGGCAAGATGCTGGCA |
| mKate_1xmiR21TS5' | Fw9 | GATCCTAGCTTATCAGACTGATGTTGAG |
| mKate_1xmiR21TS5' | Rv9 | CTAGCTCAACATCAGTCTGATAAGCTAG |
| mKate_2xmiR21TS5' | Fw10 | GATCCTAGCTTATCAGACTGATGTTGATAGCTTATCAGACTGATGTTGAG |
| mKate_2xmiR21TS5' | Rv10 | CTAGCTCAACATCAGTCTGATAAGCTATCAACATCAGTCTGATAAGCTAG |
| mKate_3xmiR21TS5' | Fw11 | GATCCTAGCTTATCAGACTGATGTTGATAGCTTATCAGACTGAT |

| | | |
|---|---|---|
| mKate_3xmiR21TS5' | Rv11 | AGCTATCAACATCAGTCTGATAAGCTAG |
| mKate_3xmiR21TS5' | Fw12 | GTTGATAGCTTATCAGACTGATGTTGAG |
| mKate_3xmiR21TS5' | Rv12 | CTAGCTCAACATCAGTCTGATAAGCTATCAACATCAGTCTGATA |
| mKate_1xmiR21TS3' | Fw15 | AGCTTTAGCTTATCAGACTGATGTTGATTAAT |
| mKate_1xmiR21TS3' | Rv15 | TAATCAACATCAGTCTGATAAGCTAA |
| mKate_2xmiR21TS3' | Fw16 | AGCTTTAGCTTATCAGACTGATGTTGATAGCTTATCAGACTGATGTTGATTAAT |
| mKate_2xmiR21TS3' | Rv16 | TAATCAACATCAGTCTGATAAGCTATCAACATCAGTCTGATAAGCTAA |
| mKate_3xmiR21TS3' | Fw17 | AGCTTTAGCTTATCAGACTGATGTTGATAGCTTATCAGACTGAT |
| mKate_3xmiR21TS3' | Rv17 | AGCTATCAACATCAGTCTGATAAGCTAA |
| mKate_3xmiR21TS3' | Fw18 | GTTGATAGCTTATCAGACTGATGTTGATTAAT |
| mKate_3xmiR21TS3' | Rv18 | TAATCAACATCAGTCTGATAAGCTATCAACATCAGTCTGATA |
| mKate_1xmiR221TS5' | Fw21 | GATCCACCTGGCATACAATGTAGATTTG |
| mKate_1xmiR221TS5' | Rv21 | CTAGCAAATCTACATTGTATGCCAGGTG |
| mKate_2xmiR221TS5' | Fw22 | GATCCACCTGGCATACAATGTAGATTTACCTGGCATACAATGTAGATTTG |
| mKate_2xmiR221TS5' | Rv22 | CTAGCAAATCTACATTGTATGCCAGGTAAATCTACATTGTATGCCAGGTG |
| mKate_3xmiR221TS5' | Fw23 | GATCCACCTGGCATACAATGTAGATTTACCTGGCATACAATGT |
| mKate_3xmiR221TS5' | Rv23 | CAGGTAAATCTACATTGTATGCCAGGTG |
| mKate_3xmiR221TS5' | Fw24 | AGATTTACCTGGCATACAATGTAGATTTG |
| mKate_3xmiR221TS5' | Rv24 | CTAGCAAATCTACATTGTATGCCAGGTAAATCTACATTGTATGC |
| mKate_1xmiR221TS3' | Fw25 | AGCTTACCTGGCATACAATGTAGATTTTTAAT |
| mKate_1xmiR221TS3' | Rv25 | TAAAAATCTACATTGTATGCCAGGTA |
| mKate_2xmiR221TS3' | Fw26 | AGCTTACCTGGCATACAATGTAGATTTACCTGGCATACAATGTAGATTTTTAAT |
| mKate_2xmiR221TS3' | Rv26 | TAAAAATCTACATTGTATGCCAGGTAAATCTACATTGTATGCCAGGTA |
| mKate_3xmiR221TS3' | Fw27 | AGCTTACCTGGCATACAATGTAGATTTACCTGGCATACAATGTA |

| | | |
|---|---|---|
| mKate_3xmiR221TS3' | Rv27 | CAGGTAAATCTACATTGTATGCCAGGTA |
| mKate_3xmiR221TS3' | Fw28 | GATTTACCTGGCATACAATGTAGATTTTTAAT |
| mKate_3xmiR221TS3' | Rv28 | TAAAAATCTACATTGTATGCCAGGTAAATCTACATTGTATGC |

**Supplementary Table 4. List of the target sites for miRNAs used in this study [7].**

| miRNA | Sequence |
|---|---|
| hsa-miR-31-5p | AGCTATGCCAGCATCTTGCCT |
| hsa-miR-21 | TCAACATCAGTCTGATAAGCTA |
| hsa-miR-221 | AAATCTACATTGTATGCCAGGT |
| miR-FF4 | CCGCTTGAAGTCTTTAATTAAA |
| miR-FF5 | AAGCACTCTGATTTGACAATTA |

**Supplementary Table 5. List of the primers used for qPCR analyses.**

| Primer | Function | Sequence (5'-3') |
|---|---|---|
| F7 | Forward primer for mKate | GGTGTCTAAGGGCGAAGAGC |
| F8 | Reverse primer for mKate | GCTGGTAGCCAGGATGTCGA |
| qPCR-EGFP-F | Forward primer for EGFP | AAGGGCATCGACTTCAAG |
| qPCR-EGFP-R | Reverse primer for EGFP | TGCTTGTCGGCCATGATATG |
| qPCR-18S-F | Forward primer for 18S | GCTTAATTTGACTCAACACGGGA |
| qPCR-18S-R | Reverse primer for 18S | AGCTATCAATCTGTCAATCCTGTC |

**Supplementary Table 6.** 5x isothermal reaction buffer recipe.

| Component | Concentration |
|---|---|
| PEG-800 | 25 % |
| Tris-HCl, pH 7.5 | 500 mM |
| $MgCl_2$ | 50 mM |
| DTT | 50 mM |
| dATP | 1 mM |
| dTTP | 1 mM |
| dCTP | 1 mM |
| dGTP | 1 mM |
| NAD | 5 mM |
| To 3 mL with dd$H_2O$ | |

**Supplementary Table 7.** Parameter fits related to Figure 4B/C and Supplementary Note 3.

| **Parameter** | **Unit** | **Value** |
|---|---|---|
| $\alpha_{P_1}$ | $Arbitrary fluorescence$ | 3108.52 |
| $\alpha_{P_2}$ | $Arbitrary fluorescence$ | 28007.3 |
| $\alpha_{P_3}$ | $Arbitrary fluorescence$ | 2198.66 |
| $\beta_{M_1}$ | $Unitless$ | 9275.79 |
| $\beta_{M_2}$ | $Unitless$ | 1017.34 |
| $\beta_{M_3}$ | $Unitless$ | 709.432 |
| $k_{m_{M_1}}$ | $ng$ | 0.00400709 |
| $\gamma_{M_2}$ | $Unitless$ | 1.35779 |
| $\gamma_{M_3}$ | $Unitless$ | 14.5078 |
| $\theta_{M_1}$ | $Unitless$ | 286.573 |
| $\theta_{M_2}$ | $Unitless$ | 126.876 |
| $\kappa$ | $Arbitrary fluorescence$ | 17.2643 |
| $h$ | $Unitless$ | 31 |

**Supplementary Table 8.** Parameter fits related to Figure 5c and Supplementary Note 4 endogenous microRNA-based iFFL.

| **Parameter** | **Unit** | **Value** |
|---|---|---|
| $\beta_{M_1}$ | $Unitless$ | 0.746056 |
| $\beta_{M_2}$ | $Unitless$ | 32.429 |
| $k_{m_{M_1}}$ | $Unitless$ | 59.104 |
| $\gamma_{M_2}$ | $Unitless$ | 0.480552 |
| $\gamma_m$ | $Unitless$ | 33.2046 |
| $\theta_{M_2}$ | $Unitless$ | 227.144 |

**Supplementary Table 9.** Parameter fits related to Figure 5d and Supplementary Note 4 synthetic microRNA-based iFFL.

| **Parameter** | **Unit** | **Value** |
|---|---|---|
| $\beta_{M_1}$ | $Unitless$ | 1.89289 |
| $\beta_{M_2}$ | $Unitless$ | 0.0152265 |
| $\beta_{M_3}$ | $Unitless$ | 1.43347e-18 |
| $k_{m_{M_1}}$ | $Unitless$ | 3.21455 |
| $\gamma_{M_2}$ | $Unitless$ | 2.04539e-16 |
| $\gamma_{M_3}$ | $Unitless$ | 1.91369 |
| $\theta_{M_2}$ | $Unitless$ | 9.80712 |
| $\theta_{M_3}$ | $Unitless$ | 30.272 |

**Supplementary Table 10.** Parameter fits related to Supplementary Figure 19 and Supplementary Note 4 endogenous microRNA-based iFFL.

| **Parameter** | **Unit** | **Value** |
|---|---|---|
| $\beta_{M_1}$ | $Unitless$ | 1.01232e-9 |
| $\beta_{M_2}$ | $Unitless$ | 0.943493 |
| $k_{m_{M_1}}$ | $Unitless$ | 3.57119 |
| $\gamma_{M_2}$ | $Unitless$ | 2.85075e-11 |
| $\gamma_m$ | $Unitless$ | 0.173874 |
| $\theta_{M_2}$ | $Unitless$ | 10.9904 |

**Supplementary Table 11.** Parameter fits related to Supplementary Figure 21 and Supplementary Note 4 endogenous microRNA-based iFFL.

| **Parameter** | **Unit** | **Value** |
|---|---|---|
| $\beta_{M_1}$ | $Unitless$ | 2.81166 |
| $\beta_{M_2}$ | $Unitless$ | 1.95439e-5 |
| $k_{m_{M_1}}$ | $Unitless$ | 2.32439 |
| $\gamma_{M_2}$ | $Unitless$ | 1.1604e-13 |
| $\gamma_m$ | $Unitless$ | 6.65805 |
| $\theta_{M_2}$ | $Unitless$ | 6.38791 |

**Supplementary Table 12.** Parameter fits related to Supplementary Figure 22 and Supplementary Note 4 synthetic microRNA-based iFFL.

| **Parameter** | **Unit** | **Value** |
|---|---|---|
| $\beta_{M_1}$ | $Unitless$ | 9.61413e-16 |
| $\beta_{M_2}$ | $Unitless$ | 0.00222615 |
| $\beta_{M_3}$ | $Unitless$ | 3.91565e-17 |
| $k_{m_{M_1}}$ | $Unitless$ | 33.6238 |
| $\gamma_{M_2}$ | $Unitless$ | 3.19321e-14 |
| $\gamma_{M_3}$ | $Unitless$ | 13.4858 |
| $\theta_{M_2}$ | $Unitless$ | 6.45491 |
| $\theta_{M_3}$ | $Unitless$ | 30.2392 |
